# spatiAlytica: Viewer-Grounded Multimodal Agentic System for Interactive Spatial Omics Analysis

**DOI:** 10.64898/2026.04.29.721735

**Authors:** Arun Das, Kexun Zhang, Jifeng Song, Meiru Han, Angela Chen, Wen Meng, Hugh Galloway, Po-Yuan Chen, Sumin Jo, Zhentao Liu, Md Musaddaqul Hasib, Adam Officer, Harsh Sinha, Yu-Chiao Chiu, Shou-Jiang Gao, Lei Li, Yufei Huang

## Abstract

Spatial transcriptomics and proteomics map tissue architecture and cellular interactions, but analysis remains limited by programming demands and text-centered AI agents that lack viewer grounding and cross-turn context. We present spatiAlytica, a viewer-centric multimodal interactive agentic system embedded in the Napari viewer that enables non-programmer biologists to perform iterative, hypothesis-driven spatial omics analysis via natural language. spatiAlytica couples viewer-state serialization, agentic memory, biological concept-to-data-field mapping, code generation and debugging, Spatial VQA, and grounded interpretation to support an exploratory analysis and interpretive reasoning workflow. We introduce spatiAlyticaBench, a comprehensive benchmark spanning 222 single-turn spatial analytical coding questions, 178 multi-turn sequential workflow questions, and 7,350 image-grounded reasoning questions. spatiAlytica outperformed strong agentic baselines, while using less time and tokens. Case studies across Kaposi’s sarcoma, colorectal cancer, and ovarian cancer recapitulated known spatial patterns and uncovered progressive CD8 T-cell dysfunction during KS progression.

---

Spatial transcriptomic and proteomic technologies, including image-based in situ transcriptomics [1–4], highly multiplexed imaging [5–8], and spatial barcoding [9–11], enable mapping of cell states and cell-cell interactions across intact tissues [12, 13], revealing revealing immune-evasive spatial niches, therapeutic response, and clinical outcome [14–17]. However, extracting such insights remains technically demanding. Biologists often lack both the coding skills and methodological expertise required, and existing graphical interfaces partly ease this barrier but remain limited to interactive viewing, without support for core analyses such as spatial statistics, niche analysis, or differential expression.

Large language models (LLMs) [18] offer a natural bridge between biological questions and computational tools, and recent agentic systems (AutoBA [19], BioMANIA [20], CellAgent [21], spatialAgent [22], BioMedAgent [23], scChat [24]) demonstrate that LLM agents can automate substantial parts of omics analysis. However, these systems share architectural limitations for interactive spatial biology. Also, general-purpose LLMs generate code that runs but produces biologically invalid results that non-expert users cannot detect [25]. Existing agentic systems are text-centered and not coupled to the spatial viewer, limiting their ability to reason about the current field of view or selected regions. Their memory is typically a flat conversation history, preventing reliable reuse of prior results and disrupting iterative hypothesis testing. Most systems also emphasize execution over interpretation and require users to read code and validate outputs, limiting accessibility for biologists without programming skills [26].

To address these gaps, we developed *spatiAlytica*, a multimodal interactive agentic system for spatial biology analysis. Designed for non-programmer biologists, it automatically resolves biological concepts to dataset columns, accepts natural-language conditions, and returns responses in plain biological language. Embedded in the Napari viewer [27], it treats spatial plots and figures as queryable objects, enabling users to zoom into a region and query the current view. A central orchestrator coordinates specialized subagents for data resolution, code generation, error correction, reporting, and image analysis, while a hybrid tool repository and structured memory sub-system support iterative multi-turn analysis and a unified exploratory analysis-interpretation (EA-IR) workflow.

We also developed *spatiAlyticaBench*, a comprehensive benchmark with three settings: 222 single-turn analytical questions, 178 sequential multi-turn workflow questions, and 7,350 pairs of image-based question-answers over spatial and analytical plots. On this benchmark, spatiAlytica consistently outperformed existing approaches in analysis accuracy, multi-turn reasoning, and viewer-grounded interpretation. Ablation studies show that the agentic memory is essential for robust multi-turn workflows, while the code debugger improves single-turn reliability by recovering first-pass failures. Case studies in Xenium Kaposi sarcoma, CODEX colorectal cancer, and Visium ovarian cancer show that spatiAlytica recapitulates published findings and surfaces a progressive CD8 T-cell dysfunction trajectory during Kaposi’s sarcoma progression that was not described in the dataset’s original single-cell spatial transcriptomic analysis.

## 1 Results

### 1.1 A viewer-centric multimodal interactive agentic architecture for spatial biology

spatiAlytica embeds a multimodal interactive agentic system within the Napari [27] viewer (Fig. 1a; Supplementary Fig. 1a), allowing queries over the underlying spatial dataset saved as an `AnnData` [28] object and the current visual state. The active viewer state, including layers, viewport, legends, color mappings, and user annotations, is serialized as structured JSON and injected into every required sub-agent prompt as a first-class input, so sub-agents reason about the biologist’s current field of view. spatiAlytica automatically infers coordinate columns, categorical labels, and sample identifiers, enabling biologists without programming experience to specify analyses in natural language. Results are returned as linked viewer layers, plots, and biologically framed textual summaries.

**Fig. 1.**
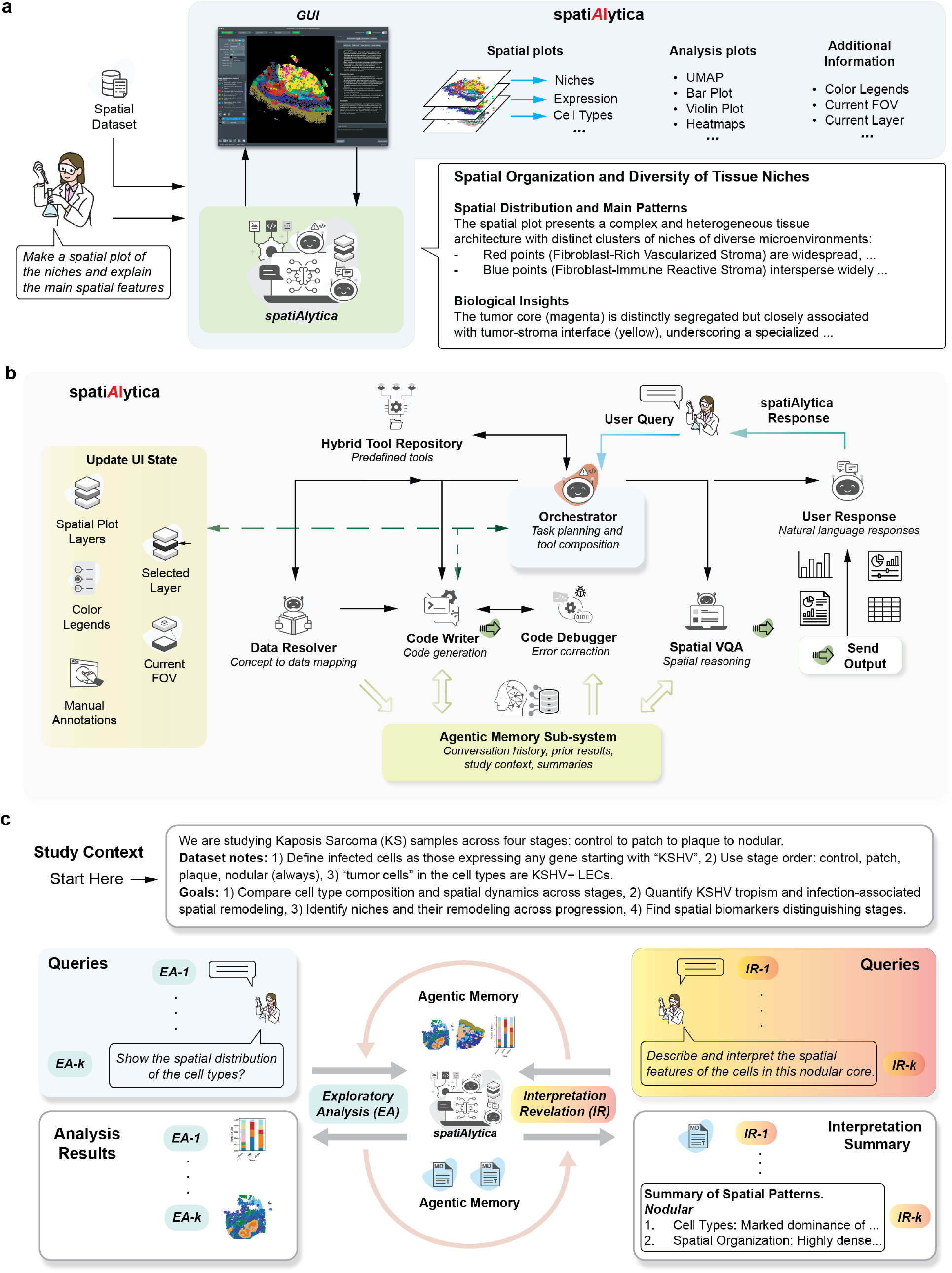
spatiAlytica architecture and interactive workflow. (A) The biologist-centered GUI integrates spatial datasets with a multimodal interactive agentic system. Users load datasets, issue natural-language queries, and receive spatial plots, statistical outputs, and biological interpretations within the Napari viewer. (B) Multimodal interactive agentic architecture. User queries are processed by a central orchestrator that delegates to five specialized sub-agents (data resolver, code writer, code debugger, spatial VQA, and user response). A hybrid tool repository provides domain-specific analytical tools, and the agentic memory sub-system stores conversation context and intermediate results across turns. (C) The exploratory analysis and interpretive reasoning (EA-IR) loop. Users alternate between EA queries that invoke computational tools and generate quantitative outputs, and IR queries that prompt biological synthesis from accumulated results.

These capabilities arise from a modular multimodal interactive agentic architecture and hybrid tool repository (Fig. 1b; Supplementary Fig. 1b; Methods). An **orchestrator** decomposes each request into a plan spanning data exploration, spatial plotting, statistical testing, code execution, and image interpretation. The **data resolver sub-agent** maps biological concepts onto dataset-specific `AnnData` fields through a three-step protocol (fuzzy matching, schema lookup, categorical-value listing; Methods). The **code writer** and **code debugger** sub-agents generate executable code and recover from execution failures. The **spatial VQA sub-agent** consumes the viewer state and rendered image to reason over legends, annotated regions, and the active field of view. The **user response sub-agent** translates tool outputs into plain biological language under anti-hallucination rules grounded in executed tool results. All sub-agents share an **agentic memory sub-system** with three tiers (short-term conversation history, intermediate-results registry tagged with viewport state, and persisted long-term store of named artifacts), supporting anaphoric reuse and multi-hop reasoning across turns. Together, viewer coupling, data-aware analysis, tiered memory, and grounded interpretation distinguish spatiAlytica from prior agentic biology systems (Supplementary Table 1).

These components together enable a unified EA-IR loop (Fig. 1c). Each session begins with a structured study context capturing biological background, dataset mappings, and experimental groups (Supplementary Note 2; Methods), which the orchestrator broadcasts as persistent session-level guidance. EA turns trigger tool-based computation yielding spatial layers, plots, and quantitative summaries; the memory sub-system registers each output together with the active viewport state. IR turns interpret these memory-retained artifacts biologically against the current viewer state. This closed-loop design progressively converts inspectable analytical evidence into biological insight within a single interactive workflow.

### 1.2. spatiAlyticaBench: a comprehensive benchmark for spatial omics conversational analysis

To evaluate spatiAlytica’s interactive, sequentially dependent, and viewer-grounded capabilities, we constructed **spatiAlyticaBench**, a benchmark with three complementary subsets: spatiAlyticaBench-ST for one-shot tool-use correctness, spatiAlyticaBench-MT for state retention under sequential dependencies, and spatiAlyticaBench-ImageQA for grounded reasoning over rendered spatial visualizations, spanning 11 datasets from seven spatial omics platforms and three single-cell RNA-seq references, split roughly between data-returning (250) and plot-saving (150) outputs (Fig. 2; Supplementary Tables 2, 3).

**Fig. 2.**
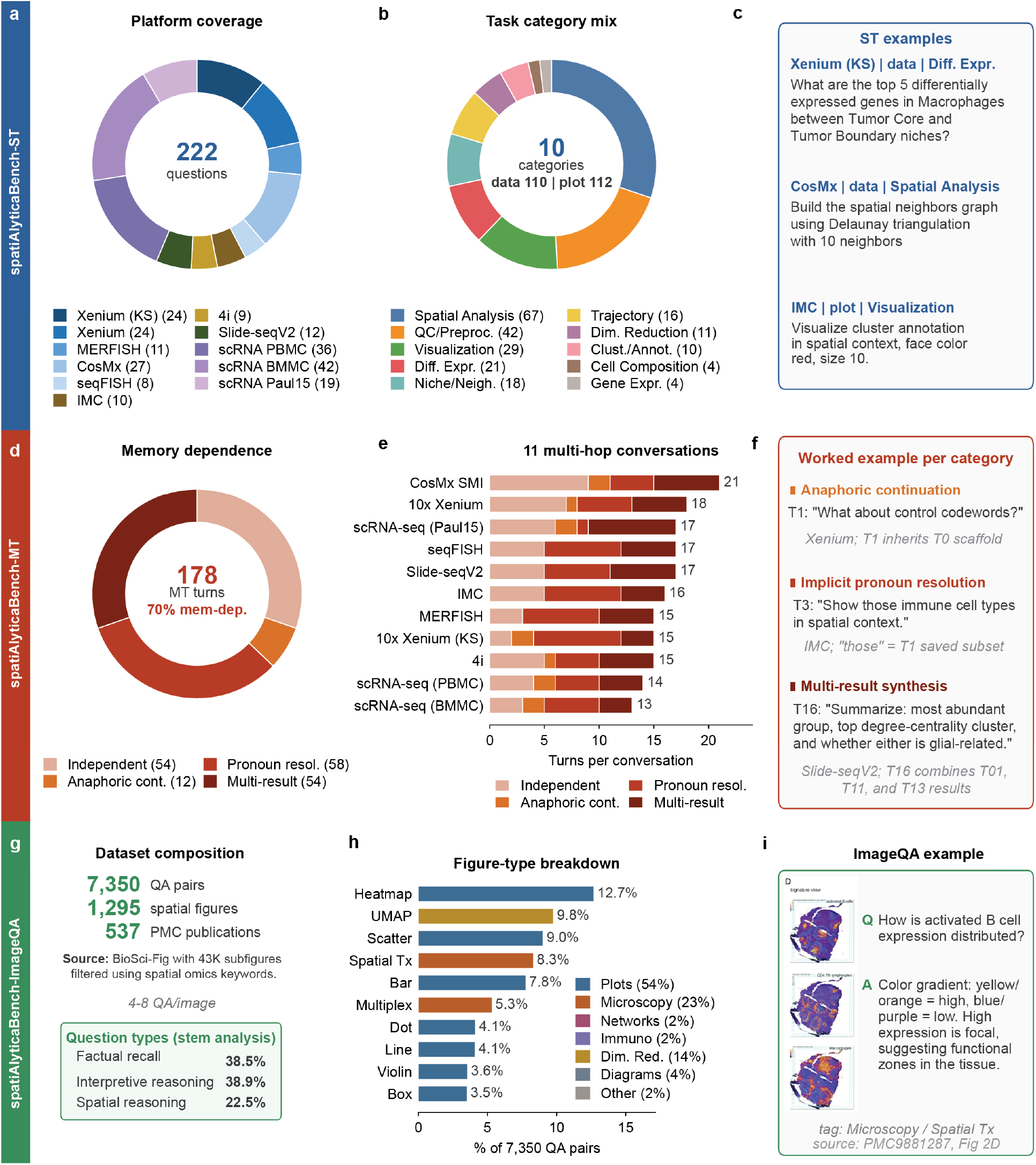
spatiAlyticaBench: a unified spatial omics conversational-analysis benchmark in three flavors (§1.2). (A–C) *spatiAlyticaBench-ST* (222 single-turn code-generation questions): (A) platform coverage across seven spatial omics platforms and three single-cell RNA-seq references; (B) ten task categories split between data-returning (110) and plot-saving (112) outputs; (C) representative ST questions with their `platform / output-type / category` tagging and `Evaluator`-class ground-truth specification. (D–F) *spatiAlyticaBench-MT* (178 turns across 11 multi-hop conversations of 13–21 turns each): (D) memory-dependence breakdown (124 of 178 turns memory-dependent); (E) per-conversation turn counts under the fourway memory-dependence taxonomy; (F) worked example per memory-dependent category, annotated with the prior-turn anchors that resolve each follow-up. (G–I) *spatiAlyticaBench-ImageQA* (7,350 question–answer pairs over 1,295 spatial omics subfigures from 537 PMC publications): (G) dataset composition with question-stem analysis; subfigures separated from BioSci-Fig [30] bounding-box annotations and processed through a LLaVA-Med-style GPT-4o pipeline; (H) figure-type breakdown grouped under seven parent categories; (I) representative ImageQA pair (source: PMC9881287, Fig. 2D). Per-platform, per-task, and per-conversation breakdowns: Supplementary Tables 2, 3, 4, 5.

**spatiAlyticaBench-ST** tests one-shot translation of a natural-language request into an analysis pipeline. We adapted the publicly hosted Squidpy and Scanpy tutorial collections and in-house datasets to 222 independent questions, each pairing a query with a pre-loaded `AnnData` object and a ground-truth Python implementation in a standardized `Evaluator` class (Fig. 2a–c; Methods). The questions were grouped to 10 bioinformatic tasks spanning spatial analysis, quality control, visualization, differential expression analysis, niche analysis, and more (Supplementary Table 4).

**spatiAlyticaBench-MT** captures dependencies that single-turn evaluation cannot: a follow-up like *“what about in macrophages?”* is unanswerable without conversation history, and intermediate-result errors cascade across turns. To our knowledge, this is the first systematic multi-turn conversational benchmark in agentic spatial biology [29]. We composed 11 multi-hop conversations of 13–21 turns each (178 turns total; Fig. 2e) on the same 11 datasets used in ST, structured around three categories of memory dependence: *anaphoric continuation, implicit pronoun resolution*, and *multi-result synthesis* (Fig. 2d,f). 124 of 178 turns (69.7%) are memory-dependent; multi-result synthesis forms a dedicated *memory retrieval* task category exclusive to this subset (Supplementary Table 5).

**spatiAlyticaBench-ImageQA** tests viewer-grounded visual reasoning. It comprises 7,350 question– answer pairs over 1,295 spatial transcriptomics subfigures from 537 distinct PMC publications (Fig. 2g– i). Subfigures were selected from the BioSci-Fig dataset [30] and QA pairs generated through a LLaVA-Med-style [31] pipeline (Methods). Because both questions and reference answers are LLM-generated, we calibrated the headline judge against human evaluation and a cross-family reference-free protocol [32, 33] (§1.5; Methods).

### 1.3. spatiAlytica outperforms baseline agents on single-turn analytical queries with higher efficiency

We evaluated spatiAlytica across four LLM backends (Supplementary Table 6) on *spatiAlyticaBench-ST*. Each question provided a natural-language query and a pre-loaded `AnnData` object; generated code was compared against ground-truth implementations via the hierarchical comparison pipeline (Methods). We benchmarked against BioMedAgent and BioMANIA (Scanpy and Squidpy modes) under matched backends (Fig. 3a; Supplementary Tables 7, 8, 9). No study-context prompt was injected in any benchmark run (Methods).

**Fig. 3.**
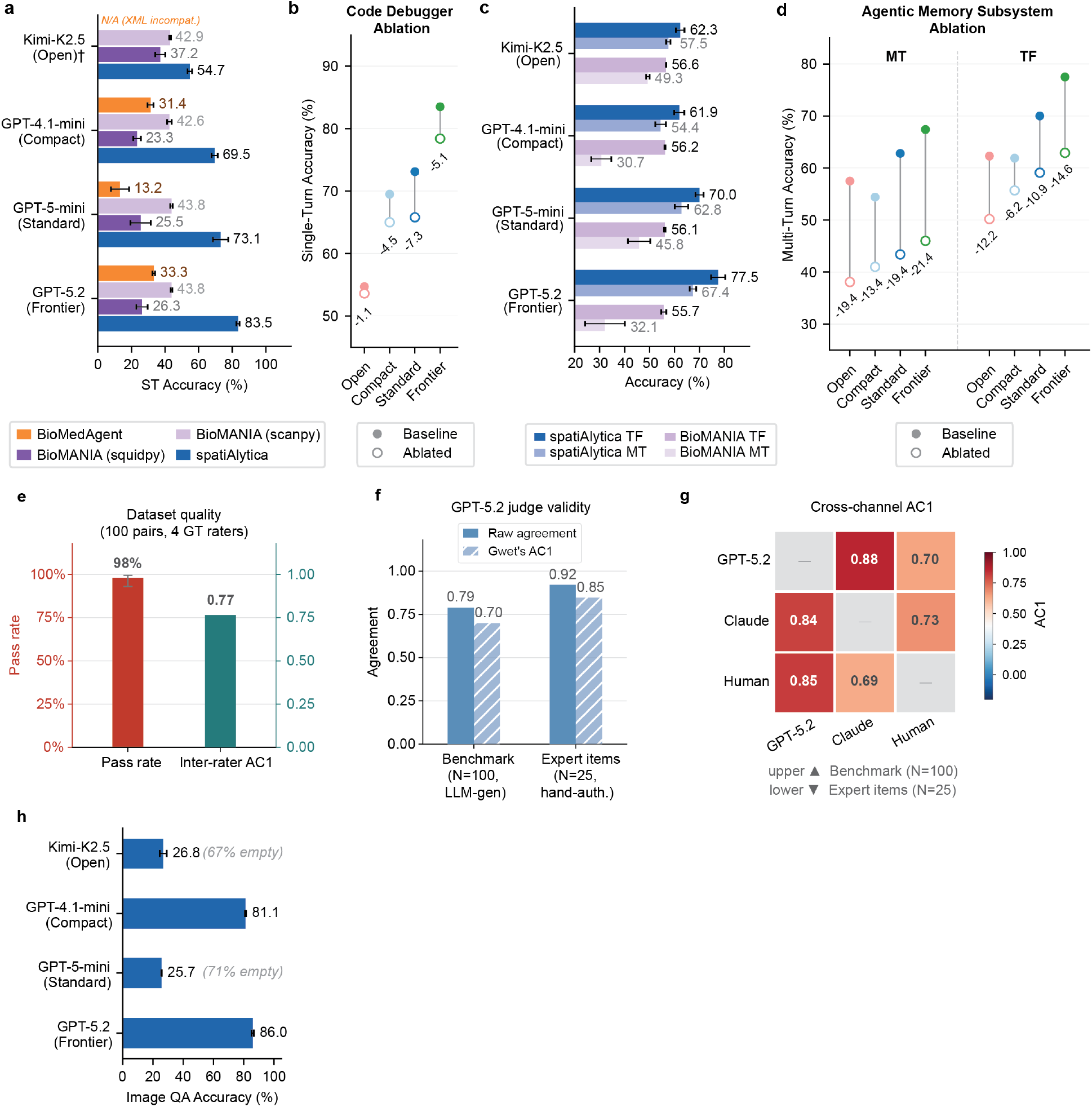
Benchmark evaluation of spatiAlytica’s analytical outputs. Code QA panels (A–D) report mean *±* s.d. across *N* =3 runs; spatiAlyticaBench-ImageQA (H) across *N* =2. (A) spatiAlyticaBench-ST accuracy across four LLM backends (222 questions) for spatiAlytica, BioMANIA (by library), and BioMedAgent; BioMedAgent+Kimi-K2.5 was not evaluable (Supplementary Tables 8, 9). (B) No-debugger ablation on spatiAlyticaBench-ST: baseline (filled) vs. ablated (open); deltas labeled. (C) spatiAlyticaBench-MT (178 questions) under cascading (MT) and teacher-forcing (TF) protocols: spatiAlytica vs. macro-averaged BioMANIA, per backend. (D) No-memory ablation on spatiAlyticaBench-MT (MT and TF); baseline vs. ablated; deltas labeled. (E) Validation dataset quality on the 100-pair spatiAlyticaBench-ImageQA subsample (*N* =4 human raters): human pass rate and inter-rater Gwet’s AC1. (F) GPT-5.2 judge validity against the human majority: raw agreement and Gwet’s AC1 on LLM-generated benchmark items (*N* =100) and hand-authored expert items (*N* =25). (G) Pairwise Gwet’s AC1 among GPT-5.2, Claude Opus 4.6, and the human majority; upper triangle, benchmark (*N* =100); lower triangle, expert items (*N* =25). (H) spatiAlyticaBench-ImageQA judge-pass rate (7,350 pairs) across four backends; empty-response fractions annotated for Standard and Open-source. Details in Supplementary Note 1.5.

spatiAlytica consistently outperformed both baselines across all backends (Fig. 3a; Supplementary Tables 7, 8, 9). In the Frontier configuration, it reached 83.5 *±* 0.9% (Wilson 95% CI [80.5, 86.1]) versus 43.8 *±* 0.5% / 26.3 *±* 3.4% for BioMANIA (Scanpy/Squidpy) and 33.3 *±* 0.8% for BioMedAgent, margins of 14.4–47.6 percentage points (pp) and 38–60 pp, respectively. Accuracy scaled with backend capability (54.7% to 83.5%), whereas BioMANIA accuracy was nearly flat across Scanpy backends (42.6–43.8%), and BioMedAgent accuracy was lower across all commercial backends; the gap widened at stronger backends, indicating that architectural contribution grows with model capability.

Ablating the code debugger sub-agent reduced single-turn accuracy by 4.5–7.3 percentage point (pp) across commercial backends (Fig. 3b; Supplementary Table 10), confirming that automated error diagnosis recovers a meaningful fraction of first-pass failures, most pronounced for mid-capability models and limited for the open-source backend where errors were less amenable to repair. These gains came at lower compute: 2.63 LLM calls per question versus 4.65 (BioMANIA) and 36.53 (BioMedAgent), with Frontier mean latency 89.6 s versus 180.3 s and 313.7 s (Supplementary Table 11). It is to be noted that single-turn accuracy measures one-shot correctness. However, interactive spatial biology is inherently sequential, with follow-up queries reusing prior results and errors propagating across turns.

### 1.4. Agentic memory limits cascading errors in multi-turn spatial analysis

We evaluated spatiAlytica on *spatiAlyticaBench-MT* to test context-dependent multi-turn analysis. Two protocols were used: natural multi-turn (MT), where the agent carries forward its own outputs, and teacher forcing (TF), where each turn receives the correct ground-truth `AnnData` state, isolating per-turn accuracy from cascading error. We compared with BioMANIA under matched backends but BioMedAgent was not evaluated under this protocol because it does not support multi-turn interaction, and no other cited agentic-biology system (AutoBA, CellAgent, SpatialAgent, scChat) reports multi-turn conversational evaluation. During evaluation, no study-context prompt was injected in any run. Under MT, spatiAlytica outperformed BioMANIA across all backends (Fig. 3C; Supplementary Tables 12, 13). spatiAlytica accuracy ranged from 54.4 *±* 2.1% (Compact) to 67.4 *±* 1.3% (Frontier, conversation-level bootstrap 95% CI [65.8, 75.3]; Supplementary Table 14), while BioMANIA ranged from 19.6–52.4%. Notably, spatiAlytica with the open-source backend (57.5 *±* 0.9%) exceeded the Compact back-end despite lower single-turn accuracy, suggesting that multi-turn performance depends strongly on the agent architecture, not only on model strength. Teacher forcing further separated per-turn accuracy from error accumulation. Under TF, spatiAlytica increased with backend strength (62.3→ 77.5%), whereas BioMANIA remained nearly flat (55.1–57.1%). The smaller MT → TF gap for (4.8–10.1 pp vs. 13.2–35.8 pp for BioMANIA) shows that spatiAlytica preserves prior results more reliably, approaching its own TF upper bound.

To test which sub-system drives this near TF upper-bound behavior, we ablated the agentic memory sub-system and the code debugger on spatiAlyticaBench-MT (Fig. 3d; Supplementary Table 10). Removing memory reduced MT accuracy by 13.4–21.4 pp across backends, whereas removing the code debugger had negligible MT effect (− 1.6 to +1.2 pp). Among the two ablated sub-systems, the multi-turn-specific memory, not execution retry, is the dominant driver of multi-turn accuracy on this benchmark, consistent with the benchmark’s memory-dependent design. Higher MT accuracy also came with low per-query compute (45–74 k tokens; $0.009–$0.039 per query; Supplementary Table 12).

### 1.5. spatiAlytica produces expert-calibrated image-grounded responses

We evaluated the Spatial VQA sub-agent on *spatiAlyticaBench-ImageQA*. Responses were evaluated using a fixed LLM judge (GPT-5.2), following established LLM-as-judge protocols [32, 33]. The judge scored each response against the reference answer on a 1–5 scale, with scores ≥ 3 counted as passes.

Because both benchmark questions and answers were generated by GPT-4o, we first validated the spatiAlyticaBench-ImageQA itself through blinded human review (Methods). On a 100-item stratified sub-sample, four bioinformaticians confirmed high benchmark quality, with a 98% pass rate and inter-rater Gwet’s AC1 of 0.77 (Fig. 3e), supporting that the generated QA pairs were biologically meaningful and well grounded in the underlying images.

We next evaluated the reliability of the GPT-5.2 judge (Methods). On the same benchmark subset, GPT-5.2 tracked the human majority vote well (raw agreement 0.79; AC1 0.70), and agreement was even higher on 25 hand-authored expert items (0.92 raw agreement; AC1 0.85; Fig. 3f). Under a reference-free comparison, GPT-5.2, Claude, and the human majority also showed similar cross-channel agreement (AC1 0.69–0.88; Fig. 3g), indicating that evaluation is not an artifact of single model family self-reference and reducing concerns about circularity.

Across backends, the Frontier (GPT-5.2) configuration achieved a pass rate of 85.9 *±* 0.7% and the Compact (GPT-4.1-mini) configuration 81.1 *±* 0.4% (Fig. 3h, Supplementary Table 15, 16). Lower Standard and Open-source performance was largely driven by empty responses rather than incorrect visual reasoning; among non-empty responses, pass rates exceeded 84% across all configurations (Supplementary Table 17). These results show that the spatial VQA sub-agent produces image-grounded responses consistent with LLM-generated references and calibrated to expert judgment on a stratified subsample.

### 1.6. spatiAlytica recapitulates stage-dependent architecture and viral tropism in Kaposi’s sarcoma

To demonstrate spatiAlytica’s agentic workflow for hypothesis-driven spatial discovery, we applied it to a Xenium Kaposi’s sarcoma (KS) [34, 35] dataset profiling 18 tissue microarray cores across all four disease stages (control, patch, plaque, nodular; 301,689 cells, 308-gene panel; Methods) [36]. A brief study context defined KSHV^+^ lymphatic endothelial cells (LECs) as tumor cells and KSHV-prefixed genes as infection markers (full interaction transcript: Supplementary Note 6), grounding all downstream reasoning in the biological domain and enabling subsequent natural-language queries to drive the EA-IR workflow.

We first used spatiAlytica to characterize the coordinated architectural shift during KS progression through stage-stratified cell type composition (EA1; Fig. 4a) and visualization of spatial cell-type distributions (EA2; Fig. 4b). Control tissue was dominated by keratinocytes and spinous-to-granular cells with minimal immune or tumor populations; patch lesions showed the first emergence of KSHV^+^ LEC tumor cells together with modest fibroblast, macrophage, and CD8 T-cell expansion. Plaque lesions exhibited peak fibroblast and macrophage abundance alongside continued tumor-cell accumulation, while nodular disease was marked by dominant KSHV^+^ LEC tumor populations, sustained macrophage enrichment, and near-complete depletion of keratinocytes. Prompted to synthesize these signals, spatiAlytica produced a stage-resolved trajectory (IR1; Fig. 4c) describing progressive epidermal erosion, stromal and vascular expansion, tumor consolidation, and a transition from early immune engagement in patch lesions to immune dilution in advanced disease, consistent with published KS pathology [36].

**Fig. 4.**
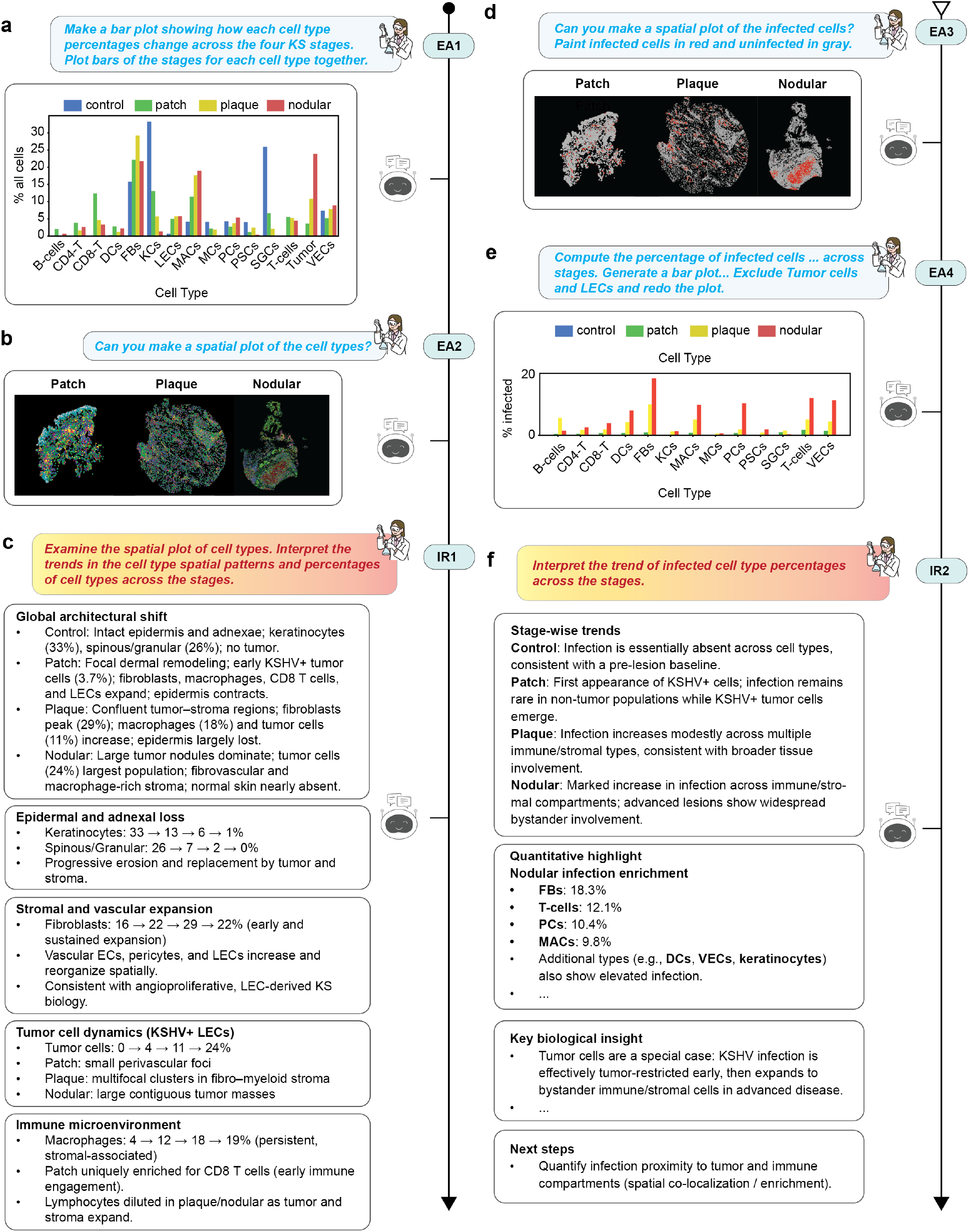
Agentic analysis recapitulates cell-type and infection-associated remodeling during Kaposi’s sarcoma progression. (A) Cell-type composition across four KS stages showing progressive loss of keratinocytes and expansion of fibroblast, macrophage, and tumor populations (EA1). (B) Spatial visualization of annotated cell types across representative patch, plaque, and nodular cores revealing progressive architectural reorganization (EA2). (C) Interpretive synthesis integrating compositional and spatial trends into a stage-resolved trajectory of KS progression (IR1). (D) Spatial maps of KSHV infection status showing stage-dependent changes in viral burden and distribution (EA3). (E) Infected cell percentages stratified by cell type and disease stage, including reanalysis excluding tumor cells to reveal infection in non-tumor compartments (EA4). (F) Interpretive synthesis describing progression from tumor-restricted infection in early lesions to widespread involvement of immune and stromal cell types in nodular disease (IR2).

We next queried KSHV infection patterns across stages (EA3–4; Fig. 4d,e), and spatiAlytica revealed sparse infected cells in patch, broader distribution in plaque, and large spatially consolidated infected regions in nodular disease. Asked to interpret these trends, spatiAlytica recovered the published KSHV tropism model (IR2; Fig. 4f): infection is absent in controls, initiates in LEC-lineage tumor cells in patch, expands to immune and stromal bystanders in plaque, and becomes widespread across non-tumor compartments in nodular disease [36].

### 1.7 spatiAlytica reveals progressive CD8-T cell dysfunction during Kaposi’s sarcoma progression

Building on these cell-type shifts, we used spatiAlytica for niche-based analysis of the spatial immune microenvironment [37, 38]. A natural-language request yielded ten recurrent niches, automatically annotated by composition (EA5; Fig. 5a; full interaction transcript: Supplementary Note 7). Stage-resolved quantification (EA6; Fig. 5b) and spatiAlytica’s interpretation (IR3; Fig. 5c) recapitulated the progressive remodeling [36]: keratinocyte-rich niches (∼ 75%) in control, expanded immune-infiltrated (∼ 38%) and fibroblast-enriched (∼ 21%) niches in patch, a shift toward fibro-tumor stroma with contracting immune niches indicative of exclusion in plaque, and tumor-core (∼ 19%) and macrophage-rich stromal niches in nodular disease.

**Fig. 5.**
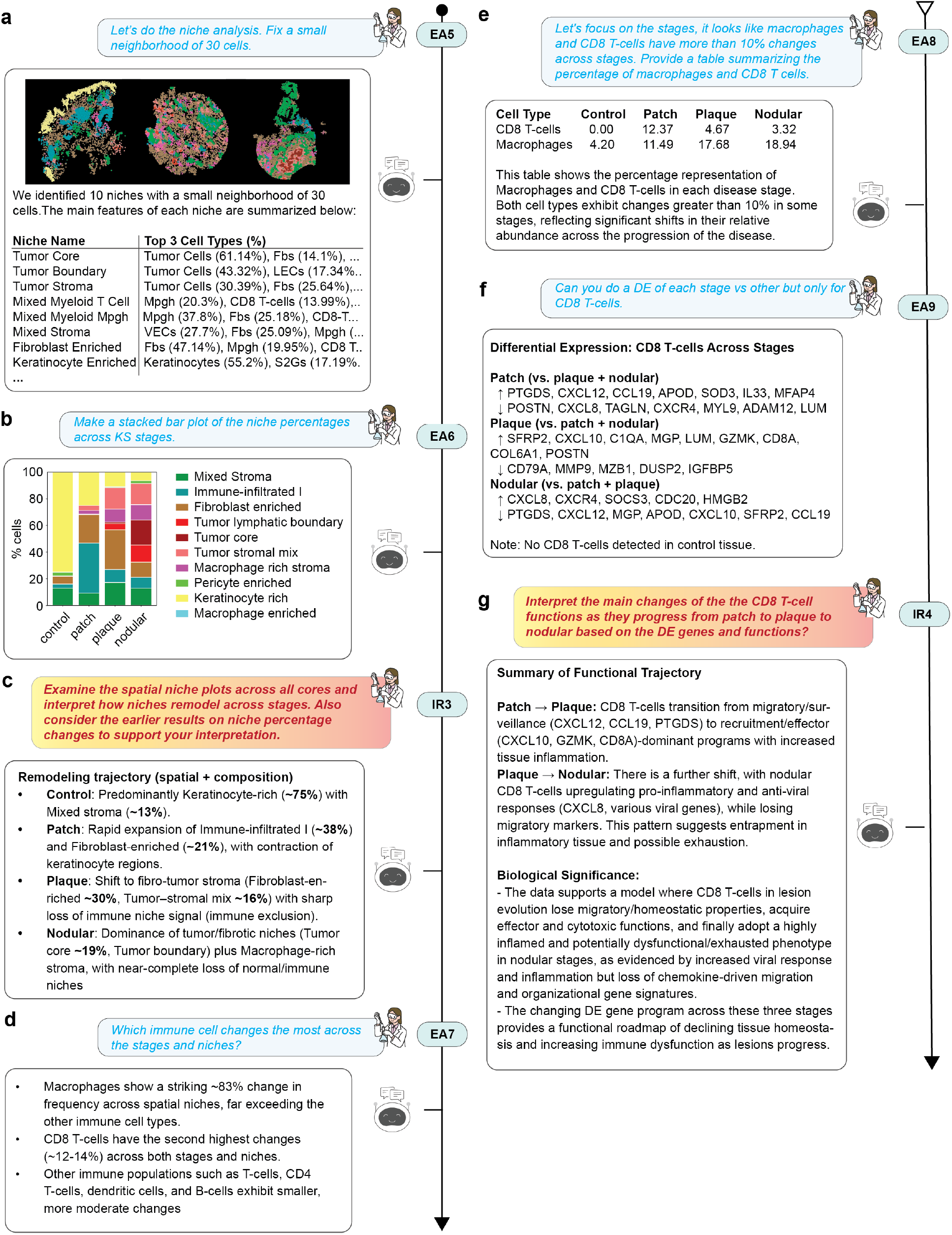
Agentic niche and T-cell analysis identifies stage-dependent immune remodeling during Kaposi’s sarcoma progression. (A) Neighborhood-based niche analysis using a 30-cell radius identifies ten recurrent spatial niches across KS stages, with automatic biological naming based on dominant cell-type composition (EA5). (B) Niche composition quantified across stages showing systematic remodeling from keratinocyte-rich to tumor-and stroma-dominated niches (EA6). (C) Interpretive synthesis integrating spatial distributions and niche abundances into a stage-resolved trajectory of microenvironmental remodeling (IR3). (D) Immune population comparison across stages and niches identifies macrophages and CD8 T-cells as the most dynamically remodeling compartments (EA7). (E) Stage-resolved quantification showing non-monotonic CD8 T-cell trajectory (peak in patch, progressive decline in plaque and nodular) contrasting with continuous macrophage accumulation (EA8). (F) Differential expression analysis revealing stage-dependent transcriptional programs in CD4 and CD8 T-cells, with recruitment genes in patch, ECM remodeling in plaque, and tissue retention in nodular (EA9). (G) Integrated interpretation supporting a model of progressive T-cell functional impairment from active cytotoxic engagement to a tissue-retained, functionally diminished state (IR4).

Immune-cell comparison across stages and niches (EA7; Fig. 5d) identified macrophages (∼ 83% niche variation) and CD8-T cells (∼ 12–14%) as the most dynamic populations. Stage-resolved quantification (EA8; Fig. 5e) showed a non-monotonic CD8-T trajectory (0%, 12.37%, 4.67%, 3.32% across stages), opposite to continuous macrophage accumulation (from 4.20% to 18.94%), extending prior reports of an inverse macrophage-CD8-T relationship in KS [39] and motivating analysis of CD8-T function.

Stage-specific differential expression (EA9; Fig. 5f) yielded distinct stage signatures. Patch lesions upregulated chemoattraction and recruitment genes (*CXCL12, CCL19, PTGDS*) and effector markers (*GZMK, IL33*) while downregulating fibroblast-inflammatory genes (*POSTN, CXCL8, LUM*). Plaque lesions upregulated interferon-responsive (*CXCL10*), effector (*GZMK, CD8A*), and matricellular genes (*SFRP2, MGP, LUM*). Nodular lesions upregulated stress, inflammatory, and retention genes (*SOCS3, HMGB2, CXCR4, CXCL8*) while downregulating early recruitment and interferon-responsive genes (*PTGDS, CXCL12, CCL19, CXCL10*). spatiAlytica’s interpretation (IR4; Fig. 5g) described a progression from migratory surveillance in patch, through cytotoxic engagement with the desmoplastic stroma in plaque, to tissue-retained dysfunction in nodular disease, a CD8-T cell functional shift not previously reported for KS [36].

### 1.8. spatiAlytica reproduces subtype-specific niches and outcome-linked immune states in colorectal cancer

To demonstrate generalization beyond spatial transcriptomics, we applied spatiAlytica to 56-marker CODEX [5] CRC data (140 leukocyte-dense intratumoral regions across Crohn’s-like reaction (CLR) and diffuse inflammatory infiltration (DII) subtypes [37]; full interaction transcript: Supplementary Note 9). Subtype-stratified spatial visualization and composition queries (EA1–2; Fig. 6a,b) and spatiAlytica’s interpretation (IR1; Fig. 6c) recovered the published CLR-DII contrast: CLR was B-cell-enriched with organized lymphoid and TLS-like structure, whereas DII was granulocyte-and macrophage-enriched with a diffuse, innate-inflammatory pattern.

**Fig. 6.**
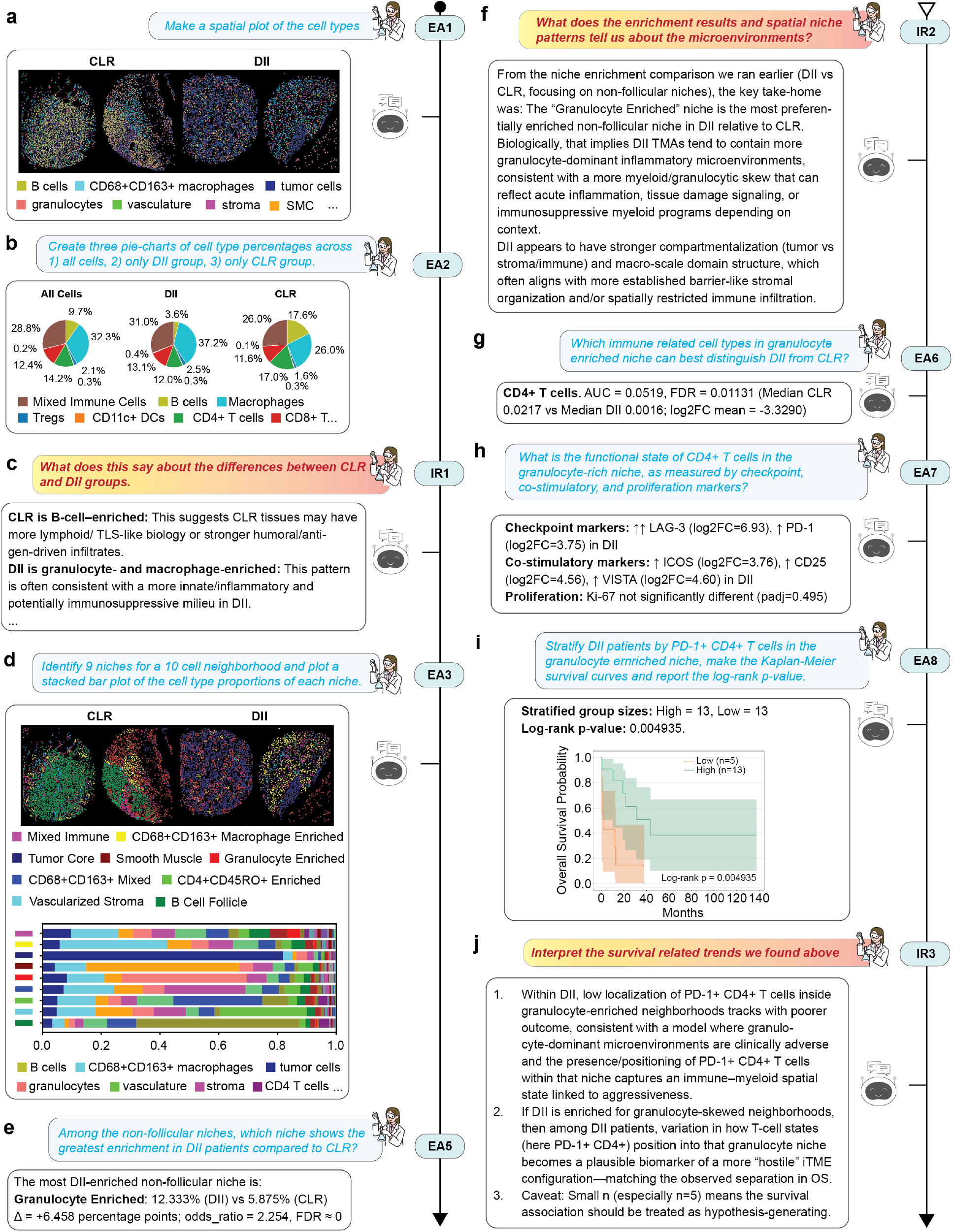
spatiAlytica reproduces subtype-specific niche organization and survival-linked immune states in CODEX colorectal cancer. (A) Spatial plots of cell types across representative CLR and DII cores (EA1). (B) Cell-type proportions for all cells, DII, and CLR groups (EA2). (C) Interpretive summary of CLR-DII compositional differences highlighting organized lymphoid structure in CLR versus diffuse infiltration in DII (IR1). (D) Neighborhood niche analysis identifying nine spatial niches with automatic biological naming and per-niche cell-type proportions (EA3). (E) Granulocyte Enriched niche identified as the most DII-enriched non-follicular niche (Δ = +6.458 pp; odds ratio = 2.254) (EA5). (F) Interpretation of niche enrichment and spatial microenvironment structure (IR2). (G) CD4^+^ T cells identified as the top immune discriminator between subtypes within the granulocyte-enriched niche (EA6). (H) CD4^+^ T-cell functional state characterized by checkpoint, co-stimulatory, and proliferation markers within the granulocyte-enriched niche (EA7). (I) Kaplan–Meier curves stratifying DII patients by PD-1^+^ CD4^+^ T-cell localization in granulocyte-enriched neighborhoods (log-rank *p* = 0.005; high *n* = 13, low *n* = 5) (EA8). (J) Interpretation linking PD-1^+^ CD4^+^ T-cell spatial positioning to clinical outcome in DII, flagged as hypothesis-generating given the small cohort size (IR3).

A niche-analysis request yielded nine recurrent niches, automatically annotated by composition in labels matching those in the original study (EA3; Fig. 6d). Subtype-enrichment analysis (EA4–5, IR2; Fig. 6e,f) identified the B-cell follicle/TLS-like niche as CLR-dominant, consistent with TLS-associated CLR biology [40–42], and a granulocyte-enriched niche as the top additional DII discriminator, both reproducing the published analysis.

We then focused on the granulocyte-enriched niche to link its spatial signature to immune function and clinical outcome. Cell-type stratification (EA6; Fig. 6g) recovered the reported CD4-T cell subtype difference, and marker characterization (EA7; Fig. 6h) reproduced the checkpoint and co-stimulatory marker profiles distinguishing the subtypes. A survival analysis query showed that DII patients with low PD1^+^ CD4-T cell localization in granulocyte-enriched neighborhoods had significantly poorer overall survival (log-rank *p* = 0.005; EA8; Fig. 6i). spatiAlytica’s synthesis (IR3; Fig. 6j) flagged the association as hypothesis-generating given the small cohort (*n*=18; high *n*=13, low *n*=5), illustrating the anti-hallucination constraints that restrict claims to the data. Together, this sequence shows spatiAlytica carrying a multi-step CODEX analysis from niche discovery to within-niche immune-state characterization and outcome association [43] in a single conversational workflow.

spatiAlytica further generalized to Visium data, recovering chemotherapy-driven cancer-associated fibroblast (CAF) remodeling in high-grade serous ovarian cancer (HGSOC) [44] and confirming cross-platform applicability (Supplementary Note 5; full interaction transcript: Supplementary Note 8; Supplementary Fig. 4).

### 2 Discussion

Real-world spatial biology analysis is both technically demanding and inherently iterative and hypothesis-driven. A biologist inspects a spatial plot, forms a hypothesis, conducts analyses, interprets the result in biological context, and asks a follow-up that depends on prior turns. Supporting this workflow requires tight integration with the spatial viewer, persistent analytical state, and biologically grounded interpretation, capabilities that are particularly important for biologists without programming expertise. Existing agentic biology systems, despite strengths in conversational flexibility or domain-optimized execution, do not jointly provide these features: most remain text-centered, lack coupling to the spatial viewer, and cannot maintain analytical state across turns (Supplementary Table 1). Our results suggest that these are not peripheral features, but architectural requirements for interactive spatial analysis.

spatiAlytica addresses these gaps through a viewer-integrated, memory-equipped multi-agent architecture. To evaluate these requirements, we introduced *spatiAlyticaBench*, a three-part benchmark spanning one-shot executable analysis, context-dependent multi-turn reasoning, and viewer-grounded visual interpretation. Across these benchmarks, each architectural component contributed measurably. In the most relevant sequential multi-turn setting, spatiAlytica reached 67.4% versus 32.1% for BioMANIA; single-turn accuracy also showed a consistent advantage (83.5% vs. 43.8%/33.3%), with gaps widened at stronger backends. These indicate that system architecture, not model scale alone, is the current bottleneck for agentic spatial biology. Particularly, structured memory, not code retry, drove multi-turn robustness: ablating memory reduced accuracy by 13.4–21.4 pp, while ablating the Code Debugger had minimal multi-turn effect. The smaller MT-TF gap for spatiAlytica (4.8–10.1 pp vs. 13.2–35.8 pp for BioMANIA) confirms that cascading state errors, rather than per-turn reasoning, are the primary bottleneck in multi-step analysis. Visual grounding extends this further: by coupling the Spatial VQA Agent to the Napari viewer, spatiAlytica supports queries tied to the current field of view and spatial relationships, a capability absent from text-only systems, that enabled direct queries about KSHV infection patterns in specific spatial regions during the Kaposi’s sarcoma (KS) case study.

These architectural choices yield biological insight. In KS, spatiAlytica recapitulated published findings and revealed a progressive CD8 T-cell dysfunction trajectory, from migratory surveillance in patch lesions, through cytotoxic engagement in plaque, to tissue-retained dysfunction in nodular disease, a staged functional shift not previously reported in KS that may inform therapeutic targeting of the tumor immune microenvironment. In colorectal cancer and ovarian cancer, the system reproduced subtype-specific niche architectures and cross-platform applicability, with the EA-IR loop enabling progressive hypothesis refinement within single conversational sessions.

Several limitations remain, including dependence on an upfront study context for grounded responses, variability and cost inherited from LLM backends, and reduced Spatial VQA performance on very dense plots. More broadly, our evaluations remain system-level assessments, and broader validation across users, assays, and multimodal settings will be needed. Even so, these results suggest that future AI systems for spatial biology should move beyond text-only assistance toward viewer-grounded, stateful, and interpretation-aware workflows.

## 3 Methods

### 3.1. System architecture

spatiAlytica implements a multimodal interactive agentic architecture that couples LLMs [18] to an interactive, image-centered Napari viewer [27] and a reproducible Python analysis engine. All LLM calls are dispatched through LiteLLM [45], which provides a unified interface across providers (OpenAI, Anthropic, Together AI, and local endpoints). The architecture is organized into four layers: (i) a *widget layer* (Napari dock widget) for dataset loading, column configuration, spatial and scientific plot visualizations, and legends; (ii) an *agent layer* in which an orchestrator sub-agent decomposes queries into tool calls and delegates to five specialized sub-agents (data resolver, code writer, code debugger, spatial VQA sub-agent, and user response sub-agent), all operating over a shared agentic memory sub-system; (iii) a *hybrid tool repository* of composable, type-validated tools for data inspection, statistical analysis, spatial methods, and visualization; and (iv) a *prompt layer* of role-specific, context-aware templates that enforce grounding in tool outputs and reduce hallucination. The system maintains an explicit UI state representation (active layers, legends, viewport, annotations) that is updated after each action and used by agents to interpret follow-up queries tied to the current view, enabling closed-loop coupling between computation, visualization, and agent reasoning (Supplementary Note 1).

We use this multimodal interactive agentic system to capture three integrated capabilities of spatiAlytica: 1) multimodality (natural-language queries, serialized viewer state, rendered images, and executable code as first-class inputs and outputs), 2) interactivity (conversational, viewer-coupled refinement across turns), and 3) an orchestrator/sub-agent decomposition in which named, delegated sub-agents share a structured workspace, consistent with the classical blackboard architecture [46]. Cooperative inter-agent dynamics in spatiAlytica include the code writer to code debugger correction cycle on shared code artifacts and the user response sub-agent’s grounded synthesis over upstream sub-agents’ outputs.

### 3.2 Core sub-agents

The orchestrator sub-agent [47, 48] is the central coordinator: it classifies each query as exploratory analysis (EA) or interpretation (IR), retrieves context from the agentic memory sub-system, selects and sequences tools, and delegates to specialized agents. Tool execution proceeds in an iterative loop (up to 10 iterations) until a final response is produced (Supplementary Fig. 1b). The **data resolver sub-agent** maps biological terms onto concrete `AnnData` keys through two-phase LLM-based namespace classification and candidate selection; the orchestrator additionally performs local fuzzy matching on column and gene names (via rapidfuzz/SequenceMatcher) and delegates to the data resolver sub-agent when the LLM is needed, caching results to avoid redundant LLM calls.

The **code writer sub-agent** generates executable Python code after an internal planning loop (up to 10 rounds) that inspects dataset structure and validates column names. The **code debugger sub-agent** corrects execution failures through an execute-correct-retry cycle (up to 3 attempts), handling common spatial omics failure patterns including categorical mismatches, sparse matrix extraction, and API misuse. The **spatial VQA sub-agent** performs multimodal interrogation of spatial visualizations by accessing the UI state representation and analyzing Napari canvas screenshots. It also carries out interrogation of the scientific plots created in the resizable Matplotlib widget in spatiAlytica UI. The **user response subagent** synthesizes tool outputs into structured narrative responses grounded in recorded results. Because interpretation is conditioned on the active visualization state, responses remain synchronized with the user’s current view (e.g., a zoomed-in tumor–stroma interface) and the exact color mappings and overlays currently displayed. Detailed agent specifications are provided in Supplementary Note 1.

### 3.3. Data handling

spatiAlytica natively supports `AnnData` [28] as the primary data representation, exposing expression matrices, per-cell and feature metadata, and gene identifiers. Spatial coordinates are read from configured columns and used to generate Napari layers.

### 3.4. Hybrid tool repository and code execution

The hybrid tool repository exposes 13 type-validated tools to the orchestrator via function-calling schemas, spanning four categories: data inspection (column resolution, value enumeration), analytical workflows (differential expression, niche analysis, spatial autocorrelation, cell density), visualization (spatial plots, UMAP, pie charts), and image analysis (multimodal VQA). Structured tools use Pydantic schemas for parameter validation, and tools can be chained in sequences of up to 10 iterations per query. When a request requires bespoke computation, the `WriteCodeTool` delegates to the code writer sub-agent, which generates Python code executed in a controlled environment with pre-imported spatial omics libraries (Scanpy [49], Squidpy [38], Matplotlib, seaborn). Spatial plots are routed through the `SpatialPlotTool` to maintain Napari layer interactivity. All stochastic operations use `random state=42`. Failed executions trigger the code debugger’s iterative correction cycle (up to 3 attempts). The complete tool inventory is described in Supplementary Note 1.

### 3.5. Visualization and UI state management

Spatial visualizations are rendered as interactive Napari layers with categorical or continuous colormaps and synchronized legends. Non-spatial scientific visualizations are rendered through a resizable Matplotlib widget integrated directly inside spatiAlytica UI. The system maintains an explicit UI state representation encoding active layers, color mappings, annotations, and viewport configuration. This state is updated after each action and stored in the agentic memory sub-system, enabling agents to interpret follow-up requests tied to the current view (e.g., “in this region” or “compare the two visible layers”). This closed-loop coupling between tools, visualization, and agent reasoning supports iterative hypothesis-driven exploration. Furthermore, both the spatial and scientific visualizations can be exported to a variety of vector and raster image formats.

### 3.6. System prompt design

Sub-agent behavior is governed by role-specific, context-aware prompt templates that define tool-use policies, dataset access rules, error handling, and visualization conventions. Prompts constrain agents to (i) request resolution of ambiguous dataset references via the data resolver sub-agent, (ii) ground quantitative claims in tool outputs and executed code, and (iii) provide reproducible narratives that reference the performed computations.

### 3.7. Agentic memory sub-system and interaction logging

spatiAlytica maintains a shared agentic memory sub-system, implemented using LangChain [50], that provides three tiers of state management: (i) *short-term memory* for conversation history; (ii) *long-term memory* for validated column mappings, analysis state, and a registry of persisted results (columns, embeddings, and stored objects added to the `AnnData` object); and (iii) *intermediate result memory* for recent tool outputs and tool call specifications. The system additionally generates conversation summaries to provide condensed context for downstream agents. All memory is persisted to disk, enabling session continuity. For each query, the orchestrator assembles a short structured context dictionary (conversation summary, dataset metadata, column mappings, recent tool results, and viewport state) that is appended to the agent’s system prompt, enabling follow-up handling and incremental refinement across turns. Comprehensive session-based interaction logs capture all queries, agent responses, executed code, generated visualizations, and structured telemetry for full analytical provenance and workflow replay (Supplementary Note 1).

### 3.8. Study context design

Each analysis session can optionally begin with a structured **study context** that grounds agent reasoning in the biological and technical context of the dataset. The study context follows a standardized template with six categories: (i) dataset overview (platform, tissue, sample organization); (ii) column naming conventions (mapping dataset-specific keys to semantic concepts such as cell type, disease stage, and sample identifier); (iii) experimental group definitions (value mappings and logical ordering for progression analyses); (iv) cell type interpretations (synonyms, hierarchical relationships, marker-based criteria); (v) visualization standards (color palettes, axis ordering); and (vi) analysis focus areas (scientific objectives and priority analyses). When provided, the complete study context is stored in the agentic memory sub-system upon session initialization and remains accessible to all agents, enabling downstream queries to reference prior definitions implicitly. A template and worked examples are provided in Supplementary Note 2.

### 3.9. Model configurations

We evaluated spatiAlytica under four model configurations (Table 6). The three proprietary configurations hold the user response, data resolver, and memory components at GPT-4.1-mini and progressively upgrade the orchestrator and code debugger from GPT-4.1-mini (Compact) through GPT-5-mini (Standard) to GPT-5.2 (Frontier); the Open-source configuration uses Kimi-K2.5 for every agent role via Together AI’s OpenAI-compatible endpoint. All LLM calls route through LiteLLM. For spatiAlyticaBench-ImageQA, the orchestrator model serves as the vision model and GPT-5.2 is the fixed judge across all configurations.

### 3.10. Benchmark datasets and evaluation

To systematically evaluate spatiAlytica, we constructed *spatiAlyticaBench* (Results §1.2), a unified benchmark partitioned into three flavors: *spatiAlyticaBench-ST* (single-turn code generation), *spatiAlyticaBench-MT* (sequential multi-turn code generation), and *spatiAlyticaBench-ImageQA* (viewer-grounded visual question answering). Together, these flavors assess code-generation correctness, visual-interpretation accuracy, and cross-turn context retention across 11 spatial transcriptomics datasets from 7 platforms and 3 scRNA-seq references (Supplementary Tables 2).

#### 3.10.1. spatiAlyticaBench-ST code QA benchmark

*spatiAlyticaBench-ST* (Results §1.2; 222 single-turn questions; independent; data and plot outputs) tests one-shot translation of a natural-language analytical request into a working Scanpy/Squidpy pipeline; per-platform, per-dataset, and per-task counts are reported in Results §1.2 and Supplementary Tables 3 and 4. Each QA pair consists of a natural-language question, a ground-truth Python implementation encapsulated in a standardized `Evaluator` class, and metadata specifying the output type (data or plot), required libraries (Scanpy, Squidpy, Matplotlib), and the expected result format. Questions were derived from the publicly hosted Squidpy and Scanpy tutorial collections and in-house datasets (the Kaposi’s sarcoma Xenium dataset contributes 24 ST questions independent of any tutorial) and enhanced with structured specifications to ensure unambiguous evaluation.

#### 3.10.2. spatiAlyticaBench-MT sequential multi-turn code QA benchmark

*spatiAlyticaBench-MT* (Results §1.2; 178 turns across 11 multi-hop conversations of 13–21 turns each; sequential; data and plot outputs) extends the code QA framework to interactive, multi-step analysis where each turn depends on the `AnnData` state produced by the preceding turns; per-conversation counts under the four-way memory-dependence taxonomy are reported in Results §1.2 and Supplementary Table 5. Each turn shares the same QA-pair structure as in *spatiAlyticaBench-ST* (natural-language question, `Evaluator`-class ground-truth Python implementation, output-type and library metadata), with the conversation defining the cumulative state expected at that turn. Conversations are designed to mirror how biologists iteratively explore spatial data, with three categories of memory-dependent questions of increasing difficulty: (i) anaphoric continuation, where questions like “What about in Macrophages?” are entirely ambiguous without knowing what analysis was performed in the prior turn; (ii) implicit pronoun resolution, where questions like “Is it one of the nuclear structure groups we identified earlier?” require resolving “it” to a specific prior result and cross-referencing with a subset defined several turns back; and (iii) multi-result synthesis, where questions like “Cross-reference all spatial results: the top enriched pair, the degree centrality champion, and the clustering champion” require recalling and combining distinct results scattered across the conversation. Each turn is annotated with a `FollowUp` flag indicating whether prior conversational context is required for resolution. This structure enables evaluation of the system’s ability to maintain analytical context, reuse intermediate results, and reason over accumulated evidence across turns.

#### 3.10.3. spatiAlyticaBench-ImageQA benchmark

The *spatiAlyticaBench-ImageQA* (Results §1.2; 7,350 QA pairs over 1,295 spatial omics subfigures) was constructed using BioSci-Fig [30], a scientific figure-separation dataset of 7,174 compound figures with 43,183 manually annotated subfigure bounding boxes drawn from biomedical literature. Because BioSci-Fig provides compound figures together with their subfigure-level bounding-box annotations, we used those bounding boxes to extract individual subfigures from compound figures published in spatial omics papers. Source papers were identified by keyword filtering on the title and abstract using the terms *spatial transcriptomics, ST, single-cell spatial transcriptomics, Xenium, Visium, CosMx, MERFISH, seqFISH, Slide-seq, IMC, spatial proteomics*, and *CODEX*, yielding 1,295 subfigure images from 537 distinct PMC publications. For each selected subfigure, QA pairs were generated following the LLaVA-Med [31] data curation pipeline: a structured prompt incorporating the figure, figure’s original caption and figure-referencing inline sentences from the source paper was passed to GPT-4o, which generated between 4 and 8 question–answer (QA) pairs per image.

Generated QA pairs were filtered to remove duplicates, trivially short answers, and questions not grounded in the visual content. Because both the reference answers and the QA generation are LLM-produced (GPT-4o), this benchmark assesses the spatial VQA sub-agent’s ability to produce spatially grounded descriptions consistent with LLM-generated references rather than human-adjudicated ground truth. Evaluation is performed using a fixed LLM judge (GPT-5.2) [32], distinct from the QA generation model, that scores each response on a 1–5 scale assessing semantic equivalence with the reference answer, accounting for paraphrasing and equivalent biological interpretations while penalizing factual errors and hallucinated spatial relationships. A score ≥ 3 is considered a pass. To validate this LLM judge against human expert judgment, we conducted a blinded human evaluation on a stratified subsample of 100 QA pairs (see below).

#### 3.10.4. Human validation of the spatiAlyticaBench-ImageQA judge

To validate the GPT-5.2 judge, we conducted a blinded human evaluation on a stratified subsample of 100 image QA pairs, balanced across three question types (factual, interpretive, spatial) and five judge-score bins (1–5), with 6–7 pairs per stratum to ensure calibration coverage at the pass/fail decision boundary (judge score ≥ 3). Nine distractor pairs, in which the model answer was replaced with an answer from an unrelated image–question pair, were interspersed per rater as attention checks (Supplementary Fig. 3d), yielding 109 pairs per annotator. Four senior bioinformaticians (two with PhDs) independently and concurrently scored every pair, blinded to model identity and judge score; annotators were recruited by open request on the basis of availability, without selection for prior experience with spatiAlytica. Each annotator scored every pair on two axes: a binary accept/reject decision (“Is the answer biologically correct and would you accept it?”) and a 5-point quality rubric (1 = incorrect/hallucinated, 5 = completely correct and complete). Because annotators assessed biological correctness directly rather than similarity to the LLM-generated reference, this validation is independent of the GPT-4o question-generation pipeline. Human ground truth for each pair was computed as the majority of four binary decisions (threshold ≥ 2*/*4 for acceptance) and the median of the four 5-point scores.

To test robustness of the headline to the choice of judge model family, we re-scored the same 100 pairs with two independent LLM judge families from two corporate entities (GPT-5.2, OpenAI; Claude Opus 4.6 high, Anthropic) under a reference-free protocol in which each judge sees only the image, the question, and the assistant’s answer, removing the GPT-4o reference from the judge loop entirely and mirroring the human-annotation protocol. Inter-annotator agreement (Krippendorff’s *α* on binary accept and 5-point rubric, pairwise Cohen’s *κ* across all rater pairs), judge-vs-human confusion matrices with conditional acceptance probabilities, the calibration-adjusted headline with 10,000-sample bootstrap 95% CI, per-question-type breakdown, per-judge-bin calibration curve, leave-one-rater-out sensitivity, per-judge pass rates with Wilson 95% CIs, pairwise Gwet’s AC1 across the two LLM judge families and the human majority vote, and per-question-category performance for all channels are reported in Supplementary Note 1.5 and summarized in Fig. 3e,f and the Results.

#### 3.10.5 Automated evaluation pipeline

We built a multi-stage evaluation pipeline that compares agent-generated outputs against precomputed ground-truth (GT) answers. Programmatic comparison takes precedence over language-model-based judgment. Three protocols were used. *Single-turn* (ST) presents each question independently and compares outputs programmatically. *Multi-turn* (MT) runs sequential conversations in which each turn builds on the `AnnData` state produced by the previous turn, allowing errors to cascade. *Teacher-forcing* (TF) replaces the agent’s state at each turn with the correct GT state, isolating per-turn accuracy from cascading failure. The comparison pipeline applies three tiers in order of decreasing determinism. 1) Targeted `AnnData` key extraction on the cumulative GT state, with structured state diffs that identify the keys modified by GT and by the agent. 2) Programmatic equality of return values dispatched by Python type: `None` checks; scalar and string equality with whitespace and code-fence normalization; `numpy.allclose` for dense arrays, with dense fallbacks for sparse matrices; and `pandas.testing.assert_frame_equal` or `assert_series_equal` for DataFrames and Series. 3) Type-specific relaxations for known stochastic or ambiguous outputs: PCA sign-flip canonicalization, UMAP and t-SNE embedding correlation, Adjusted Rand Index on cluster labels (threshold ≥ 0.9 for single-column cluster Series and ≥ 0.8 for multi-column clustering DataFrames), rank-genes-groups gene-name overlap on the `names`, `scores`, `pvals`, and `pvals_adj` columns, and proportion-versus-percentage rescaling when scalars or numeric Series differ by exactly 100 *×* . Outputs that fail every programmatic tier fall through to an LLM judge (GPT-5.2) [32]. The judge is called directly against the OpenAI API with `seed`=42 and a JSON-format response. The agent’s thirdparty API base, when in use, is never invoked for the judge. The judge prompt encodes an explicit rubric. Empty or `None` payloads on both sides are equivalent, and the case is short-circuited without an API call. Whitespace and incidental logging differences are ignored. A substantive-payload mismatch (one side contains tables or arrays the other lacks) is not equivalent. Ranked gene lists are equivalent at ≥ 80% overlap with medium confidence. A value expressed as a proportion (for example, 0.5) and the same value expressed as a percentage (for example, 50.0) are equivalent. Plot questions are scored separately by GPT-5.2 vision on a 1–5 scale on both code-similarity and rendered visual-similarity axes, with 3-attempt retries and a PDF-to-PNG fallback. The combined score 0.1 code+0.9 plot counts as a pass at ≥ 3. All evaluation subprocesses set fixed random seeds across Python `random`, NumPy, Scanpy, and `PYTHONHASHSEED` (seed = 42). The complete evaluation pipeline, including programmatic comparison tolerances, key-name recovery, structural alignment, and error classification, is described in Supplementary Note 1.6.

#### 3.10.6 Head-to-head comparison with BioMANIA and BioMedAgent

We performed controlled comparisons against two agentic biology baselines: BioMANIA [20], selected because it supports the same libraries (Scanpy, Squidpy), accepts conversational input, and includes execution retries (up to 5 per question); and BioMedAgent [23], a recent general-purpose bioinformatics multi-agent system organized into six specialized sub-agents (Linguist, Prompt Engineer, Tool Scorer, Workflow Designer, Programmer, Summary Analyst) included as a contextual single-turn baseline rather than a spatial-biology head-to-head peer. Both systems received the identical 222 single-turn questions, groundtruth code, and comparison pipeline as spatiAlytica; no study context is injected in any benchmark run (study-context prompts are used only in case studies). BioMANIA was also evaluated under multi-turn and teacher-forcing protocols on the 178-question spatiAlyticaBench-MT; BioMedAgent was evaluated under the single-turn protocol only. All four LLM configurations were routed through the same LiteLLM dispatch layer for both baselines to ensure identical model versions and temperature settings (*T* =0). Because BioMANIA operates through a multi-step conversational protocol, we automated its interaction via a two-tier controller (deterministic rule layer for execution confirmation and disambiguation, plus a lightweight LLM controller for parameter filling) that replicates the interactive confirmation step required by BioMANIA’s native UI; the LLM-controller overhead is negligible because most actions are handled by the rule layer. This automation represents an upper bound on BioMANIA performance. BioMANIA requires library-specific backends (Scanpy or Squidpy), so we evaluated each backend separately and report results stratified by library. The BioMedAgent Kimi-K2.5 configuration could not be reliably evaluated because the Together AI endpoint does not emit BioMedAgent’s required XML response tags, aborting the pipeline before code execution (Supplementary Note 1). Full per-mode, per-library BioMANIA results are reported in Supplementary Table 8 (single-turn) and Supplementary Table 13 (multi-turn); BioMedAgent single-turn results are reported in Supplementary Table 9; and a per-query compute and latency comparison for the Frontier configuration across all three systems is provided in Supplementary Table 11.

### 3.11 Reproducibility

spatiAlytica will be distributed on PyPI as `spatialytica-napari` (exposed to users as spatiAlytica); v0.3.5 was used for all results in this study. Source code, documentation, and benchmark data will be released at https://github.com/Huang-AI4Medicine-Lab/spatialytica-napari, with a Python 3.11 conda environment specification and pinned `requirements.txt` provided. The package is built around napari (v0.6.6) for the graphical interface, AnnData (v0.12.3) [28], Scanpy (v1.11.5) [49], and Squidpy (v1.8.1) [38] for spatialomics data structures and analyses, and LangChain (v0.3.27) [50] with LangGraph (v1.0.1) for subagent orchestration; provider-agnostic model routing is handled by LiteLLM (v1.78.7). Pinned versions for all dependencies that materially affect benchmark results are listed in Table 18, and a complete lockfile is provided in the repository. Language-model providers and per-agent decoding parameters are declared in a single `config.json` file; all benchmark results use the four configurations listed in Supplementary Table 6. We note two inherent limitations: LLM outputs are non-deterministic even with identical prompts, and model providers may update or deprecate versions. To mitigate these factors, we report all benchmark results as averages over multiple independent runs and archive exact model identifiers alongside all evaluation outputs. Each benchmark run was executed in complete isolation: distinct SLURM jobs with unique run identifiers, separate output directories for per-query logs and persisted memory, and fresh memory state initialized at the start of each session. No state was shared between runs or between evaluation conditions (ST, MT, TF, spatiAlyticaBench-ImageQA, ablations), ensuring that reported metrics reflect independent, clean-room execution rather than cumulative session state. Table 18 summarizes the key reproducibility parameters.

## Data availability

All benchmark datasets, evaluation scripts, prompts, per-query logs, judge outputs, and human-annotation artifacts will be released publicly, with a versioned snapshot archived on Zenodo. The spatiAlyticaBench-ST and spatiAlyticaBench-MT case lists were derived from the publicly hosted Scanpy and Squidpy tutorial datasets, converted manually to an AnnData h5ad format and will be archived on Zenodo (Scanpy clustering tutorial, https://scanpy.readthedocs.io/en/stable/tutorials/basics/clustering.html; Squidpy “Analysis of spatial datasets” tutorial collection, https://squidpy.readthedocs.io/en/latest/notebooks/tutorials/#analysis-of-spatial-datasets-using-squidpy). The case-study spatial transcriptomics datasets (Kaposi’s sarcoma, HGSOC, CRC CODEX) are previously published and publicly available; accessions and download instructions are provided in the original publications cited in the Methods. Per-dataset source links are summarized in Supplementary Table 2.

## Code availability

spatiAlytica will be available as an open-source Python package (`pip install spatialytica-napari`, version 0.3.5 at time of submission). The complete source code, benchmark scoring pipeline, statistical-analysis scripts, prompt templates, and cross-family LLM-judge runners (GPT-5.2, OpenAI; Claude Opus 4.6, Anthropic) will be released publicly at https://github.com/Huang-AI4Medicine-Lab/spatialytica-napari at acceptance, with the submission release archived on Zenodo. A pinned conda environment specification (Python 3.11), `requirements.txt`, and lockfile are distributed with the source release. All LLM calls were dispatched through LiteLLM against dated API snapshots: `gpt-5.2-2025-12-11`, `gpt-5-mini-2025-08-07`, `gpt-4.1-mini-2025-04-14`, and `together ai/moonshotai/Kimi-K2.5` (rolling alias; evaluation window 2026-03-01 to 2026-04-01). All stochastic operations are seeded (`random state=42`).

## Acknowledgements

This study was supported by grants from the National Institutes of Health (CA096512, CA284554, CA278812, CA291244, and CA124332 to S.-J.G.; U01CA279618 and R21GM155774 to Y.H.; R35GM154967 to Y.-C.C.), UPMC Hillman Cancer Center Startup Funds to S.-J.G. and Y.H., and in part by award P30CA047904. This study was also supported in part by the National Cancer Institute of the National Institutes of Health under Award Number P30CA047904 as Shark Tank funding to A.D. and Y.H. This study was also supported in part by the Commercialization Gap Fund Award by the University of Pittsburgh Office of Innovation and Entrepreneurship (OIE) to A.D. and Y.H. This research was also supported in part by the University of Pittsburgh Center for Research Computing, RRID:SCR 022735. Specifically, this work used the HTC cluster, which is supported by NIH award number S10OD028483.

## Author contributions

A.D. and Y.H. conceived and designed the study. A.D. developed the spatiAlytica system. A.D., K.Z., A.C., L.L., and Y.H. developed the evaluation framework. J.S. performed data collection for the spatiAlyticaBench-ImageQA dataset. M.H. performed data collection for spatiAlyticaBench-ST. A.D. created the spatiAlyticaBench-MT dataset. H.G., A.D., and Y.H. developed the CAF remodeling case study. W.M., A.D., S.-J.G., and Y.H. developed the Kaposi’s sarcoma case studies. P.C., Y.-C.C., A.D., and Y.H. developed the HGSOC case study. S.J., Z.L., M.M.H., A.O., and H.S. assisted with evaluation. Y.H. supervised the study. All authors contributed to writing and approved the final manuscript.

## Supplementary information

Supplementary Figures S1–S4, Supplementary Tables 1–18, Supplementary Notes 1–9.

## References

[1] Chen, K. H., Boettiger, A. N., Moffitt, J. R., Wang, S. & Zhuang, X. Spatially resolved, highly multiplexed RNA profiling in single cells. Science 348, aaa6090 (2015).

[2] Janesick, A. et al. High resolution mapping of the tumor microenvironment using integrated single-cell, spatial and in situ analysis. Nature Communications 14, 8353 (2023).

[3] Eng, C.-H. L. et al. Transcriptome-scale super-resolved imaging in tissues by RNA seqFISH+. Nature 568, 235–239 (2019).

[4] Wang, X. et al. Three-dimensional intact-tissue sequencing of single-cell transcriptional states. Science 361, eaat5691 (2018).

[5] Goltsev, Y. et al. Deep profiling of mouse splenic architecture with CODEX multiplexed imaging. Cell 174, 968–981 (2018).

[6] Giesen, C. et al. Highly multiplexed imaging of tumor tissues with subcellular resolution by mass cytometry. Nature Methods 11, 417–422 (2014).

[7] Gut, G., Herrmann, M. D. & Pelkmans, L. Multiplexed protein maps link subcellular organization to cellular states. Science 361, eaar7042 (2018).

[8] He, S. et al. High-plex imaging of RNA and proteins at subcellular resolution in fixed tissue by spatial molecular imaging. Nature Biotechnology 40, 1794–1806 (2022).

[9] Stahl, P. L. et al. Visualization and analysis of gene expression in tissue sections by spatial transcriptomics. Science 353, 78–82 (2016).

[10] Rodriques, S. G. et al. Slide-seq: a scalable technology for measuring genome-wide expression at high spatial resolution. Science 363, 1463–1467 (2019).

[11] Stickels, R. R. et al. Highly sensitive spatial transcriptomics at near-cellular resolution with Slide-seqV2. Nature Biotechnology 39, 313–319 (2021).

[12] Marx, V. Method of the Year: spatially resolved transcriptomics. Nature Methods 18, 9–14 (2021).

[13] Moses, L. & Pachter, L. Museum of spatial transcriptomics. Nature Methods 19, 534–546 (2022).

[14] Rao, A., Barkley, D., Franca, G. S. & Yanai, I. Exploring tissue architecture using spatial transcriptomics. Nature 596, 211–220 (2021).

[15] Lewis, S. M. et al. Spatial omics and multiplexed imaging to explore cancer biology. Nature Methods 18, 997–1012 (2021).

[16] Vandereyken, K., Sifrim, A., Thienpont, B. & Voet, T. Methods and applications for single-cell and spatial multi-omics. Nature Reviews Genetics 24, 494–515 (2023).

[17] Longo, S. K., Guo, M. G., Ji, A. L. & Khavari, P. A. Integrating single-cell and spatial transcriptomics to elucidate intercellular tissue dynamics. Nature Reviews Genetics 22, 627–644 (2021).

[18] OpenAI. GPT-4 technical report (2023). arXiv:2303.08774.

[19] Zhou, J. et al. An AI Agent for Fully Automated Multi-Omic Analyses. Advanced Science 11, 2407094 (2024). URL https://onlinelibrary.wiley.com/doi/abs/10.1002/advs.202407094. eprint: https://onlinelibrary.wiley.com/doi/pdf/10.1002/advs.202407094.

[20] Dong, Z., Zhou, H., Jiang, Y., Zhong, V. & Lu, Y. Y. Simplifying bioinformatics data analysis through conversation (2024). URL https://www.biorxiv.org/content/10.1101/2023.10.29.564479v2. Pages: 2023.10.29.564479 Section: New Results.

[21] Xiao, Y. et al. CellAgent: An LLM-driven Multi-Agent Framework for Automated Single-cell Data Analysis (2024). URL http://arxiv.org/abs/2407.09811. ArXiv:2407.09811 [cs].

[22] Wang, H. et al. SpatialAgent: An Autonomous AI Agent for Spatial Biology (2025). URL https://www.biorxiv.org/content/10.1101/2025.04.03.646459v1. Pages: 2025.04.03.646459 Section: New Results.

[23] Bu, D. et al. Empowering AI data scientists using a multi-agent LLM framework with self-evolving capabilities for autonomous, tool-aware biomedical data analyses. Nature Biomedical Engineering (2026). URL https://www.nature.com/articles/s41551-026-01634-6.

[24] Chiu, H.-H. et al. scChat: A large language model-powered co-pilot for contextualized single-cell RNA sequencing analysis. AIChE Journal e70285 (2026).

[25] Lubiana, T. et al. Ten quick tips for harnessing the power of ChatGPT in computational biology. PLOS Computational Biology 19, e1011319 (2023).

[26] Gao, S. et al. Empowering biomedical discovery with AI agents. Cell 187, 6125–6151 (2024).

[27] napari contributors. napari: a multi-dimensional image viewer for Python (2019). URL https://napari.org.

[28] Virshup, I., Rybakov, S., Theis, F. J., Angerer, P. & Wolf, F. A. anndata: Access and store annotated data matrices. Journal of Open Source Software 9, 4371 (2024).

[29] Wang, X. et al. MINT: Evaluating LLMs in multi-turn interaction with tools and language feedback. The Twelfth International Conference on Learning Representations (ICLR) (2024).

[30] Song, J., Das, A., Cui, G. & Huang, Y. Figex: Aligned extraction of scientific figures and captions. Findings of the Association for Computational Linguistics: EMNLP 2025 (2025).

[31] Li, C. et al. Llava-med: Training a large language-and-vision assistant for biomedicine in one day. Advances in Neural Information Processing Systems 36, 28541–28564 (2023).

[32] Zheng, L. et al. Judging LLM-as-a-judge with MT-Bench and Chatbot Arena. Advances in Neural Information Processing Systems 36 (2023).

[33] Chaves, J. M. Z. et al. Towards a clinically accessible radiology foundation model: open-access and lightweight, with automated evaluation. arXiv preprint arXiv:2403.08002 (2024).

[34] Cesarman, E. et al. Kaposi sarcoma. Nature Reviews Disease Primers 5, 9 (2019).

[35] Mesri, E. A., Cesarman, E. & Boshoff, C. Kaposi’s sarcoma and its associated herpesvirus. Nature Reviews Cancer 10, 707–719 (2010).

[36] Meng, W. et al. Spatial single-cell atlas reveals kshv-driven broad cellular reprogramming, progenitor expansion, immune and vascular remodeling in kaposi’s sarcoma. bioRxiv 2025–09 (2025).

[37] Schurch, C. M. et al. Coordinated cellular neighborhoods orchestrate antitumoral immunity at the colorectal cancer invasive front. Cell 182, 1341–1359 (2020).

[38] Palla, G. et al. Squidpy: a scalable framework for spatial omics analysis. Nature Methods 19, 171–178 (2022).

[39] Rauch, D. A. et al. Single-cell transcriptomic analysis of kaposi sarcoma. PLoS pathogens 21, e1012233 (2025).

[40] Di Caro, G. et al. Occurrence of tertiary lymphoid tissue is associated with t-cell infiltration and predicts better prognosis in early-stage colorectal cancers. Clinical Cancer Research 20, 2147–2158 (2014).

[41] Helmink, B. A. et al. B cells and tertiary lymphoid structures promote immunotherapy response. Nature 577, 549–555 (2020).

[42] Sautes-Fridman, C., Petitprez, F., Calderaro, J. & Fridman, W. H. Tertiary lymphoid structures in the era of cancer immunotherapy. Nature Reviews Cancer 19, 307–325 (2019).

[43] Danenberg, E. et al. Breast tumor microenvironment structures are associated with genomic features and clinical outcome. Nature Genetics 54, 660–669 (2022).

[44] Licaj, M. et al. Residual antxr1+ myofibroblasts after chemotherapy inhibit anti-tumor immunity via yap1 signaling pathway. Nature Communications 15, 1312 (2024).

[45] Dholakia, K., Jaff, I. & BerriAI Contributors. LiteLLM: Call all LLM APIs using the OpenAI format. https://github.com/BerriAI/litellm (2023). Version 1.78.7.

[46] Erman, L. D., Hayes-Roth, F., Lesser, V. R. & Reddy, D. R. The Hearsay-II speech-understanding system: Integrating knowledge to resolve uncertainty. ACM Computing Surveys 12, 213–253 (1980).

[47] Yao, S. et al. ReAct: Synergizing reasoning and acting in language models (2023). International Conference on Learning Representations, arXiv:2210.03629.

[48] Schick, T. et al. Toolformer: Language models can teach themselves to use tools. Advances in Neural Information Processing Systems 36, 68539–68551 (2023).

[49] Wolf, F. A., Angerer, P. & Theis, F. J. SCANPY: large-scale single-cell gene expression data analysis. Genome Biology 19, 15 (2018).

[50] Chase, H. LangChain (2022). URL https://github.com/langchain-ai/langchain.

