## Supplementary Fig. 1 for "spatiAlytica: Viewer-Grounded Multimodal Agentic System for Interactive Spatial Omics Analysis"

### Supplementary Figures

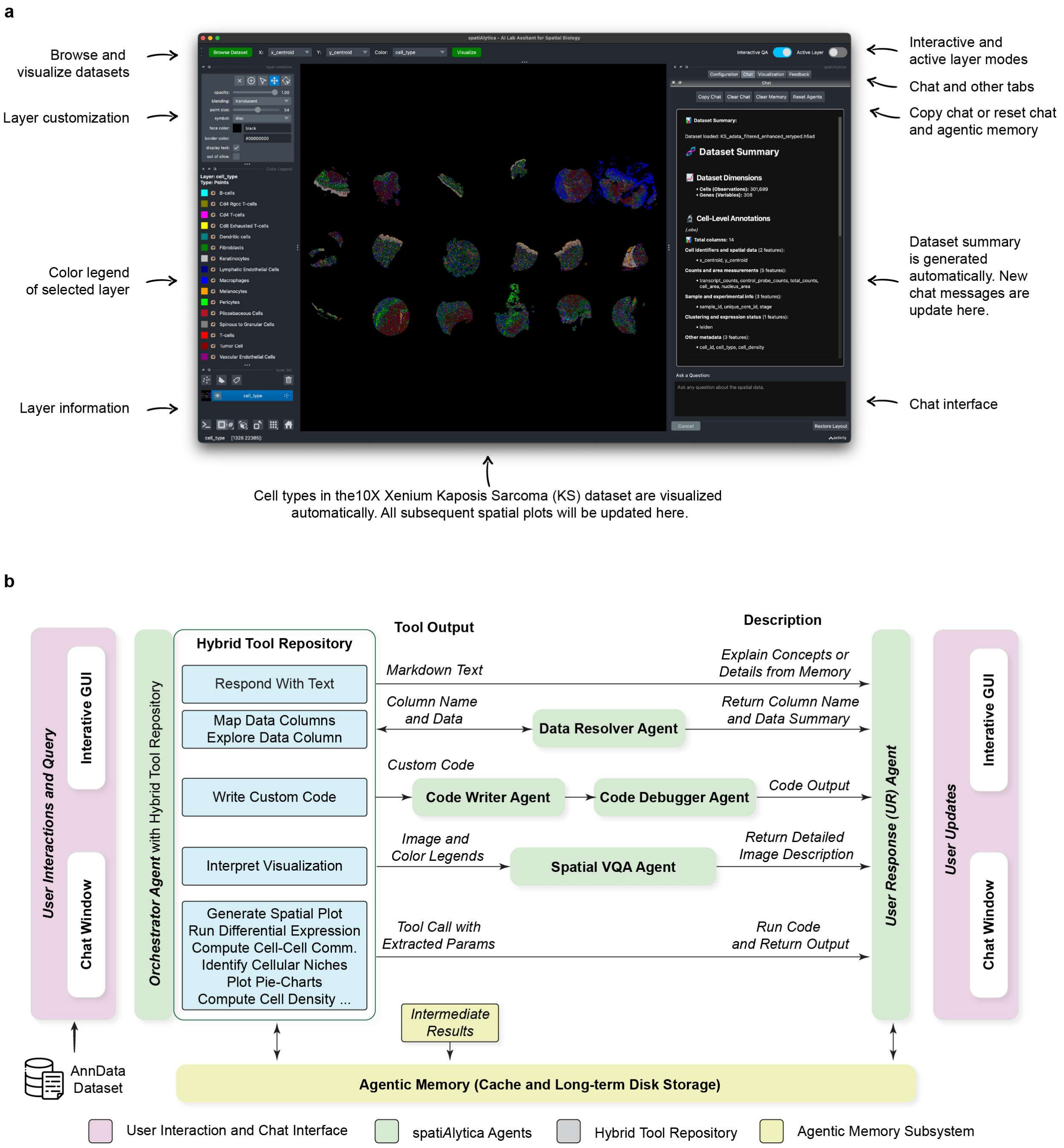


**Supplementary Figure 1 spatiAlytica user interface and multimodal interactive agentic architecture.** (A) Screenshot of the spatiAlytica Napari plugin loaded with the Xenium Kaposi's sarcoma dataset, showing the spatial viewer, color legend, auto-generated dataset summary, and chat interface. (B) Schematic of the multimodal interactive agentic system. The orchestrator sub-agent routes queries to specialized sub-agents (data resolver, code writer, code debugger, spatial VQA) or direct tool calls. All sub-agents communicate via the agentic memory sub-system; outputs are returned to the user through the user response sub-agent.
