## Supplementary Fig. 2 for "spatiAlytica: Viewer-Grounded Multimodal Agentic System for Interactive Spatial Omics Analysis"

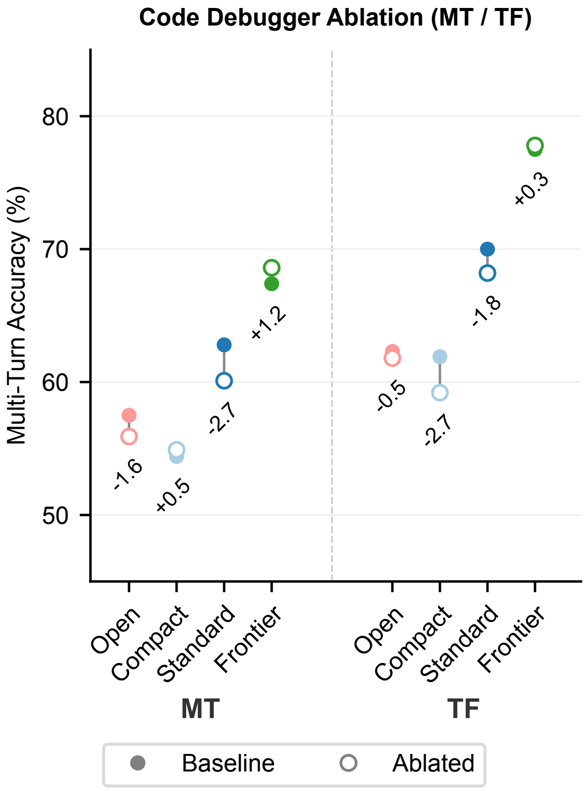


**Supplementary Figure 2  Code debugger (error correction) ablation in multi-turn and teacher-forcing modes.** Vertical dumbbell plots showing baseline (filled) versus no-EC ablated (open) accuracy for each LLM configuration in MT (left) and TF (right) modes. Deltas are small in magnitude and mixed in direction (MT: −2.7 to +1.2 pp; TF: −2.7 to +0.3 pp), indicating that the code debugger sub-agent's contribution is concentrated in single-turn execution (Fig. 3b) rather than multi-turn modes where subsequent context can compensate for uncorrected failures.
