## Supplementary Fig. 3 for "spatiAlytica: Viewer-Grounded Multimodal Agentic System for Interactive Spatial Omics Analysis"

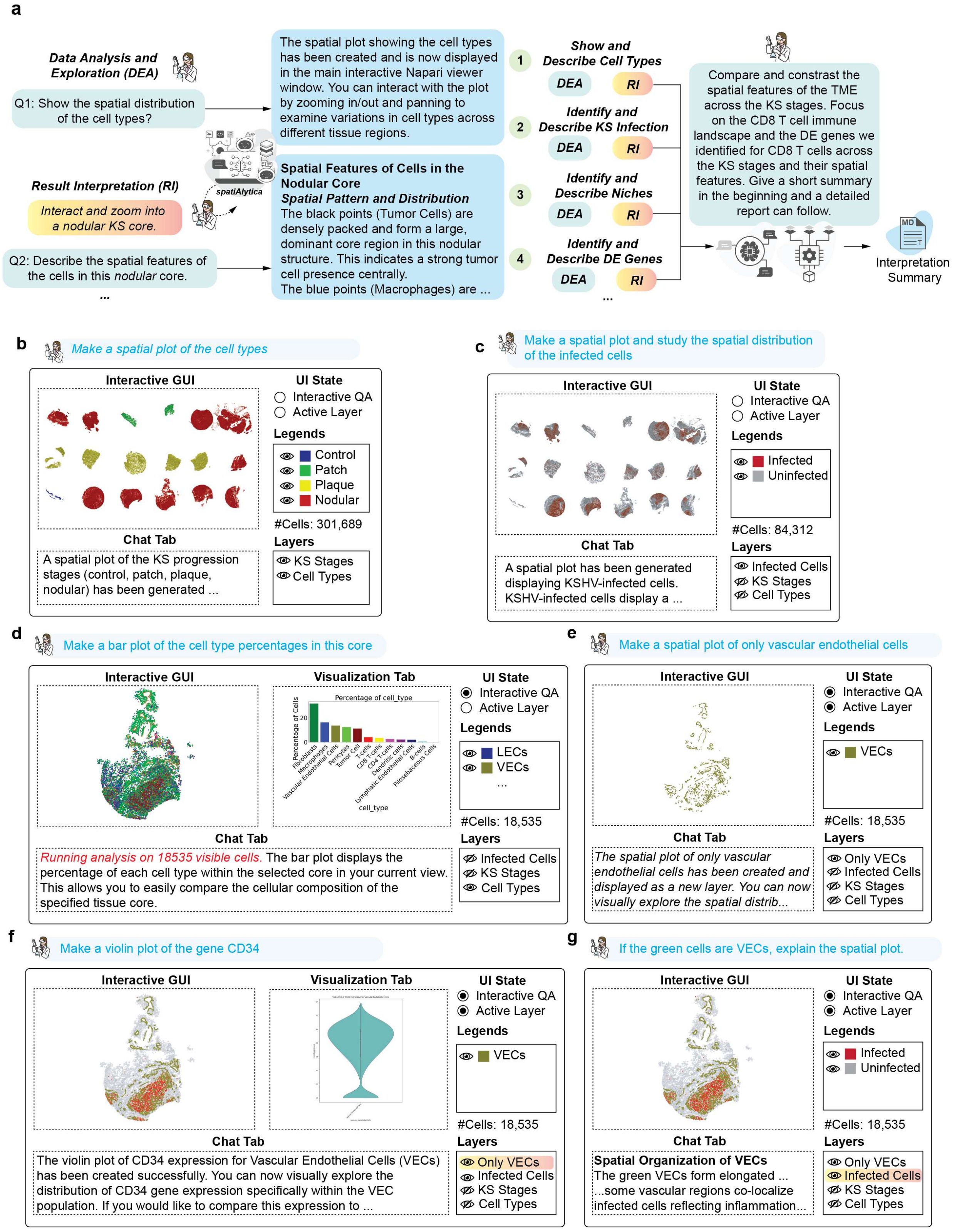


**Supplementary Figure 3 Representative QA pairs from the spatiAlyticaBench-ImageQA human-validation subsample.** Four example pairs drawn from the stratified $N=100$ human-validation subsample illustrate the question-type taxonomy used for per-category reporting and the distractor attention checks interspersed through the annotation task (Methods, Sec. 3.10.4). (A) *Factual* (`pair_7f3b139c39`): question probes direct readout of surface-marker expression from flow-cytometry plots tracking BM monocyte to oTME macrophage differentiation (Ly6C$^{high}$F4/80$^{neg}$ to F4/80$^{+}$ transition). (B) *Interpretive* (`pair_460ee1f176`): question asks the model to explain what a hexagonally-binned IBSP spatial-transcriptomics heat map represents, requiring synthesis of visual encoding (color scale, spatial layout) and biological meaning. (C) *Spatial* (`pair_7b00045591`): question probes identification of anatomical features (trabecular myocardium) from the spatial distribution of Leiden clusters across four developmental stages (P5, P9, P16, P23). (D) *Distractor* (`pair_f40f52162c`, `is_distractor = True`): the answer text has been deliberately swapped with the ground-truth response from an unrelated image-question pair (B/plasma/T/macrophage color coding around glomeruli and B-cell/PC foci), producing an obvious content mismatch with the myoneme-number box plot shown. Nine such distractor pairs were interspersed per rater as attention checks; raters whose distractor accept rate exceeded chance were flagged for exclusion. Panels reproduce the exact rendering shown to human annotators in the blinded scoring interface, including pair identifiers. Ground-truth / model answers shown beneath each image are the LLM-generated references against which the GPT-5.2 judge scored candidate answers.
