## Supplementary Fig. 4 for "spatiAlytica: Viewer-Grounded Multimodal Agentic System for Interactive Spatial Omics Analysis"

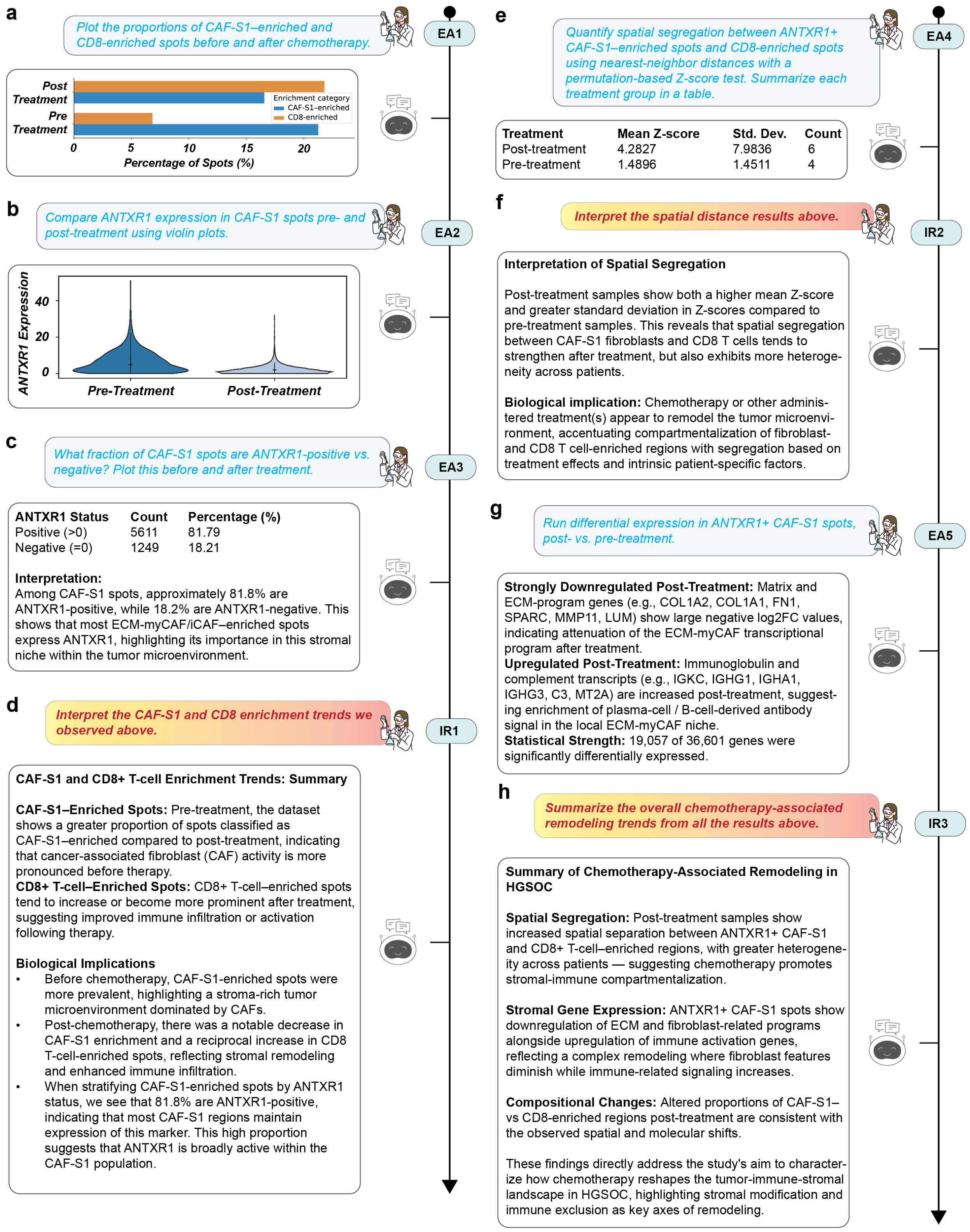


**Supplementary Figure 4 Agentic analysis of chemotherapy-driven stromal and immune remodeling in high-grade serous ovarian cancer.** (A) Proportions of CAF-S1-enriched and CD8-enriched spots before and after chemotherapy, showing pre-treatment CAF-S1 predominance and post-treatment CD8 expansion (EA1). (B) Violin plots of *ANTXR1* expression in CAF-S1 spots revealing a post-treatment decrease consistent with attenuation of the ECM-myCAF phenotype (EA2). (C) *ANTXR1*-positive versus *ANTXR1*-negative fractions among CAF-S1 spots, showing the ECM-myCAF ($ANTXR1^+$) share dropping from 90.9% pre-chemo to 82.5% post-chemo (EA3). (D) Interpretive synthesis linking pre-treatment CAF-S1 predominance and post-treatment CD8 expansion to stromal remodeling and enhanced immune infiltration (IR1). (E) Permutation-based neighborhood-enrichment between $ANTXR1^+$ CAF-S1 (ECM-myCAF) and CD8-enriched spots, with the cross-pair $Z$-score becoming more negative post-chemotherapy ($-2.05 \to -5.63$), indicating stronger spatial avoidance (EA4). (F) Interpretation of spatial distance results indicating chemotherapy strengthens spatial segregation between fibroblast and $CD8^+$ T-cell regions (IR2). (G) Differential expression in ECM-myCAF ($ANTXR1^+$ CAF-S1) spots, post-chemotherapy versus pre-chemotherapy, showing downregulation of matrix and ECM-program genes (*COL1A2*, *COL1A1*, *FN1*, *SPARC*, *MMP11*, *LUM*) and upregulation of immunoglobulin and complement transcripts (*IGKC*, *IGHG1*, *IGHA1*, *C3*) (EA5). (H) Integrated summary of chemotherapy-associated remodeling describing spatial segregation, transcriptional reprogramming, and compositional shifts consistent with stromal-immune remodeling (IR3).
