## Supplementary Table 1 for "spatiAlytica: Viewer-Grounded Multimodal Agentic System for Interactive Spatial Omics Analysis"

**Supplementary Table 1** **Feature comparison of spatiAlytica with existing spatial biology and LLM analysis tools. ✓ = supported; ○ = partial; × = not supported; – = not applicable.**

| Capability | spatiAlytica | STAgent | SpatialAgent | BioMANIA | CellAgent | Xenium Explorer | ChatGPT + Jupyter |
|---|---|---|---|---|---|---|---|
| NL query interface | ✓ Napari chat | ✓ Streamlit | ✓ CLI/script | ✓ Web | ✓ Web | × | ✓ Data-blind |
| Interactive spatial visualization | ✓ Napari multi-layer | ○ Static plots | ○ Static plots | × | × | ✓ WebGL | × Matplotlib |
| Data-aware agents | ✓ AnnData structure | ✓ Reads data | ✓ Reads data | ○ Package selection | ○ User describes data | × Manual | × Must paste |
| Spatial analysis tools | ✓ Niche, Moran's $I$, LISA, DE | ✓ Squidpy/Scanpy | ✓ 72 tools | × Package-bound | × scRNA-seq only | ○ Built-in, limited | × Hallucinates |
| Error self-correction | ✓ 3 retries | ○ Code retry | ○ Adaptive | ✓ | ✓ | – | × |
| Persistent memory | ✓ LangChain | × | ○ Short/long-term | × | × | × | × |
| Column/entity resolution | ✓ Fuzzy matching, data resolver agent | × | ○ CZIInfo tool | × | × | × | × |
| Literature integration | ✓ PubMed + S2 | ✓ Retrieval | ○ DB query | × | × | × | ○ Training data |
| Evidence tracking | ✓ Structured records | ○ Report output | × | × | × | × | × |
| Biological interpretation | ✓ Grounded sub-agent synthesis | ✓ LLM synthesis | ✓ LLM-driven | × | ○ | × | ✓ |
