## Supplementary Table 2 for "spatiAlytica: Viewer-Grounded Multimodal Agentic System for Interactive Spatial Omics Analysis"

**Supplementary Table 2 Datasets used for creating spatiAlyticaBench** Each dataset was used for both single-turn (ST) and multi-turn (MT/TF) evaluation where applicable. Cell/spot counts and gene panel sizes reflect the processed data used in evaluation. The *Source* column provides the verified upstream URL for each dataset: spatial-platform datasets are hosted as part of the Squidpy tutorial collection, the scRNA-seq references are hosted as part of the Scanpy core plotting, preprocessing-and-clustering, and trajectory-inference tutorials, and the Kaposi's sarcoma case-study dataset is from Meng et al. [36].

| Dataset | Platform | Tissue | Resolution | Cells/Spots | Genes | ST Qs | MT turns | Source |
|---|---|---|---|---|---|---|---|---|
| IMC | Imaging Mass Cytometry | Breast cancer | Subcellular | 4,668 | 34 | 10 | 16 | Squidpy tut. |
| CorePlot | scRNA-seq (10x) | PBMC | Single-cell | 700 | 765 | 36 | 14 | Scanpy tut. |
| PrepClus | scRNA-seq (10x) | Bone marrow (BMMC) | Single-cell | 17,041 | 23,427 | 42 | 13 | Scanpy tut. |
| 4i | 4i immunofluorescence | HeLa cells | Subcellular | 270,876 | 43 | 9 | 15 | Squidpy tut. |
| KS | 10x Xenium | Skin | Subcellular | 301,689 | 308 | 24 | 15 | Ref. [36] |
| MERFISH | MERFISH | Mouse brain | Subcellular | 73,655 | 161 | 11 | 15 | Squidpy tut. |
| Nanostring | NanoString CosMx | Liver | Subcellular | 87,243 | 980 | 27 | 21 | Squidpy tut. |
| SlideSeq2 | Slide-seqV2 | Mouse cerebellum | Spot-level | 41,786 | 4,000 | 12 | 17 | Squidpy tut. |
| Xenium | 10x Xenium | Breast cancer | Subcellular | 154,472 | 377 | 24 | 18 | Squidpy tut. |
| SeqFish | seqFISH | Mouse brain | Subcellular | 19,416 | 351 | 8 | 17 | Squidpy tut. |
| TrajInf | scRNA-seq (reference) | Mouse hematopoiesis | Single-cell | 2,730 | 3,451 | 19 | 17 | Scanpy tut. |
| **Total** | | | | | | **222** | **178** | |
