## Supplementary Table 3 for "spatiAlytica: Viewer-Grounded Multimodal Agentic System for Interactive Spatial Omics Analysis"

**Supplementary Table 3 Task distribution across spatiAlyticaBench categories (222 single-turn + 178 multi-turn = 400 total).** Each question produces either a data output (numeric result, Series, DataFrame) or a plot output (saved figure). Questions span 11 datasets from 7 spatial platforms and 3 scRNA-seq references.

| Task Category | Data | Plot | Total |
|---|---|---|---|
| Spatial Analysis | 61 | 34 | 95 |
| Visualization | 0 | 66 | 66 |
| Memory Retrieval | 53 | 1 | 54 |
| Cell Composition & Statistics | 44 | 1 | 45 |
| Quality Control & Preprocessing | 30 | 16 | 46 |
| Differential Expression | 7 | 16 | 23 |
| Gene Expression Analysis | 10 | 1 | 11 |
| Niche / Neighborhood Analysis | 15 | 3 | 18 |
| Trajectory / Pseudotime | 6 | 11 | 17 |
| Clustering & Cell Annotation | 13 | 1 | 14 |
| Dimensionality Reduction | 11 | 0 | 11 |
| **Total** | **250** | **150** | **400** |
