## Supplementary Table 4 for "spatiAlytica: Viewer-Grounded Multimodal Agentic System for Interactive Spatial Omics Analysis"

**Supplementary Table 4 spatiAlyticaBench-ST per-platform task distribution (222 single-turn questions).** Rows are spatial omics platforms (and a combined scRNA-seq reference row covering CorePlot, PrepClus, and TrajInf); columns are the 10 ST task categories using compact labels (Spatial: Spatial Analysis; QC: Quality Control and Preprocessing; Viz: Visualization; DE: Differential Expression; Niche: Niche / Neighborhood Analysis; Traj: Trajectory / Pseudotime; DimRed: Dimensionality Reduction; Clust: Clustering and Cell-type Annotation; Comp: Cell Composition and Statistics; GeneE: Gene Expression Analysis). Cell values are question counts; 0 indicates the category is not exercised on that platform.

| **Platform** | **Spatial** | **QC** | **Viz** | **DE** | **Niche** | **Traj** | **DimRed** | **Clust** | **Comp** | **GeneE** | **Total** |
|---|---|---|---|---|---|---|---|---|---|---|---|
| IMC | 10 | 0 | 0 | 0 | 0 | 0 | 0 | 0 | 0 | 0 | 10 |
| 4i | 9 | 0 | 0 | 0 | 0 | 0 | 0 | 0 | 0 | 0 | 9 |
| MERFISH | 8 | 0 | 0 | 2 | 0 | 0 | 0 | 0 | 0 | 1 | 11 |
| CosMx SMI | 13 | 9 | 1 | 0 | 0 | 0 | 3 | 1 | 0 | 0 | 27 |
| Slide-seqV2 | 10 | 0 | 0 | 0 | 2 | 0 | 0 | 0 | 0 | 0 | 12 |
| 10x Xenium | 11 | 8 | 1 | 2 | 14 | 0 | 3 | 2 | 4 | 3 | 48 |
| seqFISH | 6 | 0 | 0 | 0 | 2 | 0 | 0 | 0 | 0 | 0 | 8 |
| scRNA-seq references | 0 | 25 | 27 | 17 | 0 | 16 | 5 | 7 | 0 | 0 | 97 |
| **Total** | **67** | **42** | **29** | **21** | **18** | **16** | **11** | **10** | **4** | **4** | **222** |
