## Supplementary Table 5 for "spatiAlytica: Viewer-Grounded Multimodal Agentic System for Interactive Spatial Omics Analysis"

**Supplementary Table 5 spatiAlyticaBench-MT per-conversation memory-dependence breakdown (178 turns across 11 multi-hop conversations).** For each conversation: total turns and counts under the four-way memory-dependence taxonomy (anaphoric continuation, implicit pronoun resolution, multi-result synthesis, and independent / seed turns). The first three categories collectively constitute the memory-dependent turns (124 total, 69.7%); the multi-result-synthesis bucket coincides with the dedicated *Memory Retrieval* task category. Classifications were assigned by manual review of every turn against the taxonomy definitions in §1.2

| Dataset | Platform | Turns | Anaph. | Pronoun | Multi-res. | Indep. | Mem-dep. |
|---|---|---|---|---|---|---|---|
| IMC | IMC | 16 | 0 | 7 | 4 | 5 | 11 |
| CorePlot | scRNA-seq | 14 | 2 | 4 | 4 | 4 | 10 |
| PrepClus | scRNA-seq | 13 | 2 | 5 | 3 | 3 | 10 |
| 4i | 4i | 15 | 1 | 4 | 5 | 5 | 10 |
| KS | 10x Xenium | 15 | 2 | 8 | 3 | 2 | 13 |
| MERFISH | MERFISH | 15 | 0 | 7 | 5 | 3 | 12 |
| Nanostring | CosMx SMI | 21 | 2 | 4 | 6 | 9 | 12 |
| SlideSeq2 | Slide-seqV2 | 17 | 0 | 6 | 6 | 5 | 12 |
| Xenium | 10x Xenium | 18 | 1 | 5 | 5 | 7 | 11 |
| SeqFish | seqFISH | 17 | 0 | 7 | 5 | 5 | 12 |
| TrajInf | scRNA-seq | 17 | 2 | 1 | 8 | 6 | 11 |
| **Total** | | **178** | **12** | **58** | **54** | **54** | **124** |
