## Supplementary Table 6 for "spatiAlytica: Viewer-Grounded Multimodal Agentic System for Interactive Spatial Omics Analysis"

**Supplementary Table 6 Model configurations used for benchmark evaluation.** Each configuration specifies the LLM assigned to each agent role. The three proprietary configurations share GPT-4.1-mini for sub-agents (user response sub-agent, data resolver sub-agent, and memory sub-routine), upgrading only the orchestrator and code debugger sub-agents. The open-source configuration uses Kimi-K2.5 for all roles. Dated API snapshots used for all runs: `gpt-5.2-2025-12-11`, `gpt-5-mini-2025-08-07`, `gpt-4.1-mini-2025-04-14`, and `together_ai/moonshotai/Kimi-K2.5` (rolling alias; Together AI exposes no dated snapshot, so Open-source numbers reflect the alias state during the evaluation window, 2026-03-01 to 2026-04-01).

| Sub-agent | Open-source | Compact | Standard | Frontier |
|---|---|---|---|---|
| Orchestrator | Kimi-K2.5 | GPT-4.1-mini | GPT-5-mini | GPT-5.2 |
| Code writer | Kimi-K2.5 | GPT-4.1-mini | GPT-5-mini | GPT-5.2 |
| Code debugger | Kimi-K2.5 | GPT-4.1-mini | GPT-5-mini | GPT-5.2 |
| Spatial VQA | Kimi-K2.5 | GPT-4.1-mini | GPT-5-mini | GPT-5.2 |
| Data resolver | Kimi-K2.5 | GPT-4.1-mini | GPT-4.1-mini | GPT-4.1-mini |
| User response | Kimi-K2.5 | GPT-4.1-mini | GPT-4.1-mini | GPT-4.1-mini |
| Memory sub-system | Kimi-K2.5 | GPT-4.1-mini | GPT-4.1-mini | GPT-4.1-mini |
| API provider | Together AI | OpenAI | OpenAI | OpenAI |
| License | Open-source | Proprietary | Proprietary | Proprietary |

In the codebase, the orchestrator sub-agent, code writer sub-agent, and spatial VQA sub-agent share a single configurable LLM slot; the code debugger sub-agent has its own slot but is paired with the same model as the shared slot in all four configurations reported here. The memory sub-routine performs conversation summarization and is not itself a core sub-agent.
