## Supplementary Table 7 for "spatiAlytica: Viewer-Grounded Multimodal Agentic System for Interactive Spatial Omics Analysis"

**Supplementary Table 7 Benchmark summary with 95% Wilson confidence intervals for spatiAlytica.** Mean accuracy (%) is macro-averaged across datasets; Wilson CIs are computed from pooled per-question pass/fail counts across $N$ independent runs. †: cross-run s.d. $> 3$ pp, indicating notable run-to-run variability. BioMANIA results are reported separately in Supplementary Tables 8 and 13.

| **Configuration** | **ST** ($N$=3) | **MT** ($N$=3) | **TF** ($N$=3) | **spatiAlyticaBench-ImageQA** ($N$=2) |
|---|---|---|---|---|
| Frontier (GPT-5.2) | 83.5 [80.5, 86.1] | 65.7 [61.8, 69.4] | 71.0 [67.2, 74.5] | 86.0 [85.3, 86.5] |
| Standard (GPT-5-mini) | 73.1 [69.6, 76.3]† | 61.1 [57.1, 64.9] | 63.1 [59.1, 66.9] | 25.7 [25.0, 26.4] |
| Compact (GPT-4.1-mini) | 69.5 [65.9, 72.9] | 56.7 [52.7, 60.6] | 56.8 [52.8, 60.7] | 81.1 [80.5, 81.7] |
| Open-source (Kimi-K2.5) | 54.7 [50.9, 58.4] | 51.7 [47.7, 55.7] | 56.6 [52.6, 60.5] | 26.8 [26.1, 27.5] |

Wilson CIs are computed from pooled correct/total counts across all runs. Pooled proportions may differ slightly from macro-averaged means. spatiAlyticaBench-ImageQA CIs are computed from $N$=2 runs $\times$ 7,350 QA pairs per run.
