## Supplementary Table 8 for "spatiAlytica: Viewer-Grounded Multimodal Agentic System for Interactive Spatial Omics Analysis"

**Supplementary Table 8 BioMANIA single-turn (ST) benchmark performance split by evaluation library (mean ± s.d., $N$=3 independent runs).** The benchmark comprises 222 independent questions across 11 spatial omics datasets; each question is evaluated twice (once with scanpy and once with squidpy). BioMANIA runs use the identical questions, ground-truth code, and comparison pipeline as spatiAlytica, with all four LLM configurations routed through the same LiteLLM dispatch layer (temperature $T$=0). Best accuracy per library is shown in **bold**.

| | **BioMANIA Agent Configuration** | | | |
|---|---|---|---|---|
| **Metric** | **Open-source (Kimi-K2.5)** | **Compact (GPT-4.1-mini)** | **Standard (GPT-5-mini)** | **Frontier (GPT-5.2)** |
| *Scanpy* | | | | |
| ST accuracy (%) | 42.9 ± 0.5 | 42.6 ± 1.4 | **43.8 ± 0.7** | 43.8 ± 0.5 |
| *Squidpy* | | | | |
| ST accuracy (%) | **37.2 ± 2.9** | 23.3 ± 2.3 | 25.5 ± 6.0 † | 26.3 ± 3.4 † |

†: cross-run s.d. > 3 pp. Scanpy-library performance clusters within ∼1 pp across all four backends (42.6–43.8%); squidpy-library performance is substantially lower and more variable for the three GPT backends (23.3–26.3%), driven by execution errors when the scanpy-native datasets are dispatched through squidpy (non-equivalent outputs counted as incorrect). Per-backend macro-averaged ST gaps (spatiAlytica − BioMANIA, weighted 6 scanpy + 5 squidpy): Kimi-K2.5 14.4 pp, Compact 35.7 pp, Standard 37.6 pp, Frontier 47.6 pp.
