## Supplementary Table 9 for "spatiAlytica: Viewer-Grounded Multimodal Agentic System for Interactive Spatial Omics Analysis"

**Supplementary Table 9 BioMedAgent performance on spatiAlyticaBench-ST across four model configurations (mean ± s.d., $N$=3 independent runs).** Accuracy is the fraction of 222 questions judged equivalent to ground truth. Tokens and cost are per-query averages. Best accuracy is shown in **bold**.

| Metric | BioMedAgent Configuration | | | |
|---|---|---|---|---|
| | **Open-source (Kimi-K2.5)** | **Compact (GPT-4.1-mini)** | **Standard (GPT-5-mini)** | **Frontier (GPT-5.2)** |
| Run 1 accuracy (%) | 0.0 | 33.3 | 7.7 | 33.8 |
| Run 2 accuracy (%) | 0.0 | 29.7 | 13.5 | 32.4 |
| Run 3 accuracy (%) | 0.0 | 31.1 | 18.5 | 33.8 |
| Overall accuracy (%) | $0.0 \pm 0.0^{*}$ | $31.4 \pm 1.8$ | $13.2 \pm 5.4$ † | $\mathbf{33.3 \pm 0.8}$ |
| Avg. prompt tok/query | 12,387 | 100,975 | 127,286 | 87,348 |
| Avg. completion tok/query | 16,581 | 10,324 | 42,843 | 14,920 |
| Avg. total tok/query | 28,968 | 111,299 | 170,130 | 102,267 |
| Avg. cost/query ($) | 0.004 | 0.021 | 0.119 | 1.321 |

†: cross-run s.d. > 3 pp. *Kimi-K2.5 is incompatible with BioMedAgent's strict XML-tag response protocol (e.g. `<STAGE>`, `<CODE>`): the Together AI endpoint does not reliably emit the required tags, triggering BioMedAgent's response validator to abort the multi-agent pipeline before code execution. A runtime lenient-tag patch restored pipeline flow but produced semantically empty outputs (0 code executions logged), yielding 0% across all runs; this row is shown for completeness and is not a fair measurement of Kimi-K2.5's reasoning capability. The BioMedAgent Frontier cost per query ($1.32) is ∼38× that of spatiAlytica Frontier ($0.035, Supplementary Table 7).
