## Supplementary Table 10 for "spatiAlytica: Viewer-Grounded Multimodal Agentic System for Interactive Spatial Omics Analysis"

**Supplementary Table 10  Ablation study: full-system baseline accuracy (%) and accuracy change (pp) when removing error correction (no-EC) or memory (no-mem).** Each cell shows mean ± s.d. across $N$=3 independent runs; ablation cells additionally report (Δ vs. full system). Negative Δ indicates the sub-system improves accuracy when present. Memory ablation in ST is not applicable (N/A).

| | | **spatiAlytica Agent Configuration** | | | |
|---|---|---|---|---|---|
| **Ablation** | **Mode** | **Open-source (Kimi-K2.5)** | **Compact (GPT-4.1-mini)** | **Standard (GPT-5-mini)** | **Frontier (GPT-5.2)** |
| Full system | ST | $54.7 \pm 1.4$ | $69.5 \pm 1.8$ | $73.1 \pm 4.5$ | $83.5 \pm 0.9$ |
| No error correction | ST | $53.6 \pm 2.1$ (−1.1) | $65.0 \pm 1.8$ (−4.5) | $65.8 \pm 2.4$ (−7.3) | $78.4 \pm 3.1$ (−5.1) |
| No memory | ST | N/A | | | |
| Full system | MT | $57.5 \pm 0.9$ | $54.4 \pm 2.1$ | $62.8 \pm 2.7$ | $67.4 \pm 1.3$ |
| No error correction | MT | $55.9 \pm 1.9$ (−1.6) | $54.9 \pm 2.2$ (+0.5) | $60.1 \pm 3.0$ (−2.7) | $68.6 \pm 1.8$ (+1.2) |
| No memory | MT | $38.1 \pm 2.3$ (−19.4) | $41.0 \pm 0.6$ (−13.4) | $43.4 \pm 1.2$ (−19.4) | $46.0 \pm 0.5$ (−21.4) |
| Full system | TF | $62.3 \pm 1.7$ | $61.9 \pm 2.0$ | $70.0 \pm 1.6$ | $77.5 \pm 2.8$ |
| No error correction | TF | $61.8 \pm 1.2$ (−0.5) | $59.2 \pm 1.9$ (−2.7) | $68.2 \pm 0.4$ (−1.8) | $77.8 \pm 2.1$ (+0.3) |
| No memory | TF | $50.2 \pm 0.7$ (−12.2) | $55.7 \pm 2.1$ (−6.2) | $59.1 \pm 1.7$ (−10.9) | $62.9 \pm 0.9$ (−14.6) |
