## Supplementary Table 11 for "spatiAlytica: Viewer-Grounded Multimodal Agentic System for Interactive Spatial Omics Analysis"

**Supplementary Table 11 Per-query compute and latency comparison for the Frontier (GPT-5.2) configuration across spatiAlytica, BioMANIA, and BioMedAgent on the spatiAlyticaBench-ST (222 questions).** Token counts and latencies are per-query averages; LLM calls count main-agent requests per question.

| Metric | spatiAlytica F3 | BioMANIA F3 | BioMedAgent F3 |
|---|---|---|---|
| *Token usage* | | | |
| Total tokens/query | 45,178 | 11,556 | 102,267 |
| Prompt tokens/query | 44,912 | 10,872 | 87,348 |
| Completion tokens/query | 266 | 683 | 14,920 |
| *Latency* | | | |
| Mean elapsed (s) | 89.6 | 180.3 | 313.7 |
| Median elapsed (s) | 51.5 | 51.3 | 260.8 |
| Min / Max (s) | 17 / 600 | 20 / 1,200 | 98 / 1,853 |
| *Latency by outcome* | | | |
| Mean elapsed, correct (s) | 85.4 | 72.8 | 291.3 |
| Mean elapsed, wrong (s) | 111.0 | 228.8 | 325.2 |
| *Compute per question* | | | |
| Main-agent LLM calls | 2.63 | 4.65 | 36.53 |

spatiAlytica achieves the highest ST accuracy (83.5%) with 2.6× fewer LLM calls per question than BioMANIA (4.65) and ∼14× fewer than BioMedAgent (36.53), and with mean latency comparable to BioMANIA and 3.5× faster than BioMedAgent. The BioMedAgent pipeline issues roughly 37 main-agent calls per question across six sub-agents (Linguist, Prompt Engineer, Tool Scorer, Workflow Designer, Programmer, Summary Analyst), explaining its high token usage and latency.
