## Supplementary Table 12 for "spatiAlytica: Viewer-Grounded Multimodal Agentic System for Interactive Spatial Omics Analysis"

**Supplementary Table 12 spatiAlyticaBench-MT performance (mean ± s.d., $N$=3 independent runs).** The benchmark comprises 178 questions organized as 11 multi-hop conversations (13–21 turns each). MT: cascading protocol; TF: teacher-forcing protocol. Best accuracy per mode is shown in **bold**.

| | **spatiAlytica Agent Configuration** | | | |
|---|---|---|---|---|
| **Metric** | **Open-source (Kimi-K2.5)** | **Compact (GPT-4.1-mini)** | **Standard (GPT-5-mini)** | **Frontier (GPT-5.2)** |
| MT accuracy (%) | $57.5 \pm 0.9$ | $54.4 \pm 2.1$ | $62.8 \pm 2.7$ | $\mathbf{67.4 \pm 1.3}$ |
| TF accuracy (%) | $62.3 \pm 1.7$ | $61.9 \pm 2.0$ | $70.0 \pm 1.6$ | $\mathbf{77.5 \pm 2.8}$ |
| MT→TF gain (pp) | +4.8 | +7.5 | +7.2 | +10.1 |
| Avg. tokens/query (MT) | 58,272 | 57,775 | 74,050 | 45,091 |
| Avg. cost/query (MT, $) | 0.009 | 0.013 | 0.014 | 0.039 |
