## Supplementary Table 13 for "spatiAlytica: Viewer-Grounded Multimodal Agentic System for Interactive Spatial Omics Analysis"

**Supplementary Table 13 BioMANIA performance on spatiAlyticaBench-MT split by evaluation library (mean ± s.d., $N$=3 independent runs).** The benchmark comprises 178 questions organized as 11 multi-hop conversations (13–21 turns each); each conversation is run twice (once with scanpy and once with squidpy). MT: cascading protocol; TF: teacher-forcing protocol. Best accuracy per mode × library is shown in **bold**.

| | **BioMANIA Agent Configuration** | | | |
|---|---|---|---|---|
| **Metric** | **Open-source (Kimi-K2.5)** | **Compact (GPT-4.1-mini)** | **Standard (GPT-5-mini)** | **Frontier (GPT-5.2)** |
| *Scanpy* | | | | |
| MT accuracy (%) | **47.1 ± 1.2** | 19.6 ± 2.7 | 40.3 ± 8.3 † | 27.4 ± 10.1 † |
| TF accuracy (%) | **56.6 ± 0.0** | 55.4 ± 0.3 | 55.6 ± 0.5 | 55.1 ± 1.8 |
| MT→TF gain (pp) | +9.5 | +35.8 | +15.3 | +27.7 |
| *Squidpy* | | | | |
| MT accuracy (%) | 52.0 ± 0.0 | 43.9 ± 8.1 † | **52.4 ± 0.2** | 37.8 ± 12.6 † |
| TF accuracy (%) | 56.6 ± 0.0 | **57.1 ± 0.0** | 56.6 ± 0.0 | 56.4 ± 0.3 |
| MT→TF gain (pp) | +4.6 | +13.2 | +4.2 | +18.6 |

†: cross-run s.d. > 3 pp. In TF mode, all backends and both libraries converge to 55.1–57.1% (spread ≤ 2.0 pp), indicating that when cascading error is eliminated BioMANIA's single-turn ceiling is the binding constraint regardless of backend or library. MT performance is sharply lower and shows backend-dependent inversion: Kimi-K2.5 leads in MT (47.1%–52.0%) while GPT-5.2 is the lowest (27.4%–37.8%), reversing the rank order observed in spatiAlytica. Large MT→TF gains for GPT backends (+13.2 to +35.8 pp) quantify cascading-error loss in BioMANIA's multi-turn architecture.
