## Supplementary Table 14 for "spatiAlytica: Viewer-Grounded Multimodal Agentic System for Interactive Spatial Omics Analysis"

**Supplementary Table 14 Code QA multi-turn robustness checks.** Conversation-level 95% bootstrap CIs (2,000 resamples) for MT and TF pass rates; rankings are unchanged relative to the pair-level estimate. ARI-threshold sensitivity was also verified on single-turn ($t \in \{0.80, 0.85, 0.90, 0.95\}$): all 23 ARI-judged verdicts across backends pass at every threshold, so the 0.9 cutoff does not affect the reported accuracies. Plot-evaluation weight sensitivity was verified by recomputing combined scores at alternative code/plot weightings; backend rank orders on MT are preserved at $(0.1, 0.9)$, $(0.3, 0.7)$, $(0.5, 0.5)$, and $(0.7, 0.3)$.

| | MT | | | TF | | |
|---|---|---|---|---|---|---|
| **Configuration** | **Pair (%)** | **95% CI** | $n_{\text{conv}}$ | **Pair (%)** | **95% CI** | $n_{\text{conv}}$ |
| Compact (GPT-4.1-mini) | 55.4 | [49.6, 61.0] | 9 | 57.0 | [49.3, 65.1] | 11 |
| Standard (GPT-5-mini) | 63.5 | [61.8, 65.5] | 9 | 63.1 | [55.0, 70.9] | 11 |
| Frontier (GPT-5.2) | **70.3** | [65.8, 75.3] | 9 | **71.0** | [64.7, 77.3] | 11 |
| Open-source (Kimi-K2.5) | 57.7 | [54.1, 61.8] | 9 | 56.6 | [50.4, 62.8] | 11 |
