## Supplementary Table 15 for "spatiAlytica: Viewer-Grounded Multimodal Agentic System for Interactive Spatial Omics Analysis"

**Supplementary Table 15 spatiAlyticaBench-ImageQA judge-pass rates (mean ± s.d., $N$=2 runs; 7,350 QA pairs from 1,295 spatial transcriptomics subfigures).** GPT-5.2 serves as the fixed judge. A score $\geq 3$ (1–5 scale) is a pass. Low pass rates for Standard and Open-source are driven by high empty-response rates. The metric measures LLM-judge agreement with LLM-generated reference answers, not ground-truth accuracy.

| | **spatiAlytica Agent Configuration** | | | |
|---|---|---|---|---|
| **Metric** | **Open-source (Kimi-K2.5)** | **Compact (GPT-4.1-mini)** | **Standard (GPT-5-mini)** | **Frontier (GPT-5.2)** |
| Accuracy (%) | $26.8 \pm 2.3$ | $81.1 \pm 0.4$ | $25.7 \pm 0.1$ | $\mathbf{86.0 \pm 0.7}$ |
| Avg. judge score (1–5) | 1.87 | 3.70 | 1.94 | **3.99** |
