## Supplementary Table 16 for "spatiAlytica: Viewer-Grounded Multimodal Agentic System for Interactive Spatial Omics Analysis"

**Supplementary Table 16 spatiAlyticaBench-ImageQA outcome breakdown (mean % of pairs ± s.d. across full runs; 7,350 pairs per run, $N$=2 runs except Open-source where one partial rerun is excluded).** Image-level cluster-aware 95% CI on the pass rate (Correct column) is reported alongside the pair-level estimate using a 2,000-sample bootstrap over 1,295 subfigures; rankings are unchanged.

| **Configuration** | **Empty (%)** | **Incorrect (%)** | **Correct (%)** | **Pair-level** | **Image-bootstrap 95% CI** |
|---|---|---|---|---|---|
| Compact (GPT-4.1-mini) | $0.0 \pm 0.00$ | $18.9 \pm 0.45$ | $81.1 \pm 0.45$ | 81.08 | [79.83, 82.31] |
| Standard (GPT-5-mini) | $70.1 \pm 0.50$ | $4.3 \pm 0.48$ | $25.6 \pm 0.02$ | 25.59 | [24.45, 26.80] |
| Frontier (GPT-5.2) | $0.1 \pm 0.03$ | $14.0 \pm 0.62$ | $\mathbf{86.0 \pm 0.64}$ | 86.0 | [84.94, 86.96] |
| Open-source (Kimi-K2.5) | $68.2 \pm 2.88$ | $5.1 \pm 0.45$ | $26.6 \pm 2.42$ | 26.64 | [25.57, 27.78] |
