## Supplementary Table 17 for "spatiAlytica: Viewer-Grounded Multimodal Agentic System for Interactive Spatial Omics Analysis"

**Supplementary Table 17 spatiAlyticaBench-ImageQA judge-pass rates at alternative score cutoffs and conditional on non-empty response emission (mean ± s.d. across $N$=2 independent runs; 7,350 pairs per run).** The Non-empty $\geq 3$ column restricts the denominator to responses where a non-empty answer was emitted. Tier rank order (Frontier/Compact > Standard/Open-source) is preserved at all three judge cutoffs.

| **Configuration** | **All $\geq 2$ (%)** | **All $\geq 3$ (%)** | **All $\geq 4$ (%)** | **Non-empty $\geq 3$ (%)** |
|---|---|---|---|---|
| Open-source (Kimi-K2.5) | $30.8 \pm 2.7$ | $26.6 \pm 2.4$ | $21.0 \pm 2.3$ | $83.9 \pm 0.0$ |
| Compact (GPT-4.1-mini) | $95.0 \pm 0.3$ | $81.1 \pm 0.5$ | $64.7 \pm 1.2$ | $81.1 \pm 0.5$ |
| Standard (GPT-5-mini) | $29.0 \pm 0.3$ | $25.7 \pm 0.0$ | $23.2 \pm 0.0$ | $85.6 \pm 1.4$ |
| Frontier (GPT-5.2) | $\mathbf{97.4 \pm 0.3}$ | $\mathbf{86.0 \pm 0.6}$ | $\mathbf{73.8 \pm 1.3}$ | $\mathbf{86.0 \pm 0.6}$ |
