## Supplementary Table 18 for "spatiAlytica: Viewer-Grounded Multimodal Agentic System for Interactive Spatial Omics Analysis"

**Supplementary Table 18  Reproducibility details and pinned software dependencies for all benchmark evaluations.** Agent model assignments follow Table 6. All evaluations used the same judge model, decoding settings, and codebase version. Package versions are those installed in the `spatialytica-env` conda environment used to run the benchmarks; floor constraints in `requirements.txt` permit newer compatible releases. A complete dependency manifest is provided in the repository.

| Package / Parameter | Version / Value |
|---|---|
| *Spatial omics* | |
| AnnData [28] | 0.12.3 |
| Scanpy [49] | 1.11.5 |
| Squidpy [38] | 1.8.1 |
| libpysal | 4.13.0 |
| esda | 2.8.0 |
| GeoPandas | 1.1.1 |
| python-igraph | 1.0.0 |
| louvain | 0.8.2 |
| *Agent and LLM layer* | |
| LangChain [50] | 0.3.27 |
| LangGraph | 1.0.1 |
| LiteLLM | 1.78.7 |
| OpenAI SDK | 1.109.1 |
| instructor | 1.11.3 |
| tiktoken | 0.12.0 |
| Chroma | 1.2.1 |
| *Numerical core and runtime* | |
| Python | 3.11 |
| NumPy | 2.3.4 |
| pandas | 2.3.3 |
| pydantic | 2.12.3 |
| napari | 0.6.6 |
| *Reproducibility parameters* | |
| Judge model (all evaluations) | GPT-5.2 (gpt-5.2-2025-12-11) |
| Temperature (all agent calls) | 0.0 |
| Seed | 42 |
| Max retries per LLM call | 3 |
| Max error-correction iterations | 3 |
| Routing / fallback | LiteLLM; no fallback between providers |
| Codebase version (evaluated) | 0.3.5 |

Async streaming calls use the provider default temperature. The verbatim system prompts and role-specific instructions for all six core sub-agents (orchestrator, code writer, code debugger, spatial VQA, user response, and data resolver) plus the niche-naming sub-routine are version-pinned in the repository at `src/napari_spatial_gpt/prompts/` and may be inspected directly.
