## Supplementary Note 1 for "spatiAlytica: Viewer-Grounded Multimodal Agentic System for Interactive Spatial Omics Analysis"

### Supplementary Note 1 Detailed Methods

This supplementary note provides implementation details for the methods described in the Online Methods section.

#### Supplementary Note 1.1 System architecture details

***Widget layer (Napari GUI).***

The Napari dock widget is the primary user entry point for interactive analysis. It supports loading spatial datasets (`AnnData`), configuring key columns (e.g., spatial coordinates, cell-type labels, sample identifiers), selecting color mappings, and displaying outputs (plots, tables, and narratives). The widget also maintains the UI state representation describing the current viewer configuration, including active spatial layers, color legends, user annotations, selected layer, and the current viewport (field of view). This state is treated as first-class context for downstream agent reasoning.

***Closed-loop coupling between sub-agents, UI state, and user interpretation.***

spatiAlytica implements closed-loop coupling by synchronizing (i) tool-based computation, (ii) the UI state representation, and (iii) agent reasoning. After each analytical or plotting action, the system updates both the visualization (Napari layers and legends) and the corresponding UI state in memory. This enables iterative workflows in which users zoom into regions, toggle layers, add annotations, and ask region-specific questions while the system maintains a consistent and auditable analysis trajectory. The spatial VQA sub-agent operates directly on the current UI state (and, when required, screenshots of the active canvas), ensuring that image-centered questions remain aligned to what is visible in the viewer.

#### Supplementary Note 1.2 Core sub-agent specifications

***Orchestrator Sub-Agent.***

The orchestrator sub-agent is the central coordinator responsible for (i) parsing the user's intent and classifying it as exploratory analysis (EA) or interpretation (IR), (ii) retrieving relevant context from the agentic memory sub-system (including prior tool results, the current UI state representation, and persisted result registries), (iii) selecting and sequencing tools from the hybrid tool repository, and (iv) delegating subtasks to specialized agents when required. The orchestrator constructs explicit action plans (rather than free-form responses) so that results are grounded in executed code, validated tool outputs, and tracked visualization state. Tool execution proceeds in an iterative loop: the LLM selects one or more tools, their results are appended to the conversation, and the LLM is re-queried until it produces a final textual response or a maximum of 10 iterations is reached.

***Data resolver sub-agent.***

The data resolver sub-agent maps high-level biological terms (e.g., "cell type", "niche", "disease group") and feature identifiers (e.g., gene symbols) onto concrete dataset keys in `AnnData.obs`, `AnnData.var_names`, and `AnnData.uns`. Resolution proceeds in two phases: first, a namespace relevance classifier determines which data compartments (metadata, variable names, or stored results) are pertinent to the query; second, within each relevant namespace, the agent performs exact and fuzzy matching with domain-specific heuristics (e.g., preferring text-label cell-type columns over numeric IDs, distinguishing existing niche assignments from the original cell-type column for new niche analyses). Results are cached with an LRU cache (50 entries) to avoid redundant LLM calls during multi-step workflows.

***Code writer sub-agent.***

The code writer sub-agent generates executable Python code from natural-language task descriptions provided by the orchestrator. Before writing code, the agent may invoke tools (e.g., `ResolveColumnNamesTool`, `GetColumnValuesTool`) in an internal planning loop of up to 10 rounds to inspect dataset structure and validate column names. Generated code executes in a controlled environment with access to standard spatial omics libraries (Scanpy, Squidpy, Matplotlib). If execution fails, control passes to the code debugger sub-agent for error correction.

***Code debugger sub-agent.***

The code debugger sub-agent corrects failed Python code produced by the code writer sub-agent. When code execution raises an error, the code debugger sub-agent receives the failed code, exception trace, and execution context (including the current `AnnData` schema and whether the interactive QA mode is turned on) and proposes minimal corrective edits. The corrected code is then re-executed under the same controlled environment. This execute-correct-retry cycle repeats for up to three attempts (configurable via `max_tries`), after which the system reports the failure. The agent handles common failure patterns in spatial omics code, including categorical data type mismatches, sparse matrix extraction, incorrect Scanpy and Squidpy API usage, and column-name resolution errors.

###### *Spatial VQA sub-agent.*

The spatial VQA sub-agent enables multimodal, image-centered interrogation of spatial visualizations. It accesses the UI state representation (active layers, legends, viewport, and annotations) and can analyze screenshots of the current Napari canvas to answer questions about spatial organization, co-localization, boundaries, and regional enrichment. The interpretation is conditioned on the active visualization state and whether the interactive QA mode is turned on. Hence, the responses remain synchronized with the user's current view (e.g., a zoomed-in tumor–stroma interface), number of cells in the view, and the exact color mappings and overlays currently displayed.

###### *User response sub-agent.*

The user response sub-agent synthesizes tool outputs (tables, statistics, plots, logs) and sub-agent findings (e.g., data resolver mappings, VQA interpretations) into clear, structured narrative responses. It formats results for interactive consumption, highlights salient effect sizes and spatial patterns, and proposes sensible next analytical steps while maintaining strict grounding in recorded outputs.

#### Supplementary Note 1.3 Hybrid tool repository details

The hybrid tool repository exposes 13 type-validated tools to the orchestrator via OpenAI-compatible function-calling schemas. These span four categories: (i) *data inspection*: `ResolveColumnNamesTool` (fuzzy-matches biological terms to `AnnData` keys) and `GetColumnValuesTool` (enumerates unique categorical levels); (ii) *analytical workflows*: `DifferentialExpressionTool` (flexible pairwise differential expression (DE) analysis with Wilcoxon, $t$-test, or logistic regression), `NicheAnalysisTool` (KNN neighborhood composition clustering with automatic biological naming), `MoransIAnalysisTool` (univariate and bivariate spatial autocorrelation via LISA), and `CellDensityTool` (local cell-type density estimation); (iii) *visualization*: `SpatialPlotTool` (interactive Napari layers with categorical or continuous coloring), `UMAPPlotTool` (embedding visualizations), `PieChartTool` (categorical distributions), and `WriteCodeTool` (general-purpose code generation for custom plots and computations); and (iv) *image analysis and response*: `AnalyzeImageTool` (multimodal VQA on current viewer state), `RespondWithTextTool` (direct narrative responses), and `TerminateTool` (session control). Structured tools (`DifferentialExpressionTool`, `NicheAnalysisTool`, `MoransIAnalysisTool`, `CellDensityTool`, and `SpatialPlotTool`) use Pydantic request schemas for parameter validation.

###### *Code execution system.*

When a request requires bespoke computation or custom plotting, the `WriteCodeTool` delegates to the code writer sub-agent, which generates executable Python code in a controlled environment with comprehensive output capture (stdout/stderr) and artifact retention (figures, tables, intermediate variables). The dataset is exposed as `adata` and `data` for compatibility with common conventions, and the environment provides pre-imported scientific Python libraries (`pandas`, `numpy`, Scanpy, Squidpy, `matplotlib`, and `seaborn`). Non-spatial plots are rendered through a resizable Matplotlib widget (`plot_in_matplotlib_widget`), while spatial plots are routed exclusively through `SpatialPlotTool` to maintain Napari layer interactivity. All stochastic operations are seeded with `random_state=42` for reproducibility. Each execution is logged to enable end-to-end provenance.

#### Supplementary Note 1.4 Agentic memory sub-system details

###### *Memory implementation.*

The agentic memory sub-system is implemented using LangChain [50]. Short-term memory stores the current conversation history as a sequence of human and assistant messages using `InMemoryChatMessageHistory`. Long-term memory uses an `InMemoryStore` backed by JSON persistence to retain validated column and gene mappings, analysis state (dataset identifiers, active filters, key findings), and a registry of persisted results that tracks which columns, embedding keys, and stored objects were added to the `AnnData` object during which queries. Intermediate result memory caches recent tool outputs (e.g., differential expression tables, niche composition vectors, code execution results) and recent tool call specifications for follow-up reuse. When an LLM is available for memory operations, the system additionally generates 2–3 sentence conversation summaries to provide condensed context for downstream agents.

All memory is persisted to disk as JSON files under the user's `SPATIALYTICA_HOME/memory/` directory, enabling session continuity.

***Interaction logging and provenance.***

spatiAlytica maintains comprehensive session-based interaction logs that capture all user interactions, agent responses, and generated artifacts for full analytical provenance. Each analysis session is assigned a unique session identifier and stored in a dedicated session folder containing: (i) query-focused markdown logs with user questions and agent responses, (ii) agent decision and tool call logs, (iii) generated code logs, (iv) structured telemetry events in newline-delimited JSON (NDJSON) format capturing per-query timing breakdowns (memory context retrieval, agent calls, tool execution, code execution, UI updates), API call metadata (model, request/response sizes, durations), and asset paths, (v) timing summary spreadsheets with per-query and aggregate statistics, and (vi) an assets directory with saved visualizations (PNG) and intermediate analysis results (CSV). This multi-format logging enables full replay of analytical workflows, quantitative performance analysis, and transparent documentation for manuscript preparation and peer review. In the GUI, feedback can be captured as structured signals associated with prompts, tool selections, and output usefulness. In the CLI, all actions and outputs are logged, enabling high-throughput execution and full replay of analyses for validation, auditing, and manuscript preparation.

#### Supplementary Note 1.5 Human validation of the spatiAlyticaBench-ImageQA judge

Because both the QA pairs and the judge scores in the spatiAlyticaBench-ImageQA are LLM-generated, we conducted a blinded human evaluation to calibrate the GPT-5.2 judge against expert human judgment and to assess whether the model answers are biologically correct independently of the LLM-generated reference answers.

***Sampling.***

From the 7,350 QA pairs evaluated under the Frontier configuration (GPT-5.2), we drew a stratified subsample of 100 pairs balanced across three question types (factual, spatial, interpretive) and five judge-score bins (1–5), yielding 6–7 pairs per stratum (3 types × 5 bins = 15 strata). Score-bin stratification ensures adequate calibration coverage at the pass/fail decision boundary (judge score $\geq 3$), which is the threshold that determines the headline judge-pass rate. Within each stratum, pairs were sampled uniformly at random with a preference for unique source images to reduce image-level dependence. Nine distractor pairs were generated by replacing the model answer with an answer from an unrelated image–question pair drawn from a different question type, yielding 109 total pairs per annotator. Distractor pairs serve as attention checks: an alert annotator should reject clearly mismatched answers.

***Annotators and blinding.***

Four senior bioinformaticians (two with PhDs) served as independent raters. All had deep technical background and hands-on experience with spatial transcriptomics platforms. Annotators were recruited by open request on the basis of availability, without selection for prior experience with spatiAlytica or for agreement with any prior scoring. All raters were blinded to model identity, GPT-5.2 judge score, distractor status, and pair provenance; they saw only the spatial image, the question text, and the model answer. Pair order was independently randomized per annotator (seeded by annotator ID for reproducibility) so that no two annotators evaluated pairs in the same sequence. The design is fully crossed: all four raters scored the same 100 real pairs plus their 9 randomly assigned distractors.

***Annotation task.***

Each annotator independently scored every pair on two axes:

1. A 5-point rubric: 5 = completely correct and complete; 4 = correct with minor omissions; 3 = partially correct, acceptable; 2 = partially wrong, unacceptable; 1 = incorrect or hallucinated.
2. A binary accept/reject decision: "Is the answer biologically correct and would you accept it?" (Yes/No).

An optional free-text notes field allowed annotators to flag specific hallucinations or concerns. Critically, the binary question asks about *biological correctness*, not about agreement with the LLM-generated reference answer; this decouples the human evaluation from the GPT-4o question-generation pipeline.

***Adjudication rule.***

With four raters and a fully crossed design, human ground truth for each pair is the majority of four binary decisions with a $\geq 2/4$ acceptance threshold, and the median of four 5-point scores for the rubric. Ties at 2–2 on the binary axis are resolved in favor of acceptance (reflecting the dominant marginal of the dataset, on which the human accept rate is $\geq 97\%$). As a sensitivity check, we also recomputed all downstream statistics under a strict $\geq 3/4$ rule and a unanimity rule (all four agree); the reported pass rates, AC1 values, and calibration-adjusted headline shift by $\leq 2$ pp across adjudication rules.

***Statistical analyses.***

Inter-annotator agreement over all four human raters was quantified using Krippendorff's $\alpha$ on binary acceptance (nominal metric) and on the 5-point rubric (interval metric), reported overall and stratified by question type. Pairwise agreement was additionally reported via Cohen's $\kappa$ for every rater pair. Judge calibration was assessed by treating the majority-adjudicated human labels as ground truth and computing a $2 \times 2$ confusion matrix of the judge's binary pass/fail decision (threshold $\geq 3$) against the human binary accept/reject, with conditional acceptance probabilities $P(H|\text{judge pass})$ and $P(H|\text{judge fail})$ reported with Wilson 95% CIs. A calibration-adjusted spatiAlyticaBench-ImageQA accuracy is computed as $\hat{p}_{\text{adj}} = \pi \cdot P(H|\text{pass}) + (1-\pi) \cdot P(H|\text{fail})$, where $\pi = 0.859$ is the judge pass rate on the full 7,350-pair benchmark under the Frontier configuration; a 95% confidence interval is derived from a 10,000-sample bootstrap resampling pairs with replacement within each judge-pass stratum. This calibration-adjusted value is reported as a sensitivity check under the reference-anchored protocol and is not the headline metric; the headline is the direct human majority pass rate on the 100-pair subsample, and the robustness of this pass rate to judge model family is established via the cross-family LLM-judge analysis below. Leave-one-rater-out sensitivity analysis re-computed all primary statistics on each $(N-1)$-rater subset to verify that no single rater drives any reported quantity. A per-bin calibration curve (human accept fraction against judge score bin 1–5), a per-question-type human acceptance breakdown with Wilson CIs, and a per-rater diagnostic of rubric distribution, accept rate, and winsorized time-on-pair are reported in the Supplementary Figures. F1 and AUROC statistics are reported for completeness but de-emphasized because the low-rejection base rate on the validation subset makes these metrics unstable. Attention-check distractors were analyzed separately: the distractor rejection rate across all raters was 100%.

***Cross-family LLM-judge analysis.***

Because both the benchmark questions and the reference answers were generated by GPT-4o and the original judge is itself an LLM (GPT-5.2), we conducted an independent cross-family LLM-judge analysis on the same 100-pair subsample to test whether the benchmark signal is robust to the substitution of judge model family and to the removal of the GPT-4o reference answer from the evaluation loop. Two LLM judges from two corporate entities were evaluated: GPT-5.2 (OpenAI) and Claude Opus 4.6 high (Anthropic). Every judge received exactly the same rubric, presented as a verbatim text block with explicit instructions to reject questions containing false premises and to decline to answer questions whose required content is not present in the image. Critically, no judge received the GPT-4o-generated reference answer: each judge saw only the image, the question, and the assistant's answer, and produced a 1–5 rubric score with a one-sentence rationale. This reference-free protocol mirrors the human-annotator protocol exactly and excludes the GPT-4o reference from the judge loop. Pairwise agreement between the two LLM judges and between each judge and the human majority vote was quantified using raw percent agreement, Cohen's $\kappa$, and Gwet's AC1 (all on the binary pass axis with threshold $\geq 3$). Gwet's AC1 is reported as the primary agreement coefficient because Cohen's $\kappa$ is known to be deflated when marginal probabilities are highly skewed (the first-kind paradox), which is the case here due to the $\geq 97\%$ human accept rate. Judge-family-specific pass rates with 95% Wilson CIs, within-judge across-protocols agreement (GPT-5.2 with-reference vs without-reference on the same 100 items), and per-question-category pass rates (over seven categories: legend reading, descriptive, pattern reading, quantitative rank-order, adversarial false-premise, adversarial distractor, and multi-step reasoning) are reported in the accompanying analysis artifacts. The cross-family analysis is fully reproducible: the universal judge prompt, per-judge runner scripts, joined analysis script, and all output CSVs are provided with the manuscript code and can be re-run end-to-end in under one hour on a single workstation.

#### Supplementary Note 1.6 Automated evaluation pipeline details

***Hierarchical comparison strategy.***

The judge applies comparisons in order of decreasing determinism. First, when the GT stores results in a known `AnnData` location (e.g., `adata.uns['rank_genes_groups']`), the judge extracts that specific key from both the GT and agent `AnnData` objects and compares them programmatically. Second, the pipeline captures snapshots of the `AnnData` object before and after code execution in both subprocesses; new or changed entries in `.obs`, `.uns`, `.obsm`, and `.obsp` are compared via a structured diff. Third, explicit return values from GT and agent code are compared using a recursive programmatic comparator (described below). Fourth, when no structured objects are available (e.g., when both outputs consist solely of printed text), an LLM judge [32] serves as a fallback. Finally, a both-empty guard prevents inflated accuracy: when both GT and agent produce empty or null standard output on a data question with no structured comparison available, the verdict defaults to "not equivalent" rather than granting a false positive from in-place operations that silently modify state without producing output.

***Programmatic object comparison.***

The core comparison function dispatches on output type to apply appropriate equivalence criteria. NumPy arrays are compared using `np.allclose` with relative tolerance $10^{-5}$ and absolute tolerance $10^{-8}$, treating NaN values as equal. Pandas DataFrames are first compared via `pd.testing.assert_frame_equal` with column-reordering tolerance and relative tolerance $10^{-5}$; when this strict check fails, a tiered fallback requires both a maximum absolute difference no greater than $2 \times 10^{-3}$ and either a mean absolute difference below $5 \times 10^{-4}$ or a mean relative difference below 5%, preventing false matches on small-magnitude values such as $p$-values. Pandas Series are compared element-wise with dtype-aware tolerance; Boolean Series use direct element-wise comparison rather than aggregate counts. Sparse matrices are converted to compressed sparse row format and compared on their constituent `.data`, `.indices`, and `.indptr` arrays, with a dense-comparison fallback when sparsity structures differ. Scalar numeric values use `np.isclose` with relative tolerance $10^{-5}$; strings are compared by exact equality. Dictionaries are compared recursively, requiring GT keys to be a subset of agent keys (the agent may compute additional fields). When keys differ only in formatting (whitespace, underscores, hyphens, or case), a normalized-key fallback resolves cosmetic mismatches. Lists and tuples undergo positional comparison first; upon failure, a multiset comparison (preserving duplicate counts) checks for order-independent equivalence, which is relevant for inherently unordered results such as cell-type inventories.

***Handling stochastic algorithms.***

Several bioinformatics operations produce outputs that are correct yet non-identical across runs due to algorithmic stochasticity. For clustering results from algorithms such as Leiden or Louvain, exact label matching would yield false negatives because cluster label assignment is arbitrary. We therefore assess partition similarity using the Adjusted Rand Index (ARI). When the comparison target is a single Series of cluster labels, we require ARI $\geq 0.9$ to indicate equivalence. When the target is a DataFrame containing one or more clustering columns (e.g., `leiden`, `louvain`, or per-resolution columns), each column is checked independently and we require ARI $\geq 0.8$ on every column, reflecting the looser threshold appropriate when several stochastic clusterings are reported in a single object. For embeddings produced by UMAP or t-SNE, where coordinate-level equality is not meaningful (the embedding is invariant only up to rotation, reflection, and translation), we instead compute the absolute Pearson correlation between corresponding columns of the GT and agent embeddings, with high correlation on every column treated as equivalence. For principal component analysis, eigenvectors are sign-ambiguous; the judge detects and corrects column-wise sign flips prior to numerical comparison. For `rank_genes_groups` DataFrames, the comparator first checks that the four core columns (`names`, `scores`, `pvals`, `pvals_adj`) are present on both sides and then compares the gene-name lists per group, tolerating the small number of tie-breaking differences that arise near significance boundaries. For numeric comparisons, scalars and Series that differ by exactly a factor of 100 are treated as equivalent under a proportion-versus-percentage rescaling rule: a value of 0.5 and a value of 50.0 refer to the same underlying quantity, reported as a fraction in one case and as a percentage in the other. Metadata keys carrying no semantic content (e.g., `random_state`, `seed`, `params`) are stripped from both GT and agent dictionaries before comparison.

***Key-name-agnostic recovery and structural alignment.***

For questions that instruct the agent to store results without specifying the exact key name (e.g., "save that summary table"), the judge permits key-name differences in `AnnData` storage. When diff comparison fails due to key mismatch, the pipeline extracts payloads from both GT and agent diffs and compares values within the same container type (`obs`-to-`obs`, `uns`-to-`uns`). This recovery is gated: questions that explicitly name the storage key require exact key matching. Additionally, when type or structural mismatches render programmatic comparison inconclusive, an alignment step uses a lightweight LLM (GPT-4o-mini) to generate a Python expression transforming the agent output to match the GT structure (e.g., extracting the `.X` attribute from an `AnnData` object or converting a dictionary to a Series). The transformed result must then pass the same strict programmatic comparison. Post-alignment validation rejects expressions that collapse multi-element data to scalars or alter data dimensionality by more than tenfold.

***Plot evaluation.***

For visualization tasks, the pipeline uses a weighted combination of code similarity (weight 0.1) and visual similarity (weight 0.9), both assessed by GPT-5.2 on a 1–5 scale comparing the agent's output against the GT. The heavy visual weighting reflects the fact that multiple plotting interfaces (Scanpy high-level functions, Matplotlib, Squidpy) can produce equivalent visualizations through substantially different code. The combined score $0.1 \cdot \text{code} + 0.9 \cdot \text{plot}$ counts as a pass at $\geq 3.0$, with confidence labelled `high` at $\geq 4.0$, `medium` at $\geq 2.5$, and `low` otherwise. Each judge call is retried up to three times on transient API errors, with exponential backoff. The visual-similarity call submits the rendered PDF directly to the vision model. If the model rejects the native PDF payload, the judge converts the PDF to a 150-dpi PNG (using `pdftoppm`) and retries with the converted image. When the GT plot is unavailable due to an upstream infrastructure failure, the code-similarity score alone is used so the agent is not penalized for an issue outside its control.

***LLM judge.***

When programmatic comparison is inconclusive, an LLM judge (GPT-5.2) evaluates semantic equivalence of text outputs. The judge call is dispatched directly against the OpenAI API regardless of which API base the agent under evaluation uses. An Open-source agent backed by Together AI is therefore never authenticated against OpenAI on the agent path, and an OpenAI-backed agent never has its credentials forwarded to a third-party endpoint for the judge call. Each judge call passes `seed`=42 and constrains the response to JSON with the keys `equivalent` (boolean), `reasoning` (string), and `confidence` (`high`, `medium`, or `low`). The judge prompt enforces a conservative decision rubric. Outputs that both normalize to empty, `None`, or `null` payloads are equivalent at high confidence and short-circuited without an API call. Whitespace, code-fence wrappers, and incidental logging or warning lines (e.g., `UserWarning`, `Code executed successfully`) are normalized away before comparison. Substantive payload mismatches (one side carries tables, arrays, or dictionary content the other lacks) are not equivalent. Ranked gene lists are equivalent at $\geq 80\%$ overlap with medium confidence; this is the LLM-judge counterpart of the programmatic 80% rule, applied when the lists are presented as printed text rather than as structured DataFrames. Numeric outputs that differ by exactly a factor of 100 are equivalent under the proportion-versus-percentage rescaling rule. Uncertain cases default to `equivalent=false` with low confidence. The judge model is distinct from the agent model under evaluation, eliminating same-model evaluation bias.

***Error classification.***

Failed questions are categorized into five classes: execution errors (subdivided into syntax/runtime, timeout, and import/dependency failures), incorrect results (wrong values, wrong structure, or partial matches), infrastructure failures (API or file-system errors), logic errors (incorrect analytical approach or incomplete implementation), and unclassified errors. This taxonomy enables targeted diagnosis of agent failure modes.

***Reproducibility.***

All evaluation subprocesses enforce fixed random seeds. The seed value is 42 across Python's `random` module, NumPy, Scanpy, and Squidpy, and the same value is exported into `PYTHONHASHSEED` so that hash-randomized operations (e.g., set ordering) also become reproducible. The seed is propagated into Scanpy and Squidpy operations through runtime monkey-patching of the relevant function defaults so that any `random_state` argument left unset by the agent or GT code receives the canonical value rather than the framework default. The same seed is passed into every OpenAI judge call via the `seed`=42 request parameter, so that the JSON-format judge response is deterministic to the extent that the API surface supports it. GT outputs are precomputed and cached using a SHA-1 key derived from the code content, data file path, file size, and file modification timestamp, ensuring deterministic GT results across evaluation runs and enabling efficient re-evaluation when only agent configurations change.

#### Supplementary Note 1.7 Selection of BioMANIA as the primary baseline

BioMANIA was selected as the primary head-to-head baseline because it is the most directly comparable agentic biology system: it targets the same analytical libraries (Scanpy, Squidpy), accepts conversational natural-language input, and ships a complete end-to-end pipeline (user query → API disambiguation → code generation → execution with retries) that can be evaluated against the same ground-truth tasks we use for spatiAlytica.

We additionally benchmarked BioMedAgent [23] on the same 222 single-turn questions by substituting its multi-agent pipeline for spatiAlytica's agent call while reusing the identical ground-truth code, comparison pipeline, and LiteLLM dispatch layer. Single-turn BioMedAgent results across all four model configurations are reported in Supplementary Table 9 and compared on a per-query compute/latency basis in Supplementary Table 11. The Kimi-K2.5 backend could not be meaningfully evaluated within this framework because the Together AI endpoint does not reliably emit BioMedAgent's required XML response tags (`<STAGE>`, `<CODE>`), triggering BioMedAgent's response validator to abort the pipeline before code execution; this is a protocol-level incompatibility rather than a reasoning-capability measurement. BioMedAgent was not evaluated under the multi-turn or teacher-forcing protocols.

A head-to-head accuracy comparison against SpatialAgent [22] was not feasible within our evaluation framework. The public release (https://github.com/Genentech/SpatialAgent, accessed April 6, 2026) provides a component-level evaluation of its LLMToolSelector across eight hand-crafted bioinformatics tasks (Panel Design, Cell Type Annotation, Trajectory Analysis, Spatial Domain Detection, Cell-Cell Communication, Multimodal Integration, Spatial Niche Analysis, Literature Research), which measures tool-selection precision/recall/F1 over a catalog of 72 tools rather than end-to-end answer correctness. At the time of access, the release did not include the ground-truth code, scored datasets, or answer-grading infrastructure required to drive our shared comparison pipeline, and the full-system results in the associated preprint were not accompanied by a public reproducibility package. We therefore include SpatialAgent in the qualitative feature comparison (Supplementary Table 1) but not in the head-to-head accuracy benchmarks.

#### Supplementary Note 1.8 BioMANIA evaluation harness details

The evaluation harness substitutes BioMANIA's pipeline for spatiAlytica's agent call while reusing the identical ground-truth execution, structured comparison, and LLM-judge infrastructure. Specifically: (1) BioMANIA received the identical 222 single-turn questions with the same ground-truth code, preprocessed `AnnData` files, and hierarchical comparison pipeline. (2) BioMANIA's internal LLM calls were routed through the same LiteLLM dispatch layer to ensure identical model versions, API endpoints, and temperature settings ($T$=0). (3) BioMANIA operates through a multi-step conversational protocol that prompts the user for API disambiguation, parameter selection, and execution confirmation. The two-tier controller consists of a deterministic rule layer that auto-confirms execution prompts and selects the first candidate for disambiguation, and a lightweight LLM controller for parameter-filling states that receives only the original user question and the current BioMANIA prompt, with an explicit instruction to use no hidden benchmark answers or ground-truth code. Controller decisions are logged separately. This automated interaction represents an upper bound on BioMANIA's interactive performance, as real users may make suboptimal selections. (4) After BioMANIA's pipeline completes, the harness extracts all executed code fragments, concatenates them into a single script, and re-executes this code in an isolated subprocess with the same `AnnData`-diff instrumentation and structured output capture used for spatiAlytica, ensuring directly comparable result objects for the shared judgment pipeline. (5) BioMANIA's native execution retry mechanism was left active during evaluation, ensuring the comparison reflects BioMANIA at its full operational capability. BioMANIA requires the user to select a specific library backend (Scanpy or Squidpy) before each session; we evaluated each backend separately and report both results.
