## Supplementary Note 2 for "spatiAlytica: Viewer-Grounded Multimodal Agentic System for Interactive Spatial Omics Analysis"

### Supplementary Note 2 Study context template

Each spatiAlytica session begins with a structured study context that grounds the multimodal interactive agentic system in the biological and technical details of the dataset (see Methods, Section 3). The study context is provided as a free-text prompt at session initialization and is stored in the agentic memory subsystem, where it persists throughout the analysis and is accessible to all agents. Below we provide the general template followed by filled-in examples for the CODEX colorectal cancer and Kaposi's sarcoma case studies used in this work.

```
Template Example

We are studying [DISEASE/TISSUE] samples using [TECHNOLOGY].

Dataset structure:
 • Rename [original_column] to cell_types
 • Rename [original_column] to samples
 • Rename [original_column] to condition

Experimental groups:
 • Group A: [description]
 • Group B: [description]
 • Order of progression: [A to B to C]

Cell type notes:
 • [Cell type X] in this dataset represents [specific definition]
 • [Special state] is identified by [marker/gene criteria]

Visualization:
 • Use consistent colors for cell types and features across all plots
 • Sort results by [preferred ordering]

Goals:
1. [Primary objective]
2. [Secondary objective]
3. [Exploratory questions]
```

#### Filled-in Examples

```
A) CODEX CRC (CLR vs DII)

We are working with CODEX datasets of advanced-stage colorectal cancer (CRC).

Dataset structure:
 • Rename cluster label column to cell_types
 • Rename file name column to samples
 • Rename disease group column to disease_groups
 • Map disease_groups values: 1 to CLR, 2 to DII

Visualization:
 • Use a unique, consistent color per cell type across all plots

Goals:
 • Focus on spatial interactions within the immune tumor microenvironment (iTME)
 • Prioritize composition, neighborhood enrichment, and niche analysis across CLR vs DII
```

```
B) Kaposi's Sarcoma

We are studying KS samples across four stages: control to patch to plaque to nodular.

Dataset notes:
 • Use stage order: control, patch, plaque, nodular (always)
 • Tumor Cells in cell_types are KSHV+ LECs; LECs are uninfected
 • Define infected cells as those expressing any gene starting with KSHV

Visualization:
 • Keep cell type and niche colors consistent across all figures
```

Goals:
- Compare cell type composition and spatial dynamics across stages
- Quantify KSHV tropism and infection-associated spatial remodeling
- Identify niches and their remodeling across progression
- Find spatial biomarkers distinguishing stages

C) HGSOC (Pre vs Post Chemotherapy)

We are studying High-Grade Serous Ovarian Cancer (HGSOC) using paired 10x Visium spatial transcriptomics data collected before and after chemotherapy. HGSOC tumors contain malignant epithelial cells, immune populations, endothelial cells, and diverse cancer-associated fibroblast (CAF) states within a complex extracellular matrix (ECM).

Dataset notes:
- Samples are paired by patient; use treatment_label column consistently for pre/post chemotherapy annotation
- CAF-S1 programs identified via canonical markers: FAP, ACTA2, ANTXR1, ITGB1, FSP1
- ECM-myCAF defined as ANTXR1+ with high FAP and ACTA2 (SMA); iCAF defined as ANTXR1$^-$ with lower FAP/SMA
- CAF subtypes of interest: CAF-S1, ECM-myCAF, iCAF, CAF-S4
- CD8+ T-cell enrichment assessed using established cytotoxic T-cell markers with thresholds suitable for Visium resolution
- Mesenchymal subtype associated with poor prognosis and myofibroblastic CAF enrichment

Domain knowledge:
- The CAF-S1 compartment is heterogeneous, comprising immunosuppressive ANTXR1+ ECM-myCAF and ANTXR1$^-$ inflammatory CAF (iCAF)
- How CAF-S1 programs respond to chemotherapy, reorganize spatially, and relate to CD8+ T-cell enrichment remains unclear

Visualization:
- Keep CAF subtype and immune cell colors consistent across all figures
- Maintain consistent spatial plot styling for pre vs post chemotherapy comparisons

Goals:
- Quantify changes in CAF-S1-associated transcriptional programs (ECM-myCAF vs iCAF) after chemotherapy
- Characterize spatial distributions of CAF subtypes pre and post treatment
- Assess spatial relationship between CAF-S1 programs and CD8+ T-cell-enriched regions
- Explore how ECM-myCAF-associated signaling contributes to immunosuppressive stromal niches
- Identify spatially associated biomarkers and pathways reflecting chemotherapy-driven stromal and immune remodeling

#### Supplementary Note 2.1 Tips for effective sub-agent interaction

1. **Be explicit about conventions:** Do not assume the agent knows field-specific terminology.
2. **Define abbreviations:** Spell out acronyms on first use (e.g., iTME = immune tumor microenvironment).
3. **Specify preferred outputs:** Indicate if you want statistical tests, specific plot types, or particular file formats.
4. **Iterate and refine:** Start with focused questions before complex multi-part analyses.
5. **Request suggestions:** Ask the agent to propose relevant analyses based on your stated goals.
