## Supplementary Note 3 for "spatiAlytica: Viewer-Grounded Multimodal Agentic System for Interactive Spatial Omics Analysis"

### Supplementary Note 3 Planning Multi-hop QA

**Question:** What are the available cell types and stages?

**Execution Plan:**
- Step 1: GetColumnValues
- Step 2: GetColumnValues

**Progress:**
- Step 1/2: Executing GetColumnValues...
  Step 1 Completed. Result: Column `cell_type` unique values (12 total):
  `['T-cells', 'Pericytes', 'Macrophages', 'B-cells', 'Lymphatic Endothelial Cells', 'Fibroblasts', 'Vascular Endothelial Cells', 'Dendritic cells', 'Keratinocytes', 'Melanocytes', 'Pilosebaceous Cells', 'Spinous to Granular Cells']`
- Step 2/2: Executing GetColumnValues...
  Step 2 Completed. Result: Column `stage` unique values (4 total):
  `['control', 'patch', 'plaque', 'nodular']`

**Tool Sequence Execution Complete!**
Total Steps: 2
Tools Used: `GetColumnValues, GetColumnValues`

**Final Results:**
- Step 1 (`GetColumnValues`): Column `cell_type` unique values (12 total)
- Step 2 (`GetColumnValues`): Column `stage` unique values (4 total)

**spatiAlytica Summary**

**Available Cell Types and Stages**
**Cell Types (12 total):**
- B-cells
- Dendritic cells
- Fibroblasts
- Keratinocytes
- Lymphatic Endothelial Cells
- Macrophages
- Melanocytes
- Pericytes
- Pilosebaceous Cells
- Spinous to Granular Cells
- T-cells
- Vascular Endothelial Cells

**Stages (4 total):**
- Control
- Patch
- Plaque
- Nodular

You can analyze or visualize data based on any of these cell types or stages. Let me know if you want specific plots or comparisons!
