## Supplementary Note 4 for "spatiAlytica: Viewer-Grounded Multimodal Agentic System for Interactive Spatial Omics Analysis"

### Supplementary Note 4 Failure mode analysis

We systematically categorize failure modes observed during benchmark evaluation to guide future improvements. Failures are grouped into five categories.

***Code generation failures.***

The most common failure class involves incorrect or incomplete code generation: (i) incorrect API usage, where the agent selects the right analytical function but passes incompatible parameters or omits required arguments; (ii) library version mismatches, where the agent generates code targeting a different API version than installed; and (iii) incomplete multi-step computations, where intermediate preprocessing steps (e.g., computing neighbors before Leiden clustering) are omitted. The code debugger sub-agent recovers from a subset of these failures through iterative error correction (up to three retries), but complex multi-step dependency errors often exceed the debugger's repair capacity.

***Data resolution failures.***

The data resolver sub-agent occasionally fails to map user-specified biological concepts to dataset columns, particularly when column names are ambiguous (e.g., multiple columns containing "type" or "cluster"), when datasets use non-standard naming conventions, or when the user query references a concept not directly represented in the metadata. These failures propagate downstream, causing subsequent tool calls to operate on incorrect data fields.

***Spatial VQA failures.***

The spatial VQA sub-agent exhibits reduced accuracy on dense spatial plots with overlapping cell populations, small or crowded tissue regions where individual cells are difficult to distinguish, and plots with many legend categories (>15 cell types). In these cases, the agent may misidentify dominant cell types, overlook minority populations, or hallucinate spatial relationships not supported by the visualization. Performance is also sensitive to the current zoom level and field of view.

***Multi-turn reasoning failures.***

In multi-turn conversations, failures arise from (i) context loss, where relevant information from earlier turns is not retrieved from the memory sub-system; (ii) incorrect follow-up interpretation, where the agent misidentifies which prior result a follow-up question refers to; and (iii) state inconsistency, where the agent's internal model of the current `AnnData` state diverges from the actual state due to accumulated modifications. These failures are more pronounced in longer conversations (>5 turns) and when multiple analytical branches are explored within a single session.

***Hallucination and overinterpretation.***

As with all LLM-based systems, spatiAlytica can generate plausible but unsupported biological interpretations, particularly during IR steps. Observed cases include citing gene functions not supported by the computed differential expression results, inferring causal relationships from correlative spatial patterns, and overinterpreting small effect sizes. The system mitigates this risk by grounding interpretations in explicit tool outputs and executed code, but users should independently validate key biological claims.
