## Supplementary Note 5 for "spatiAlytica: Viewer-Grounded Multimodal Agentic System for Interactive Spatial Omics Analysis"

### Supplementary Note 5 spatiAlytica recovers chemotherapy-driven CAF remodeling in Visium-profiled HGSOC

To test cross-platform generalization, we applied spatiAlytica to a 10-section Visium HGSOC cohort (4 treatment-naive and 6 post-chemotherapy sections from 8 HGSOC patients) [44]. Natural-language queries comparing CAF-S1-enriched and $CD8^+$-enriched spot proportions (EA1; Fig. 4a) showed CAF-S1 contraction post-chemotherapy (29.7% → 22.3%) with a reciprocal $CD8^+$ T-cell expansion. Examining *ANTXR1* expression within CAF-S1 spots (EA2; Fig. 4b) revealed a post-treatment shift toward lower *ANTXR1* levels, and a follow-up quantification query (EA3; Fig. 4c) showed the $ANTXR1^+$ (ECM-myCAF) fraction within CAF-S1 dropping from 90.9% pre-chemo to 82.5% post-chemo. The user response sub-agent's synthesis (IR1; Fig. 4d) recovered the published model of chemotherapy selectively attenuating the ECM-myCAF program [44].

When asked whether $ANTXR1^+$ CAF-S1 and CD8-enriched spots changed their spatial relationship after treatment, spatiAlytica routed the query to the code writer sub-agent, which generated and executed a permutation-based neighborhood-enrichment analysis (EA4; Fig. 4e) showing the ECM-myCAF↔CD8 cross-pair $Z$-score becoming more strongly negative post-chemotherapy ($-2.05 \rightarrow -5.63$), consistent with stronger spatial avoidance of $CD8^+$ T-cells from ECM-myCAF neighborhoods (IR2; Fig. 4f). A follow-up DE query within $ANTXR1^+$ CAF-S1 spots (EA5; Fig. 4g) recovered downregulation of matrix and ECM-program genes (*COL1A2*, *COL1A1*, *FN1*, *SPARC*, *MMP11*, *LUM*) with concurrent enrichment of immunoglobulin and complement transcripts (*IGKC*, *IGHG1*, *IGHA1*, *C3*), consistent with chemotherapy-induced attenuation of the ECM-myCAF program [44] alongside post-chemo plasma-cell or B-cell-derived antibody activity in the local stromal niche. A final integrative synthesis (IR3; Fig. 4h) unified compartmental contraction, transcriptional reprogramming, and CD8 spatial exclusion into a coherent treatment-response model, demonstrating that spatiAlytica's multimodal interactive agentic workflow operates equivalently on spot-level Visium data as on single-cell Xenium resolution.
