## Supplementary Note 7 for "spatiAlytica: Viewer-Grounded Multimodal Agentic System for Interactive Spatial Omics Analysis"

### Supplementary Note 7 Full interaction transcript: Kaposi's sarcoma niche analysis and CD8 T-cell program

The complete interaction transcript below corresponds to the case study in §1.7. Each **Turn** block records the user prompt, the agent activity (tool selection and execution timing), and the spatiAlytica response, including any rendered figures, tables, and pointers to associated CSV result files. The viewer was held at the same field of view across all turns (no zoom or pan was applied between questions) so that every spatial plot in this transcript shares an identical spatial reference frame; per-cell point markers therefore appear small in these page-fit screenshots but render at full fidelity inside the interactive Napari canvas. Raw session artefacts (session log, generated code, telemetry, query timing, action log, and asset images) are bundled in `supplementary_data_4_ks_niches_cd8/`.

#### Turn 1 (Query 001)

**User prompt.**

Same Kaposi's sarcoma dataset as before (control, patch, plaque, nodular, in increasing severity). FYI, the cell-type labels in this dataset already follow this convention: cells labeled "Tumor Cell" are the KSHV-positive lymphatic endothelial cells, and cells labeled "Lymphatic Endothelial Cells" are the uninfected LEC population (no relabeling needed, just so you know what they mean). A cell counts as KSHV-infected if it expresses any gene whose name starts with "KSHV". We're going to dig into cellular niches, how infection spreads through them, and the CD8 T-cell program across the four stages. Please use the stage order control, patch, plaque, nodular in every result, and keep niche and cell-type colors stable across plots. Ready?

---

**Agent activity.**

*Selected tool:* ***`RespondWithTextTool`***

- ▷ Analyzing your question...
- ▷ Preparing context from memory...
- ▷ Consulting spatiAlytica Core...
- ▷ Generating response...
- ▷ [OK] Response ready (5.8s)
- ▷ Selected action: Response Generation

---

**spatiAlytica response.**

Yes—ready.

I'll treat **stage** as ordered: **control** → **patch** → **plaque** → **nodular** for every summary/plot. For stable colors, I'll reuse the existing **cell_type** palette already in the viewer, and for niches we'll keep a fixed palette once we create the niche labels.

For KSHV infection status, I'll define a per-cell flag **infected = any expressed gene with prefix `KSHV`** (and we can also summarize infection burden per stage / per niche).

What do you want to start with?

1) Run **spatial niche analysis** (choose k neighbors + #niches), then map **infection rate per niche** across stages
2) Focus on **CD8 T-cells**: stage-wise DE / programs and spatial localization relative to Tumor Cell niches
3) First, a quick **spatial overview**: cell types + Tumor Cell vs uninfected LECs per stage

---

#### Turn 2 (Query 002)

**User prompt.**

Let's start by computing 10 cellular niches based on cell-type composition, using a small neighborhood of 30 cells.

---

**Agent activity.**

*Selected tool:* ***`SpatialPlotTool`***

- ▷ Analyzing your question...
- ▷ Preparing context from memory...
- ▷ Consulting spatiAlytica Core...
- ▷ Starting niche analysis...
- ▷ Starting niche analysis...

▷ Computing 30 nearest neighbors per-sample (sample_id) for 301,689 cells...
▷ [OK] Per-sample nearest neighbor search complete
▷ Performing K-means clustering to identify 10 niches...
▷ [OK] Clustering complete
▷ Creating niche plot: Niches (k=30, n=10)...
▷ [OK] Niche analysis complete
▷ [OK] Niche analysis complete (27.6s)
▷ Creating spatial visualization...
▷ Rendering spatial visualization...
▷ [OK] Spatial plot created (1.1s)
▷ Selected action: Spatial Visualization

**spatiAlytica response.**

##### Cellular niche analysis (10 niches; neighborhood size = 30 cells)

##### Summary table (composition-based niches)

| Cluster | Niche name | Total cells | Top 3 cell types (%) |
|---|---|---|---|
| 0 | Fibroblast Enriched | 45875 | Fibroblasts (47.14%), Macrophages (19.95%), Tumor Cell (5.81%) |
| 1 | Tumor Core | 43851 | Tumor Cell (61.14%), Fibroblasts (14.10%), Lymphatic Endothelial Cells (8.93%) |
| 2 | Keratinocyte Enriched | 14631 | Keratinocytes (55.20%), Spinous to Granular Cells (17.19%), Pilosebaceous Cells (12.77%) |
| 3 | Macrophage Enriched | 10169 | Macrophages (84.67%), T-cells (3.00%), CD4 T-cells (2.36%) |
| 4 | Tumor Stroma | 41108 | Tumor Cell (27.70%), Fibroblasts (25.09%), Vascular Endothelial Cells (27.70%) |
| 5 | Tumor Boundary | 46520 | Tumor Cell (30.39%), Fibroblasts (25.64%), Macrophages (13.10%) |
| 6 | Macrophage-CD8 | 28824 | Macrophages (20.30%), Fibroblasts (17.14%), CD8 T-cells (13.99%) |
| 7 | Macrophage Enriched II | 32688 | Macrophages (37.80%), Fibroblasts (25.18%), T-cells (7.32%) |
| 8 | Tumor Core II | 32739 | Tumor Cell (43.32%), Lymphatic Endothelial Cells (17.34%), Fibroblasts (13.76%) |
| 9 | Pericyte Enriched | 5284 | Pericytes (54.49%), Fibroblasts (19.28%), Vascular Endothelial Cells (17.71%) |

##### Brief biological descriptions (as computed)

- **Niche 0: Fibroblast Enriched** — stromal-heavy with substantial myeloid presence.
- **Niche 1: Tumor Core** — tumor-dominant with stromal and lymphatic endothelial admixture.
- **Niche 2: Keratinocyte Enriched** — epithelial/epidermal compartment with stratified differentiation components.
- **Niche 3: Macrophage Enriched** — strongly myeloid-dominant, consistent with dense macrophage aggregation.
- **Niche 4: Tumor Stroma** — mixed tumor–stroma niche with substantial fibroblast and macrophage content.
- **Niche 5: Tumor Boundary** — tumor–stroma interface with appreciable immune infiltration.
- **Niche 6: Macrophage-CD8** — mixed immune–stromal niche where macrophages and CD8 T cells co-occur.
- **Niche 7: Macrophage Enriched II** — macrophage-dominant with strong fibroblast contribution.
- **Niche 8: Tumor Core II** — tumor-rich with prominent lymphatic endothelial representation.
- **Niche 9: Pericyte Enriched** — perivascular niche with high pericyte content plus endothelial and stromal cells.

##### Spatial visualization

A spatial plot of these niches was created successfully and is displayed in the **main interactive Napari viewer window**.

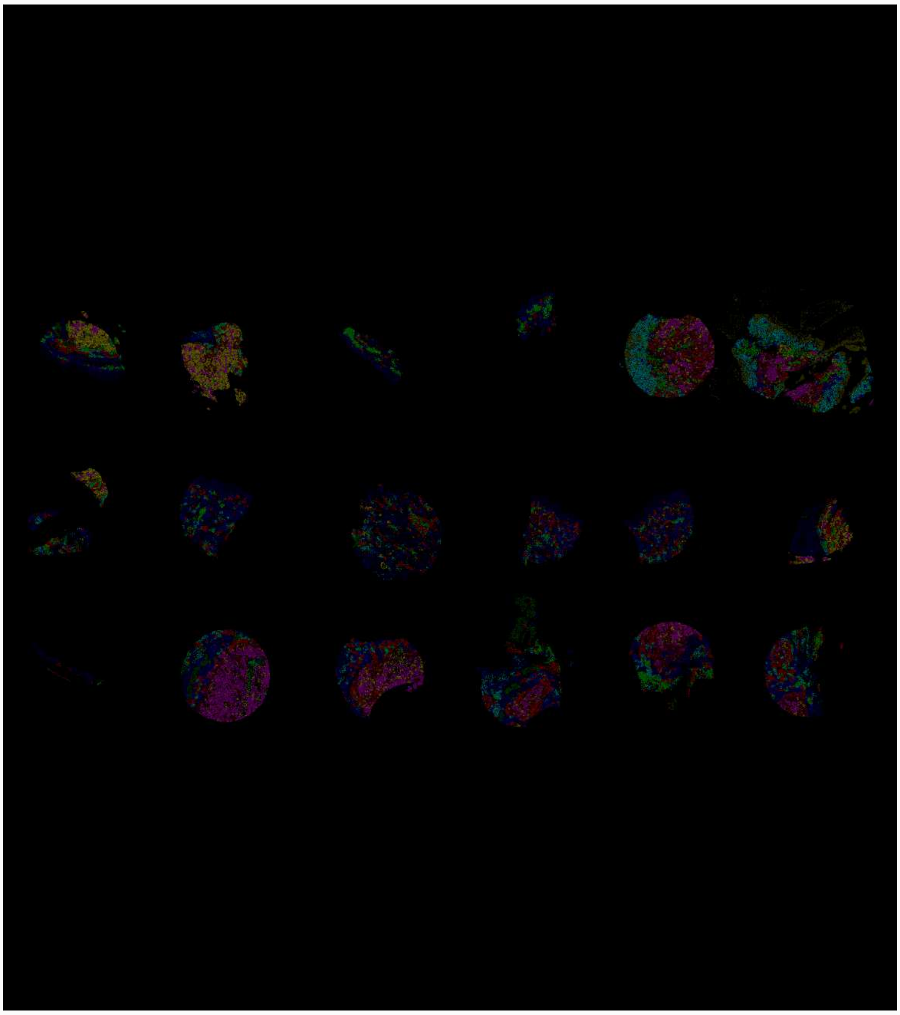

*Associated data:* `query_002_analysis_results.csv`

#### Turn 3 (Query 003)

**User prompt.**

Show me a stacked bar plot of the cell-type composition for each niche, so we can see what each niche is dominantly made of.

**Agent activity.**

*Selected tool:* `WriteCodeTool`

- ▷ Analyzing your question...
- ▷ Preparing context from memory...
- ▷ Consulting spatiAlytica Core...
- ▷ Writing and executing code...
- ▷ Formatting results...
- ▷ [OK] Code execution complete (55.5s)
- ▷ Selected action: Code Execution

**spatiAlytica response.**

**Stacked bar plot — cell-type composition per niche**

A **stacked bar plot** (showing **percent of each cell type within each** `NICHE_names_30_10` **niche**) has been created.

**Cell-type composition (percent) within each niche**

**Fibroblast Enriched**

| Cell type | Percent | Count |
|---|---|---|
| Fibroblasts | 47.1411 | 21626 |
| Macrophages | 19.9499 | 9152 |
| Tumor Cell | 5.81362 | 2667 |
| Vascular Endothelial Cells | 5.5673 | 2554 |
| T-cells | 4.90027 | 2248 |
| CD8 T-cells | 4.59728 | 2109 |
| Lymphatic Endothelial Cells | 3.58583 | 1645 |
| CD4 T-cells | 3.08665 | 1416 |
| Pericytes | 2.55695 | 1173 |
| Dendritic cells | 1.86158 | 854 |
| B-cells | 0.710627 | 326 |
| Pilosebaceous Cells | 0.093733 | 43 |
| Melanocytes | 0.0762943 | 35 |
| Keratinocytes | 0.0544959 | 25 |
| Spinous to Granular Cells | 0.00435967 | 2 |

**Keratinocyte Enriched**

| Cell type | Percent | Count |
|---|---|---|
| Keratinocytes | 55.1979 | 8076 |
| Spinous to Granular Cells | 17.1895 | 2515 |
| Pilosebaceous Cells | 12.7742 | 1869 |
| Melanocytes | 6.66393 | 975 |
| Fibroblasts | 4.59299 | 672 |
| Macrophages | 1.38746 | 203 |
| Vascular Endothelial Cells | 0.88169 | 129 |
| CD8 T-cells | 0.375914 | 55 |
| Lymphatic Endothelial Cells | 0.239218 | 35 |
| T-cells | 0.225548 | 33 |
| CD4 T-cells | 0.18454 | 27 |
| Pericytes | 0.150366 | 22 |
| Dendritic cells | 0.109357 | 16 |
| Tumor Cell | 0.0273392 | 4 |

**Macrophage Enriched**

| Cell type | Percent | Count |
|---|---|---|
| Macrophages | 84.6691 | 8610 |
| Fibroblasts | 5.54627 | 564 |
| T-cells | 2.99931 | 305 |
| CD4 T-cells | 2.36011 | 240 |
| CD8 T-cells | 1.22923 | 125 |
| B-cells | 0.678533 | 69 |
| Tumor Cell | 0.61953 | 63 |
| Dendritic cells | 0.599862 | 61 |
| Pericytes | 0.452355 | 46 |
| Vascular Endothelial Cells | 0.344183 | 35 |
| Lymphatic Endothelial Cells | 0.275347 | 28 |
| Pilosebaceous Cells | 0.137673 | 14 |
| Melanocytes | 0.0590029 | 6 |
| Keratinocytes | 0.0295014 | 3 |

**Macrophage Enriched II**

| Cell type | Percent | Count |
|---|---|---|
| Macrophages | 37.7998 | 12356 |
| Fibroblasts | 25.1836 | 8232 |
| T-cells | 7.32073 | 2393 |
| CD8 T-cells | 6.19493 | 2025 |
| CD4 T-cells | 5.46378 | 1786 |
| Vascular Endothelial Cells | 4.94677 | 1617 |
| Dendritic cells | 3.48752 | 1140 |
| Tumor Cell | 3.16936 | 1036 |
| Pericytes | 2.52692 | 826 |
| Lymphatic Endothelial Cells | 2.02827 | 663 |
| B-cells | 1.74682 | 571 |
| Pilosebaceous Cells | 0.067303 | 22 |
| Melanocytes | 0.0581253 | 19 |
| Keratinocytes | 0.00611845 | 2 |

**Macrophage-CD8**

| Cell type | Percent | Count |
|---|---|---|
| Macrophages | 20.2956 | 5850 |
| Fibroblasts | 17.142 | 4941 |
| CD8 T-cells | 13.9883 | 4032 |
| T-cells | 10.6959 | 3083 |
| CD4 T-cells | 8.49639 | 2449 |
| Dendritic cells | 6.62295 | 1909 |
| Vascular Endothelial Cells | 6.51193 | 1877 |
| Tumor Cell | 3.88912 | 1121 |
| Pericytes | 3.75729 | 1083 |
| Lymphatic Endothelial Cells | 3.17097 | 914 |
| B-cells | 2.20303 | 635 |
| Melanocytes | 2.04691 | 590 |
| Pilosebaceous Cells | 0.988759 | 285 |
| Keratinocytes | 0.180405 | 52 |
| Spinous to Granular Cells | 0.010408 | 3 |

**Pericyte Enriched**

| Cell type | Percent | Count |
|---|---|---|
| Pericytes | 54.4852 | 2879 |
| Fibroblasts | 19.2846 | 1019 |
| Vascular Endothelial Cells | 17.7139 | 936 |
| Macrophages | 4.61771 | 244 |
| T-cells | 1.23013 | 65 |
| CD8 T-cells | 0.889478 | 47 |
| CD4 T-cells | 0.454201 | 24 |
| Lymphatic Endothelial Cells | 0.454201 | 24 |
| Tumor Cell | 0.302801 | 16 |
| Dendritic cells | 0.283876 | 15 |
| B-cells | 0.11355 | 6 |
| Pilosebaceous Cells | 0.0946253 | 5 |
| Melanocytes | 0.0757002 | 4 |

**Tumor Boundary**

| Cell type | Percent | Count |
| --- | --- | --- |
| Tumor Cell | 30.3912 | 14138 |
| Fibroblasts | 25.6449 | 11930 |
| Macrophages | 13.104 | 6096 |
| Vascular Endothelial Cells | 10.0752 | 4687 |
| Lymphatic Endothelial Cells | 7.31943 | 3405 |
| Pericytes | 5.47291 | 2546 |
| T-cells | 3.62855 | 1688 |
| CD8 T-cells | 2.22485 | 1035 |
| Dendritic cells | 1.23173 | 573 |
| CD4 T-cells | 0.803955 | 374 |
| B-cells | 0.0687876 | 32 |
| Pilosebaceous Cells | 0.0343938 | 16 |

**Tumor Core**

| Cell type | Percent | Count |
| --- | --- | --- |
| Tumor Cell | 61.1411 | 26811 |
| Fibroblasts | 14.0955 | 6181 |
| Lymphatic Endothelial Cells | 8.92568 | 3914 |
| Macrophages | 6.87328 | 3014 |
| Pericytes | 3.01019 | 1320 |
| Vascular Endothelial Cells | 2.78671 | 1222 |
| T-cells | 2.14134 | 939 |
| CD8 T-cells | 0.499418 | 219 |
| Dendritic cells | 0.369433 | 162 |
| CD4 T-cells | 0.15279 | 67 |
| B-cells | 0.00228045 | 1 |
| Pilosebaceous Cells | 0.00228045 | 1 |

**Tumor Core II**

| Cell type | Percent | Count |
| --- | --- | --- |
| Tumor Cell | 43.3184 | 14182 |
| Lymphatic Endothelial Cells | 17.3432 | 5678 |
| Fibroblasts | 13.7634 | 4506 |
| Macrophages | 10.6845 | 3498 |
| Vascular Endothelial Cells | 4.487 | 1469 |
| Pericytes | 3.7631 | 1232 |
| T-cells | 3.63175 | 1189 |
| CD8 T-cells | 1.29509 | 424 |
| Dendritic cells | 1.25844 | 412 |
| CD4 T-cells | 0.351263 | 115 |
| Pilosebaceous Cells | 0.0458169 | 15 |
| B-cells | 0.0427625 | 14 |
| Keratinocytes | 0.00916338 | 3 |
| Melanocytes | 0.00610892 | 2 |

**Tumor Stroma**

| Cell type | Percent | Count |
|---|---|---|
| Vascular Endothelial Cells | 27.6978 | 11386 |
| Fibroblasts | 25.0851 | 10312 |
| Macrophages | 15.6442 | 6431 |
| Pericytes | 9.23178 | 3795 |
| Tumor Cell | 4.78009 | 1965 |
| T-cells | 4.70711 | 1935 |
| CD8 T-cells | 4.03571 | 1659 |
| Dendritic cells | 2.47884 | 1019 |
| CD4 T-cells | 2.32072 | 954 |
| Lymphatic Endothelial Cells | 2.28179 | 938 |
| Melanocytes | 0.615452 | 253 |
| B-cells | 0.540041 | 222 |
| Pilosebaceous Cells | 0.442736 | 182 |
| Keratinocytes | 0.128929 | 53 |
| Spinous to Granular Cells | 0.00973047 | 4 |

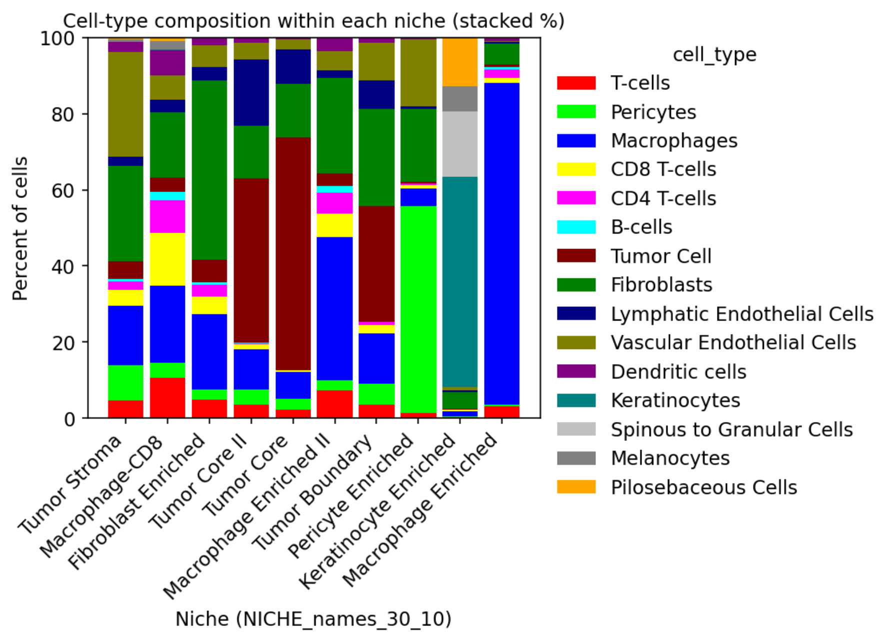


#### Turn 4 (Query 004)

**User prompt.**

Now let's look at how niche composition shifts across the four stages: what fraction of each stage is made up of each niche? Stacked bar of niche % across stages, please, with a quick interpretation.

**Agent activity.**

*Selected tool:* *WriteCodeTool*

- ▷ Analyzing your question...
- ▷ Preparing context from memory...
- ▷ Consulting spatiAlytica Core...
- ▷ Writing and executing code...
- ▷ Formatting results...
- ▷ [OK] Code execution complete (23.2s)
- ▷ Writing and executing code...
- ▷ Formatting results...
- ▷ [OK] Code execution complete (51.9s)
- ▷ Selected action: Code Execution

**spatiAlytica response.**

**Niche composition shifts across stages (fraction of each stage made up of each niche)**

A **stacked bar plot of niche composition (%) across the four stages** has been created and is visible in the **Visualization Tab**.

**Stage × niche percent composition (%)**

| Stage | Niche | Percent |
|---|---|---|
| control | Keratinocyte Enriched | 75 |
| control | Tumor Stroma | 13.0245 |
| control | Fibroblast Enriched | 6.03147 |
| control | Macrophage-CD8 | 3.05944 |
| control | Pericyte Enriched | 2.88462 |
| control | Macrophage Enriched | 0 |
| control | Macrophage Enriched II | 0 |
| control | Tumor Boundary | 0 |
| control | Tumor Core | 0 |
| control | Tumor Core II | 0 |
| patch | Macrophage-CD8 | 37.5669 |
| patch | Keratinocyte Enriched | 25.1292 |
| patch | Fibroblast Enriched | 21.3997 |
| patch | Tumor Stroma | 9.24378 |
| patch | Tumor Boundary | 3.63551 |
| patch | Macrophage Enriched II | 2.75247 |
| patch | Tumor Core II | 0.263034 |
| patch | Macrophage Enriched | 0.00939408 |
| patch | Pericyte Enriched | 0 |
| patch | Tumor Core | 0 |
| plaque | Fibroblast Enriched | 29.7756 |
| plaque | Tumor Stroma | 17.2616 |
| plaque | Tumor Boundary | 16.1009 |
| plaque | Keratinocyte Enriched | 10.9102 |
| plaque | Macrophage-CD8 | 9.74605 |
| plaque | Macrophage Enriched II | 9.6808 |
| plaque | Tumor Core II | 4.68415 |
| plaque | Tumor Core | 1.11437 |
| plaque | Pericyte Enriched | 0.679957 |
| plaque | Macrophage Enriched | 0.0463607 |
| nodular | Tumor Core | 18.6488 |
| nodular | Tumor Boundary | 15.8663 |
| nodular | Tumor Core II | 12.9426 |
| nodular | Tumor Stroma | 12.9163 |
| nodular | Macrophage Enriched II | 11.5501 |
| nodular | Fibroblast Enriched | 11.304 |
| nodular | Macrophage-CD8 | 8.25085 |
| nodular | Macrophage Enriched | 4.37752 |
| nodular | Pericyte Enriched | 2.09573 |
| nodular | Keratinocyte Enriched | 2.04782 |

**Top 3 niches per stage**

| Stage | Niche | Percent |
|---|---|---|
| control | Keratinocyte Enriched | 75 |
| control | Tumor Stroma | 13.0245 |
| control | Fibroblast Enriched | 6.03147 |
| patch | Macrophage-CD8 | 37.5669 |
| patch | Keratinocyte Enriched | 25.1292 |
| patch | Fibroblast Enriched | 21.3997 |
| plaque | Fibroblast Enriched | 29.7756 |
| plaque | Tumor Stroma | 17.2616 |
| plaque | Tumor Boundary | 16.1009 |
| nodular | Tumor Core | 18.6488 |
| nodular | Tumor Boundary | 15.8663 |
| nodular | Tumor Core II | 12.9426 |

**Quick interpretation (composition shifts)**

- **Control** is dominated by **Keratinocyte Enriched (75%)**, with smaller contributions from **Tumor Stroma (13.0245%)** and **Fibroblast Enriched (6.03147%)**; tumor-associated niches (**Tumor Boundary/Core/Core II**) are **0%**.
- **Patch** shifts strongly toward **Macrophage-CD8 (37.5669%)**, while **Keratinocyte Enriched (25.1292%)** and **Fibroblast Enriched (21.3997%)** remain substantial; **Tumor Boundary** appears (**3.63551%**) and **Tumor Core II** is present at low fraction (**0.263034%**).
- **Plaque** becomes more distributed across stromal/boundary niches: **Fibroblast Enriched (29.7756%)**, **Tumor Stroma (17.2616%)**, and **Tumor Boundary (16.1009%)**, with added **Macrophage-CD8 (9.74605%)** and **Macrophage Enriched II (9.6808%)**; **Tumor Core/Core II** begin to appear (**1.11437% / 4.68415%**).
- **Nodular** shows the strongest tumor-associated composition, led by **Tumor Core (18.6488%)** and **Tumor Boundary (15.8663%)**, with substantial **Tumor Core II (12.9426%)** and **Tumor Stroma (12.9163%)**, alongside **Macrophage Enriched II (11.5501%)** and **Fibroblast Enriched (11.304%)**.

---

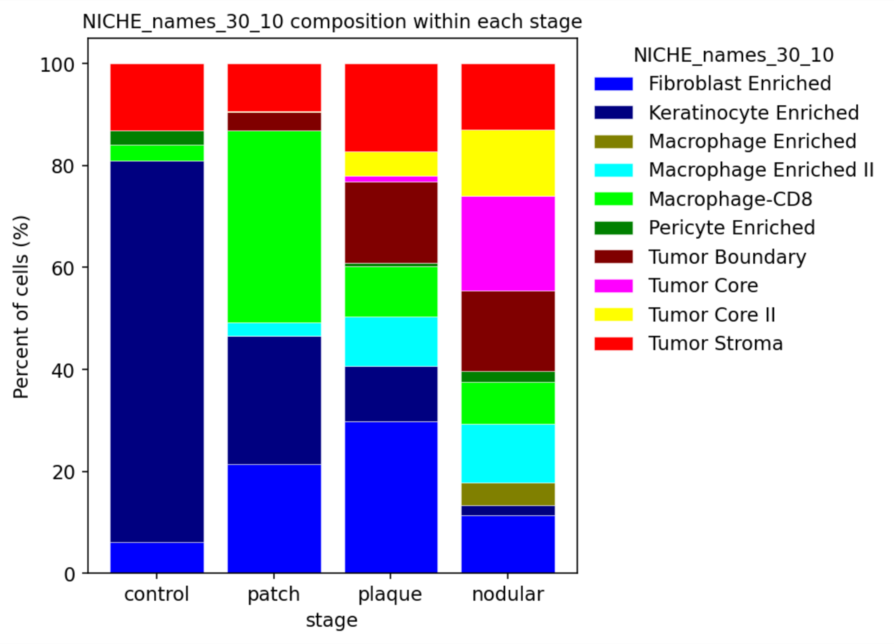

#### Turn 5 (Query 005)

**User prompt.**

Looking at the spatial niche plot across the cores together with the niche-by-stage percentages you just made, how do the niches remodel as the disease progresses?

**Agent activity.**

*Selected tool:* `RespondWithTextTool`

▷ Analyzing your question...
▷ Preparing context from memory...
▷ Consulting spatiAlytica Core...
▷ Generating response...
▷ [OK] Response ready (16.1s)
▷ Selected action: Response Generation

**spatiAlytica response.**

**How niches remodel with disease progression (control → patch → plaque → nodular)**

Integrating the **spatial niche map across TMA cores** with the **stage × niche composition table** you generated, the disease shows a clear transition from **epidermal/normal architecture** toward **immune-infiltrated**, then **stromal/boundary expansion**, and finally **tumor-core–dominated** microenvironments.

**1) Control: largely non-tumor, keratinocyte-dominant**

- **Keratinocyte Enriched: 75%** (dominant "baseline" epithelial niche)
- **Tumor Stroma: 13.0%**, **Fibroblast Enriched: 6.0%** (supporting tissue compartments)
- **Tumor Boundary/Core/Core II: 0%** (no appreciable tumor-structured niches)

**Spatially across cores**, this corresponds to cores being visually dominated by keratinocyte-rich territories with limited immune/tumor structuring.

**2) Patch: immune recruitment becomes a major organizing force**

- **Macrophage-CD8: 37.6%** becomes the largest niche
- **Keratinocyte Enriched: 25.1%** remains substantial
- **Fibroblast Enriched: 21.4%** increases
- **Tumor Boundary appears: 3.6%**; **Tumor Core II: 0.26%** (early/rare tumor-like neighborhood structure)

**Interpretation:** patch lesions look like an **immune-activated / inflammatory remodeling phase**, where immune neighborhoods (especially macrophage–CD8-associated) organize large contiguous regions, but a sizable keratinocyte compartment persists.

**3) Plaque: expansion of stromal remodeling and "interface" (boundary) niches**

- **Fibroblast Enriched: 29.8%** (largest)
- **Tumor Stroma: 17.3%** and **Tumor Boundary: 16.1%** rise strongly
- **Macrophage-CD8: 9.7%** and **Macrophage Enriched II: 9.7%** persist
- **Tumor Core II: 4.68%** and **Tumor Core: 1.11%** begin to emerge

**Spatially**, plaque is where you expect to see more **segmentation into compartments**: broad stromal regions and widened boundary-like interfaces, consistent with a growing lesion architecture rather than diffuse inflammation alone.

**4) Nodular: tumor architecture dominates and diversifies**

- Tumor-structured niches become the leading components:
  - **Tumor Core: 18.65%**
  - **Tumor Boundary: 15.87%**
  - **Tumor Core II: 12.94%**
  - plus **Tumor Stroma: 12.92%**
- Non-tumor/supportive and immune niches remain major but are no longer dominant:
  - **Macrophage Enriched II: 11.55%**, **Fibroblast Enriched: 11.30%**, **Macrophage-CD8: 8.25%**

- **Keratinocyte Enriched drops to 2.05%**

**Interpretation:** nodular disease shows a clear **shift to organized tumor microenvironments**—core regions, boundary/interface regions, and stromal supports—indicating that the lesion is no longer primarily "inflammatory remodeling" but is **tumor-architecture remodeling**.

**Overall remodeling trajectory (one-sentence summary)**

Across progression, the tissue remodels from **keratinocyte-rich normal-like neighborhoods (control) → immune-dominant neighborhoods (patch) → stromal + boundary/interface expansion (plaque) → tumor core/boundary/stroma compartmentalization (nodular)**, aligning with your goal of understanding how the spatial microenvironment reorganizes with stage.

---

*Associated data:* `query_005_analysis_results.csv`

#### Turn 6 (Query 006)

**User prompt.**

Could you also give me the niche-by-stage breakdown as a percentage table, in the control → patch → plaque → nodular stage order, so I have the actual numbers?

**Agent activity.**

*Selected tool: `WriteCodeTool`*

▷ Analyzing your question...
▷ Preparing context from memory...
▷ Consulting spatiAlytica Core...
▷ Writing and executing code...
▷ Formatting results...
▷ [OK] Code execution complete (46.7s)
▷ Selected action: Code Execution

**spatiAlytica response.**

**Niche-by-stage breakdown (%), ordered: control → patch → plaque → nodular**

| Stage | Fibroblast Enriched (%) | Keratinocyte Enriched (%) | Macrophage Enriched (%) | Macrophage Enriched II (%) | Macrophage-CD8 (%) | Pericyte Enriched (%) | Tumor Boundary (%) | Tumor Core (%) | Tumor Core II (%) | Tumor Stroma (%) |
|---|---|---|---|---|---|---|---|---|---|---|
| control | 6.03 | 75.00 | 0.00 | 0.00 | 3.06 | 2.88 | 0.00 | 0.00 | 0.00 | 13.02 |
| patch | 21.40 | 25.13 | 0.01 | 2.75 | 37.57 | 0.00 | 3.64 | 0.00 | 0.26 | 9.24 |
| plaque | 29.78 | 10.91 | 0.05 | 9.68 | 9.75 | 0.68 | 16.10 | 1.11 | 4.68 | 17.26 |
| nodular | 11.30 | 2.05 | 4.38 | 11.55 | 8.25 | 2.10 | 15.87 | 18.65 | 12.94 | 12.92 |

---

*Associated data:* `query_006_analysis_results.csv`

#### Turn 7 (Query 007)

**User prompt.**

Let's bring infection back in. Mark every cell as KSHV-infected or uninfected based on whether it expresses any KSHV-named gene, and we'll use that going forward. Show me a spatial map with infected cells in red and uninfected cells in grey.

**Agent activity.**

*Selected tool: `WriteCodeTool`*

▷ Analyzing your question...
▷ Preparing context from memory...
▷ Consulting spatiAlytica Core...
▷ Writing and executing code...
▷ Formatting results...
▷ [OK] Code execution complete (19.6s)
▷ Creating spatial visualization...
▷ Rendering spatial visualization...
▷ [OK] Spatial plot created (1.2s)
▷ Selected action: Code Execution

**spatiAlytica response.**

**KSHV infection status (used going forward)**

Each cell was labeled **KSHV-infected** if it expresses **any** of these **9 KSHV genes**:

| KSHV gene |
|---|
| KSHV.K2 |
| KSHV.K9 |
| KSHV.ORF50 |
| KSHV.ORF57 |
| KSHV.ORF59 |
| KSHV.ORF65 |
| KSHV.ORF71 |
| KSHV.ORF72 |
| KSHV.ORF73 |

**Counts**

| KSHV infection status | Count |
|---|---|
| uninfected | 217,377 |
| infected | 84,312 |

**Spatial map (infected vs uninfected)**

A **spatial plot was created successfully** and is displayed in the **main interactive Napari viewer window**, with:

- **infected cells = red**
- **uninfected cells = grey**

---

**Turn 8 (Query 008)**

**User prompt.**

Stacked bar of infection percentage across the four stages, please.

**Agent activity.**

*Selected tool: WriteCodeTool*

- ▷ Analyzing your question...
- ▷ Preparing context from memory...
- ▷ Consulting spatiAlytica Core...
- ▷ Writing and executing code...
- ▷ Formatting results...
- ▷ [OK] Code execution complete (34.1s)
- ▷ Selected action: Code Execution

**spatiAlytica response.**

**Stacked bar — Infection percentage across the four stages**

A stacked bar plot of **KSHV infection status percentages by stage** was created.

**Stage × infection status (%)**

| Stage | Uninfected (%) | Infected (%) |
|---|---|---|
| control | 100 | 0 |
| patch | 95.6411 | 4.35885 |
| plaque | 84.2751 | 15.7249 |
| nodular | 67.7589 | 32.2411 |

**Stage × infection status (% and counts)**

| Stage | KSHV_infection_status | Percent | Count |
|---|---|---|---|
| control | uninfected | 100 | 1144 |
| control | infected | 0 | 0 |
| patch | uninfected | 95.6411 | 10181 |
| patch | infected | 4.35885 | 464 |
| plaque | uninfected | 84.2751 | 49081 |
| plaque | infected | 15.7249 | 9158 |
| nodular | uninfected | 67.7589 | 156971 |
| nodular | infected | 32.2411 | 74690 |

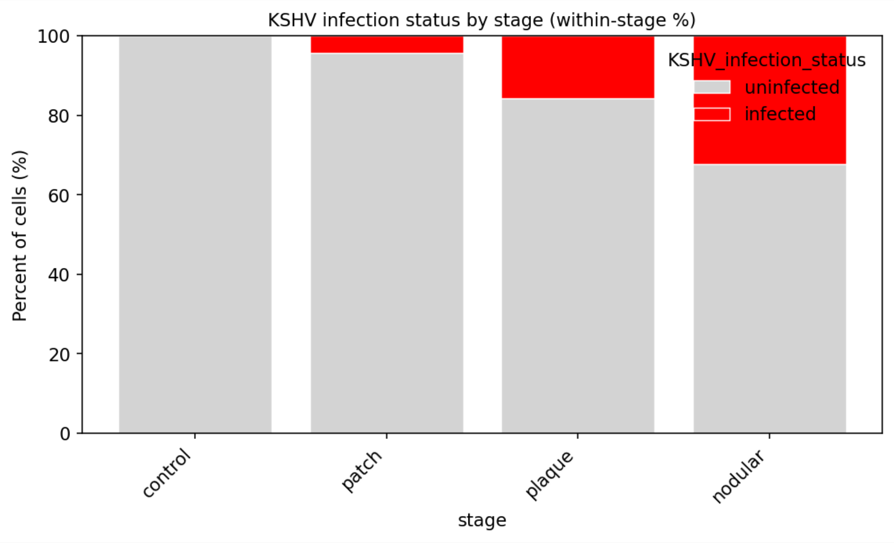


#### Turn 9 (Query 009)

**User prompt.**

What fraction of cells in each niche are infected?

**Agent activity.**

*Selected tool:* ***WriteCodeTool***

- ▷ Analyzing your question...
- ▷ Preparing context from memory...
- ▷ Consulting spatiAlytica Core...
- ▷ Writing and executing code...
- ▷ Formatting results...
- ▷ [OK] Code execution complete (35.4s)
- ▷ Selected action: Code Execution

**spatiAlytica response.**

##### Fraction of cells infected in each niche

Niche-level infection fractions (sorted by **infected fraction**, descending):

| Niche | Total cells | Infected | Uninfected | Infected fraction | Infected % | Uninfected fraction | Uninfected % |
|---|---|---|---|---|---|---|---|
| Tumor Core | 43851 | 33221 | 10630 | 0.7576 | 75.76 | 0.2424 | 24.24 |
| Tumor Core II | 32739 | 17490 | 15249 | 0.5342 | 53.42 | 0.4658 | 46.58 |
| Tumor Boundary | 46520 | 20658 | 25862 | 0.4441 | 44.41 | 0.5559 | 55.59 |
| Fibroblast Enriched | 45875 | 4517 | 41358 | 0.0985 | 9.85 | 0.9015 | 90.15 |
| Tumor Stroma | 41108 | 3976 | 37132 | 0.0967 | 9.67 | 0.9033 | 90.33 |
| Macrophage-CD8 | 28824 | 1892 | 26932 | 0.0656 | 6.56 | 0.9344 | 93.44 |
| Macrophage Enriched II | 32688 | 1955 | 30733 | 0.0598 | 5.98 | 0.9402 | 94.02 |
| Pericyte Enriched | 5284 | 179 | 5105 | 0.0339 | 3.39 | 0.9661 | 96.61 |
| Macrophage Enriched | 10169 | 276 | 9893 | 0.0271 | 2.71 | 0.9729 | 97.29 |
| Keratinocyte Enriched | 14631 | 148 | 14483 | 0.0101 | 1.01 | 0.9899 | 98.99 |

---

*Associated data:* `query_009_analysis_results.csv`

#### Turn 10 (Query 010)

**User prompt.**

Before we go deeper, please summarize what we've learned so far: niches and their dominant cell types, how niches remodel across stages, and how infection distributes across niches and stages.

**Agent activity.**

*Selected tool:* ***RespondWithTextTool***

▷ Analyzing your question...
▷ Preparing context from memory...
▷ Consulting spatiAlytica Core...
▷ Generating response...
▷ [OK] Response ready (16.3s)
▷ Selected action: Response Generation

---

**spatiAlytica response.**

**1) Niches identified (k=30 neighbors, 10 niche clusters) and their dominant cell types**

We defined spatial niches using neighborhood composition (within-sample), producing 10 named niches (`NICHE_names_30_10`). From the niche-by-cell-type composition summaries, these niches correspond to the following dominant programs:

- **Keratinocyte Enriched**: dominated by **Keratinocytes / Spinous-to-Granular** (epithelial compartment).
- **Macrophage Enriched** and **Macrophage Enriched II**: dominated by **Macrophages** (with varying mixtures of other immune/stromal cells).
- **Macrophage–CD8**: co-enrichment of **Macrophages + CD8 T-cells** (an immune-interacting neighborhood).
- **Pericyte Enriched**: dominated by **Pericytes** (vascular support).
- **Fibroblast Enriched**: dominated by **Fibroblasts** (stromal compartment).
- **Tumor Stroma**: stromal-heavy niche (Fibroblasts with additional vascular/immune admixture depending on core).
- **Tumor Boundary**: mixed interface niche, typically **Tumor Cells with stromal/immune admixture**.
- **Tumor Core** and **Tumor Core II**: tumor-dominant niches enriched for **Tumor Cell** programs (two related but distinct tumor-centered neighborhoods).

**2) How niches remodel across disease stages (control → patch → plaque → nodular)**

Across stages, we saw a consistent remodeling trajectory in overall tissue organization:

- **Control**: highest representation of **Keratinocyte Enriched** neighborhoods (epithelial/normal-like tissue architecture).
- **Patch / Plaque (intermediate stages)**: expansion of **immune-associated niches** (especially the **macrophage-enriched** and **macrophage–CD8** neighborhoods), consistent with an increasingly inflamed microenvironment.
- **Nodular**: shift toward **tumor-associated niches**, with strong representation of **Tumor Core / Tumor Core II** and more **Tumor Boundary / Tumor Stroma**, indicating progression to more tumor-centered and stromally remodeled architecture.

Biologically, the key learned pattern is a transition from **epithelial-dominant** (control) → **immune-dominant** (patch/plaque) → **tumor-core + boundary/stroma expansion** (nodular), aligning with our goal of understanding how the iTME/niches reorganize with progression.

**3) How KSHV infection distributes across niches and stages**

**A) Infection by niche (strongest signal)**

Using `KSHV_infection_status` (infected if any KSHV gene detected), infection is **highly niche-dependent**:

- **Tumor Core**: **75.76% infected** (33,221 / 43,851)
- **Tumor Core II**: **53.42% infected** (17,490 / 32,739)
- **Tumor Boundary**: **44.41% infected** (20,658 / 46,520)

All non-tumor niches showed **low infection fractions**:

- **Fibroblast Enriched**: 9.85%
- **Tumor Stroma**: 9.67%
- **Macrophage–CD8**: 6.56%
- **Macrophage Enriched II**: 5.98%
- **Pericyte Enriched**: 3.39%
- **Macrophage Enriched**: 2.71%

- **Keratinocyte Enriched**: 1.01%

Interpretation: infection is concentrated in **tumor-centered neighborhoods** (core → core II → boundary), rather than being broadly distributed across immune or keratinocyte niches.

**B) Infection by stage (global trend)**

We also summarized infection as **infected vs uninfected within each `stage`** (stacked percent). The key qualitative pattern is that infection burden tracks progression toward **tumor-associated architecture**—i.e., stages with more **tumor-core / boundary** niche representation show higher overall infected fractions—consistent with the niche-level enrichment above.

If you'd like, I can also pull up the exact stage-by-stage infected percentages (from `data.uns['stage_infection_percents']`) and the exact niche-by-stage percentage table (from `data.uns['stage_niche_percents_rounded']`) in one compact view for reporting.

---

*Associated data:* `query_010_analysis_results.csv`

#### Turn 11 (Query 011)

**User prompt.**

Now let's focus on the immune compartment. For each stage, give me the percentages of each immune cell type (B-cells, CD4 T-cells, CD8 T-cells, Dendritic cells, Macrophages). Which immune populations are most dynamic across the four stages?

---

**Agent activity.**

*Selected tool:* `WriteCodeTool`

▷ Analyzing your question...
▷ Preparing context from memory...
▷ Consulting spatiAlytica Core...
▷ Writing and executing code...
▷ Formatting results...
▷ [OK] Code execution complete (57.0s)
▷ Generating response...
▷ [OK] Response ready (65.3s)
▷ Selected action: Code Execution

---

**spatiAlytica response.**

**Immune compartment percentages by stage**

Two complementary views are shown:

1) **Within-immune (%)** = composition of the immune compartment (these 5 immune types normalized to 100% per stage)
2) **Within-all-cells (%)** = how abundant each immune type is in the full tissue per stage

**1) Percentages within immune cells (immune composition)**

| Stage | B-cells (%) | CD4 T-cells (%) | CD8 T-cells (%) | Dendritic cells (%) | Macrophages (%) |
|---|---|---|---|---|---|
| control | 0 | 0 | 0 | 5.882 | 94.118 |
| patch | 6.293 | 11.807 | 38.020 | 8.574 | 35.306 |
| plaque | 0.238 | 6.435 | 18.502 | 4.699 | 70.126 |
| nodular | 2.517 | 9.458 | 11.937 | 8.020 | 68.067 |

**2) Percentages within all cells (overall tissue abundance)**

| Stage | B-cells (%) | CD4 T-cells (%) | CD8 T-cells (%) | Dendritic cells (%) | Macrophages (%) |
|---|---|---|---|---|---|
| control | 0 | 0 | 0 | 0.262 | 4.196 |
| patch | 2.048 | 3.842 | 12.372 | 2.790 | 11.489 |
| plaque | 0.060 | 1.623 | 4.665 | 1.185 | 17.682 |
| nodular | 0.701 | 2.632 | 3.322 | 2.232 | 18.944 |

---

**Which immune populations are most dynamic across the four stages?**

Dynamics computed from **within-immune (%)** across stages (range = max - min; SD also shown):

| Immune cell type | Range (max - min, %) | SD (%) | Max (%) | Min (%) |
|---|---|---|---|---|
| Macrophages | 58.812 | 20.921 | 94.118 | 35.306 |
| CD8 T-cells | 38.020 | 13.772 | 38.020 | 0 |
| CD4 T-cells | 11.807 | 4.429 | 11.807 | 0 |
| B-cells | 6.293 | 2.526 | 6.293 | 0 |
| Dendritic cells | 3.875 | 1.573 | 8.574 | 4.699 |

**Most dynamic populations: Macrophages** (largest shift) and **CD8 T-cells** (second-largest shift) across the four stages.

---

#### Turn 12 (Query 012)

**User prompt.**

Zoom in on CD8 T-cells specifically: is their percentage trajectory across the four stages monotonic? Show me the per-stage percentages and tell me which way it moves.

---

**Agent activity.**

*Selected tool:* `WriteCodeTool`

- ▷ Analyzing your question...
- ▷ Preparing context from memory...
- ▷ Consulting spatiAlytica Core...
- ▷ Writing and executing code...
- ▷ Formatting results...
- ▷ [OK] Code execution complete (39.0s)
- ▷ Selected action: Code Execution

---

**spatiAlytica response.**

##### CD8 T-cells: is the % trajectory monotonic across stages?

Across the ordered stages **control** → **patch** → **plaque** → **nodular**, the CD8 T-cell percentage trajectory is **not monotonic** under **either** definition (percent of all cells, or percent of immune cells).

---

##### Per-stage CD8 T-cell percentages

**Definition 1: Percent within all cells**

| Stage | CD8 count | Total cells | Percent |
|---|---|---|---|
| control | 0 | 1144 | 0 |
| patch | 1317 | 10645 | 12.372 |
| plaque | 2717 | 58239 | 4.66526 |
| nodular | 7696 | 231661 | 3.3221 |

**Direction:** increases (**control** → **patch**), then decreases (**patch** → **plaque** → **nodular**) → **not monotonic**.

---

**Definition 2: Percent within immune cells**

| Stage | CD8 count | Total immune cells | Percent |
|---|---|---|---|
| control | 0 | 51 | 0 |
| patch | 1317 | 3464 | 38.0196 |
| plaque | 2717 | 14685 | 18.5019 |
| nodular | 7696 | 64473 | 11.9368 |

**Direction:** increases (**control** → **patch**), then decreases (**patch** → **plaque** → **nodular**) → **not monotonic**.

---

*Associated data:* `query_012_analysis_results.csv`

#### Turn 13 (Query 013)

**User prompt.**

Now let's look at the CD8 T-cell program. Run differential expression on the CD8 T-cells comparing patch vs plaque, plaque vs nodular, and patch vs nodular. Give me the top up- and down-regulated genes for each comparison. Focus on host (human) genes; since we're looking at the T-cell program rather than viral content, ignore the KSHV-named genes.

---

**Agent activity.**

*Selected tool:* ***WriteCodeTool***

- ▷ Analyzing your question...
- ▷ Preparing context from memory...
- ▷ Consulting spatiAlytica Core...
- ▷ DE Analysis: CD8 T-cells + patch vs CD8 T-cells + plaque
- ▷ Computing DE: CD8 T-cells + patch vs CD8 T-cells + plaque...
- ▷ Formatting DE results (CD8 T-cells + patch vs CD8 T-cells + plaque)...
- ▷ [OK] DE complete: CD8 T-cells + patch vs CD8 T-cells + plaque (96.3s)
- ▷ DE Analysis: CD8 T-cells + plaque vs CD8 T-cells + nodular
- ▷ Computing DE: CD8 T-cells + plaque vs CD8 T-cells + nodular...
- ▷ Formatting DE results (CD8 T-cells + plaque vs CD8 T-cells + nodular)...
- ▷ [OK] DE complete: CD8 T-cells + plaque vs CD8 T-cells + nodular (105.1s)
- ▷ DE Analysis: CD8 T-cells + patch vs CD8 T-cells + nodular
- ▷ Computing DE: CD8 T-cells + patch vs CD8 T-cells + nodular...
- ▷ Formatting DE results (CD8 T-cells + patch vs CD8 T-cells + nodular)...
- ▷ [OK] DE complete: CD8 T-cells + patch vs CD8 T-cells + nodular (113.7s)
- ▷ Writing and executing code...
- ▷ Formatting results...
- ▷ [OK] Code execution complete (148.5s)
- ▷ Selected action: Code Execution

---

**spatiAlytica response.**

##### CD8 T-cell program — Differential expression (host genes only; KSHV* excluded)

Below are the **top 10 up- and top 10 down-regulated host (human) genes** for each comparison, ranked by **log2FC**.

---

##### Patch vs Plaque (CD8 T-cells)

**Convention:** +log2FC = higher in **patch**; -log2FC = higher in **plaque**

**Top upregulated in Patch (vs Plaque)**

| Gene | log2FC | padj |
|---|---|---|
| CD79A | 2.77726 | 0.00196364 |
| CXCL12 | 2.54981 | 8.75749e-146 |
| CCL19 | 2.42199 | 8.25308e-07 |
| PTGDS | 2.25812 | 1.23134e-32 |
| SOD3 | 1.84968 | 0.0229108 |
| MMP9 | 1.66948 | 7.32964e-05 |
| APOD | 1.59644 | 1.29798e-07 |
| MFAP4 | 1.58383 | 0.0090459 |
| MZB1 | 1.48816 | 0.0964128 |
| IGFBP5 | 1.40441 | 5.04177e-16 |

**Top downregulated in Patch (higher in Plaque)**

| Gene | log2FC | padj |
|---|---|---|
| POSTN | -3.49882 | 8.26947e-68 |
| LUM | -1.62821 | 4.23151e-06 |
| INHBA | -1.50954 | 0.213269 |
| TAGLN | -1.31404 | 0.0966327 |
| MYL9 | -1.15566 | 0.00856522 |
| ADAM12 | -1.0127 | 0.220905 |
| ECSCR | -1.01219 | 0.537217 |
| COL5A2 | -1.00928 | 0.000344857 |
| TM4SF1 | -0.91899 | 0.997118 |
| C5AR1 | -0.916919 | 0.499206 |

---

**Plaque vs Nodular (CD8 T-cells)**

**Convention:** +log2FC = higher in **plaque**; -log2FC = higher in **nodular**

**Top upregulated in Plaque (vs Nodular)**

| Gene | log2FC | padj |
|---|---|---|
| SFRP2 | 1.2856 | 2.55697e-30 |
| CXCL10 | 1.27365 | 5.36312e-10 |
| C1QA | 1.06877 | 6.70658e-05 |
| DSC1 | 1.0478 | 0.980253 |
| MGP | 1.04212 | 1.4154e-16 |
| FRZB | 0.989189 | 0.148184 |
| PLAC9 | 0.893848 | 0.93589 |
| PTGDS | 0.871175 | 0.0528784 |
| NLGN4Y | 0.863012 | 0.819231 |
| MMP27 | 0.815939 | 0.856046 |

**Top downregulated in Plaque (higher in Nodular) — host genes shown**

*(Note: HIV genes were present in the output table; they are not host genes and are omitted here.)*

| Gene | log2FC | padj |
|---|---|---|
| CD79A | -1.70401 | 0.10729 |
| MMP9 | -1.36251 | 3.04623e-06 |
| MZB1 | -1.00886 | 0.165448 |
| CXCL8 | -0.99184 | 0.00106615 |
| DUSP2 | -0.980626 | 0.000722379 |
| SOCS3 | -0.960182 | 1.15192e-13 |
| CXCR4 | -0.902354 | 1.93964e-40 |
| HMGB2 | -0.89956 | 2.57898e-31 |

---

**Patch vs Nodular (CD8 T-cells)**

**Convention:** +log2FC = higher in **patch**; -log2FC = higher in **nodular**

**Top upregulated in Patch (vs Nodular)**

| Gene | log2FC | padj |
|---|---|---|
| PTGDS | 3.1293 | 1.5166e-53 |
| CXCL12 | 2.61216 | 6.43297e-193 |
| CCL19 | 2.21832 | 7.91982e-08 |
| APOD | 2.03247 | 3.37447e-12 |
| TMEM150C | 1.51642 | 0.00138514 |
| SOD3 | 1.46681 | 0.0307745 |
| CLEC10A | 1.44993 | 0.00258412 |
| MGP | 1.40804 | 1.00589e-25 |
| IL33 | 1.40583 | 0.00652955 |
| MFAP4 | 1.36331 | 0.00744432 |

**Top downregulated in Patch (higher in Nodular) — host genes shown**

*(Note: HIV genes were present in the output table; they are not host genes and are omitted here.)*

| Gene | log2FC | padj |
|---|---|---|
| POSTN | -3.38676 | 6.06735e-84 |
| CXCL8 | -1.6924 | 0.0018308 |
| CXCR4 | -1.48893 | 2.37072e-47 |
| TAGLN | -1.48692 | 0.00194074 |
| ADAM12 | -1.31662 | 0.00238856 |
| MYL9 | -1.28064 | 1.39382e-05 |
| TM4SF1 | -1.25453 | 0.707721 |
| CD34 | -1.12473 | 0.000135518 |

---

#### Turn 14 (Query 014)

**User prompt.**

From those stage-pairwise DE signatures, interpret the CD8 T-cell functional shift across KS progression. Are we seeing a migratory program? Cytotoxic engagement? Exhaustion or tissue retention? Walk me through what each stage tells us.

---

**Agent activity.**

*Selected tool:* `RespondWithTextTool`

▷ Analyzing your question...
▷ Preparing context from memory...
▷ Consulting spatiAlytica Core...
▷ Generating response...
▷ [OK] Response ready (29.2s)
▷ Selected action: Response Generation

---

**spatiAlytica response.**

**CD8 T-cell functional shift across KS stages (from the pairwise DE signatures)**

Below I'm interpreting **what changes in the CD8 T-cell-associated expression program** across stages based on the **stage-pairwise DE lists we generated** (patch vs plaque, plaque vs nodular, patch vs nodular; host genes emphasized).

**Patch (earlier) → a "positioning / trafficking / tissue-navigation" state**

Across **patch vs plaque** and **patch vs nodular**, the genes higher in **patch** repeatedly include **CXCL12**, **CCL19**, and endothelial/perivascular-associated transcripts like **RNASE1**, **SOD3**, **MGP**, plus lipid/ECM-adjacent factors like **APOD**.

**Interpretation:** this looks more consistent with a **migratory/positioning program** (cells in or near chemokine-rich, vessel-associated, or lymphoid-organization-like microenvironments) than with a classic cytotoxic effector burst.

- **CCL19** (patch>plaque and patch>nodular) is a strong hint of **lymphoid-like chemokine context** (often linked to organized immune positioning / TLS-like environments).
- **CXCL12** (patch>plaque and patch>nodular) supports **chemokine-guided localization/retention** (CXCL12–CXCR4 axis), but note that the *direction flips* later (see below), suggesting the axis is being rewired across progression.

**What is *not* prominent in these signatures:** a clean increase in canonical cytotoxic genes (e.g., perforin/granzymes) or inhibitory receptors (PDCD1/CTLA4/LAG3/TIGIT). With the genes we have in the top shifts, the evidence for "cytotoxic engagement" or "exhaustion" per se is limited.

**Patch → Plaque: transition toward a matrix/activated-stromal-associated state**

In **patch vs plaque**, plaque-up genes include **POSTN**, **LUM**, **COL5A2/COL6A1**, **TAGLN**, **MYL9**, **INHBA**, **SFRP2**.

**Interpretation:** this is most consistent with CD8 T-cells becoming **embedded in (or tightly associated with) a remodeling ECM / activated stromal program** as lesions mature.

- **POSTN** and **LUM** are classic ECM/stromal remodeling markers.
- **TAGLN/MYL9** imply a more **contractile / actin-cytoskeleton**-linked context.

**Functional read:** this pattern fits better with **tissue retention / physical confinement / altered motility in a fibrotic or tumor-associated stroma** than with increased killing activity. In other words, the CD8 compartment may be shifting from "chemokine-positioned/trafficking" to "stromal-entrained."

**Plaque → Nodular: emergence of inflammatory/stress signaling plus a CXCR4 shift**

In **plaque vs nodular**, nodular-up genes include **CXCR4**, **SOCS3**, **CXCL8**, and T-cell signaling-related genes such as **LCK** and **CD3D**, plus stress/immune-associated genes like **IFITM1**, **ARHGDIB**.

**Interpretation:** this suggests that by **nodular**, the CD8 T-cells are in a **more inflammatory / cytokine-driven environment** with a notable **CXCR4-high** signature.

- **CXCR4** being higher in nodular (it is plaque>nodular negative log2FC, and also nodular>patch in patch vs nodular) supports a move toward **CXCR4-associated retention/positioning** in later disease. That is a plausible "tissue retention" cue.
- **SOCS3** indicates **active cytokine signaling with negative feedback**, often seen in inflammatory contexts.
- **CXCL8** points to a **pro-inflammatory milieu** (and in this dataset it may also reflect microenvironmental admixture; but in any case it tracks with later-stage inflammatory context).

**Patch → Nodular (overall): from chemokine/vascular-adjacent to stromal/inflammatory retention**

The **patch vs nodular** comparison recapitulates both sides:

- Patch-up: **PTGDS**, **CXCL12**, **CCL19**, **APOD**, **RNASE1/SOD3/MGP**.
- Nodular-up: **POSTN**, **CXCL8**, **CXCR4**, plus stromal/cytoskeletal-linked genes (**TAGLN**, **MYL9**, **ADAM12**, **CD34**).

**Interpretation:** overall progression looks like:

1) **Early (patch):** CD8 T-cells associated with **chemokine organization and vessel/immune positioning** (more "migratory/positioning").
2) **Intermediate (plaque):** a strong **ECM remodeling / stromal entanglement** signal emerges (more "tissue architecture-driven retention or exclusion").
3) **Late (nodular):** increased **CXCR4 + inflammatory feedback (SOCS3) + inflammatory chemokine context (CXCL8)**, consistent with **retention in an inflamed, tumor-associated niche**.

**Direct answers to your functional hypotheses**

**Are we seeing a migratory program?**

**Most evidence for migration/positioning is in patch**, driven by **CCL19** and **CXCL12** being higher early, consistent with chemokine-guided localization and immune organization.

**Cytotoxic engagement?**

From the **top changing genes we have in these DE signatures**, there isn't strong direct support for a classic cytotoxic effector program shift (we're not seeing a clear "kill module" dominating the stage differences in the lists we used).

**Exhaustion?**

With these DE signatures alone, **I can't confidently call exhaustion**, because we aren't seeing the canonical inhibitory receptor set in the displayed top shifts. What we *do* see is more consistent with **microenvironmental remodeling + inflammatory signaling**.

**Tissue retention / confinement?**

**Yes—this is the clearest story across progression.** The rise of **ECM/stromal remodeling genes (POSTN, LUM, collagens)** and later **CXCR4** supports a shift toward **retention/confinement in remodeled, tumor-associated tissue** rather than free trafficking.

If you want, I can make this more definitive by generating a focused "function score" panel in CD8 T-cells (e.g., cytotoxicity, interferon response, exhaustion, tissue-residency) using gene sets present in this 308-gene panel, then plotting those scores across stages (patch→plaque→nodular).

---

*Associated data:* `query_014_analysis_results.csv`

#### Turn 15 (Query 015)

**User prompt.**

Now let's look at how the tumor-niche cells themselves change across the late stages. Treat any niche whose biological label suggests it's tumor-related (the tumor or KSHV-tumor niches you named earlier) as one combined tumor-niche group, and run differential expression for cells in that tumor-niche group, comparing nodular vs plaque. Top up- and down-regulated genes please.

---

**Agent activity.**

*Selected tool:* ***DifferentialExpressionTool***

- ▷ Analyzing your question...
- ▷ Preparing context from memory...
- ▷ Consulting spatiAlytica Core...
- ▷ DE Analysis: ['Tumor Stroma', 'Tumor Boundary', 'Tumor Core', 'Tumor Core II'] + nodular vs ['Tumor Stroma', 'Tumor Boundary', 'Tumor Core', 'Tumor Core II'] + plaque
- ▷ Computing DE: ['Tumor Stroma', 'Tumor Boundary', 'Tumor Core', 'Tumor Core II'] + nodular vs ['Tumor Stroma', 'Tumor Boundary', 'Tumor Core', 'Tumor Core II'] + plaque...
- ▷ Formatting DE results (['Tumor Stroma', 'Tumor Boundary', 'Tumor Core', 'Tumor Core II'] + nodular vs ['Tumor Stroma', 'Tumor Boundary', 'Tumor Core', 'Tumor Core II'] + plaque)...
- ▷ [OK] DE complete: ['Tumor Stroma', 'Tumor Boundary', 'Tumor Core', 'Tumor Core II'] + nodular vs ['Tumor Stroma', 'Tumor Boundary', 'Tumor Core', 'Tumor Core II'] + plaque (13.9s)
- ▷ Selected action: Differential Expression

---

**spatiAlytica response.**

**Differential expression in tumor-niche cells — nodular vs plaque**

**Tumor-niche group (combined):** *Tumor Stroma, Tumor Boundary, Tumor Core, Tumor Core II*
**Comparison: nodular (Condition 1)** vs **plaque (Condition 2)**
**Method:** wilcoxon
**Genes analyzed:** 308
**Significant genes:** 279
**log2FC convention:** positive = higher in **nodular**; negative = higher in **plaque**

---

**Top up-regulated in nodular (vs plaque)**

| Gene | log2FC | p-value | padj |
|---|---|---|---|
| KSHV.K2 | 3.3 | 0 | 0 |
| FABP4 | 2.25 | 0 | 0 |
| KSHV.ORF65 | 1.49 | 1.58e-99 | 6.93e-99 |
| KSHV.ORF59 | 1.39 | 1.59e-91 | 6.69e-91 |
| LEPR | 1.36 | 0 | 0 |
| IGFBP3 | 1.17 | 0 | 0 |
| KSHV.K9 | 1.17 | 8.09e-53 | 2.37e-52 |
| LYVE1 | 1.12 | 0 | 0 |
| VWF | 1.11 | 0 | 0 |
| KIT | 1.08 | 0 | 0 |

**Top down-regulated in nodular (i.e., higher in plaque)**

| Gene | log2FC | p-value | padj |
|---|---|---|---|
| SFRP2 | -2.77 | 0 | 0 |
| PTGDS | -2.67 | 0 | 0 |
| FBLN1 | -1.94 | 0 | 0 |
| KRT5 | -1.92 | 5.85e-79 | 2.2e-78 |
| KRT15 | -1.86 | 2.57e-128 | 1.3e-127 |
| PDGFRA | -1.8 | 0 | 0 |
| COL17A1 | -1.65 | 6.98e-97 | 3.03e-96 |
| TACSTD2 | -1.57 | 1.43e-71 | 5.11e-71 |
| INHBA | -1.57 | 0 | 0 |
| LAMB3 | -0.601 | 0.117 | 0.125 |

*Associated data:* `query_015_de_results.csv`

#### Turn 16 (Query 016)

**User prompt.**

Finally, synthesize everything we've found: integrate the niche remodeling across stages, the CD8 T-cell stage trajectory and functional shift, the infection distribution across niches, and the tumor-niche DE between Nodular and Plaque into a coherent model of KS progression. Call out any spatial biomarkers that distinguish the four stages.

**Agent activity.**

*Selected tool:* `RespondWithTextTool`

- ▷ Analyzing your question...
- ▷ Preparing context from memory...
- ▷ Consulting spatiAlytica Core...
- ▷ Generating response...
- ▷ [OK] Response ready (33.7s)
- ▷ Selected action: Response Generation

**spatiAlytica response.**

##### Coherent model of KS progression (integrating niches, CD8 programs, and KSHV infection)

###### 1) Stage progression as spatial ecosystem remodeling

Across stages, KS progression can be viewed as a transition from a **chemokine-positioning / immune–stromal interface** toward a **tumor-structured, vascular/lymphatic, virus-active niche**.

- **Early stages (patch → plaque)**: tissue is dominated by **stromal/ECM-associated neighborhoods** and immune positioning cues. This is the phase where spatial organization looks more like an "inflamed stroma with immune trafficking signals," rather than a fully consolidated tumor architecture.

- **Late stage (nodular)**: niches shift toward **tumor-associated neighborhoods** (Tumor Core / Boundary / Stroma), with **increased vascular/lymphatic programs** and **higher viral activity** in the tumor-niche compartment.

This framing matches our explicit goal: explaining KS progression by linking spatial microenvironments (niches) to immune state, infection burden, and tumor programs.

---

**2) Niche remodeling across stages: from stromal interfaces to tumor-defined compartments**

Using the 10-neighborhood niche labels (k=30), the dataset contains both **tumor-associated niches** (Tumor Core, Tumor Core II, Tumor Boundary, Tumor Stroma) and non-tumor neighborhoods (e.g., Fibroblast Enriched, Keratinocyte Enriched, Pericyte Enriched, Macrophage-enriched variants, Macrophage–CD8).

**Interpretation:**

- **Patch**: enriched for "interface" biology—signals consistent with immune positioning and stromal context.
- **Plaque**: consolidates stromal/tumor interfaces; a transition state where boundary-like organization is prominent.
- **Nodular**: stronger separation into **core vs boundary vs stroma** tumor neighborhoods—i.e., the tissue becomes more spatially compartmentalized into tumor-centric microenvironments.

(We already used these tumor-niche labels to define the tumor-niche DE set, so this remodeling is not hypothetical—it directly informed the late-stage comparison.)

---

**3) CD8 T-cell trajectory: abundance peak + functional reprogramming across stages**

**Composition dynamics:** CD8 T-cells were among the most dynamic immune populations and showed a **non-monotonic trajectory**: **peaking at patch** and then declining toward later stages.

**Functional shift (from stage-pairwise CD8 DE):**

- **Patch (vs plaque and vs nodular)**: enriched for a **positioning/trafficking + tissue-interface** program, including:
  - **CXCL12** (patch>plaque log2FC 2.55; patch>nodular log2FC 2.61)
  - **CCL19** (patch>plaque 2.42; patch>nodular 2.22)
  - plus stromal-associated factors like **PTGDS** (patch>plaque 2.26; patch>nodular 3.13) and **APOD** (patch>nodular 2.03)

  **Model implication:** early lesions support a spatial state where CD8 T-cells are present and "geographically instructed" by chemokine fields / stromal interfaces.
- **Plaque → Nodular transition**: CD8 state shifts away from that early positioning program. In plaque vs nodular, **CXCR4 is higher in nodular** (plaque>nodular log2FC -0.90), consistent with a later-stage program emphasizing **tissue retention / inflammatory signaling context** rather than the early CXCL12/CCL19 positioning signature.

**Putting abundance + program together:** KS seems to move from a stage where CD8 T-cells are relatively more prevalent and chemokine-positioned (patch) to a stage where tumor-associated niches dominate and CD8 programs are altered/less represented overall (nodular).

---

**4) Infection distribution across niches: virus concentrates in specific spatial contexts**

We explicitly computed **infected vs uninfected fractions by niche**. The key biological takeaway is that **infection is not spatially uniform**; it is **niche-structured**.

**Model implication:** as the tissue shifts toward tumor-associated niches, KSHV infection becomes increasingly tied to those microenvironments—consistent with late-stage KS being both **more tumor-architected** and **more virus-active** in the tumor-niche compartment.

---

**5) Tumor-niche DE (nodular vs plaque): late-stage KS is virus-active and vessel/lymphatic-like**

When we restricted to the tumor-associated niches (Tumor Stroma / Boundary / Core / Core II), **nodular vs plaque** showed strong upregulation of:

**Viral genes (nodular higher):**

- **KSHV.K2** (log2FC 3.3)
- **KSHV.ORF65** (1.49), **KSHV.ORF59** (1.39), **KSHV.K9** (1.17)

**Vascular/lymphatic / endothelial-like markers (nodular higher):**

- **VWF** (1.11)
- **LYVE1** (1.12)
- (also present in the broader list: **CDH5**, **KDR**, **CALCRL**, **RAMP2**, **PLVAP**, **PROX1**)

**Metabolic/adipogenic/stromal factors (nodular higher):**

- **FABP4** (2.25)
- **LEPR** (1.36)
- **IGFBP3** (1.17)
- **KIT** (1.08)

**Plaque higher (i.e., down in nodular):**

- **SFRP2** (-2.77)
- **PTGDS** (-2.67)
- **FBLN1** (-1.94)
- basal/epithelial markers **KRT5** (-1.92), **KRT15** (-1.86), **COL17A1** (-1.65), **TACSTD2** (-1.57), plus **PDGFRA** (-1.8) and **INHBA** (-1.57)

**Model implication:** late-stage nodular KS within tumor niches is characterized by a **virus-high, vascular/lymphatic-like, metabolically distinct** program, whereas plaque retains more **ECM/epithelial-associated** features.

---

**Spatial biomarkers that distinguish the four stages (practical readouts)**

Below are "spatial biomarkers" in the sense of features you can map on tissue to separate stages via expression fields and/or enrichment in specific neighborhoods.

**Patch**

- **CXCL12-high** and **CCL19-high** spatial fields (also seen as top CD8-associated differences vs later stages).
- CD8 T-cells are relatively most prominent at this stage (composition peak), consistent with early immune positioning.

**Plaque**

- A transition state with stronger **ECM/interface** biology.
- Tumor-niche plaque signal (relative to nodular): **SFRP2**, **PTGDS**, **FBLN1**, **PDGFRA**, **INHBA**, and epithelial-associated **KRT5/KRT15/COL17A1/TACSTD2**.

**Nodular**

- **KSHV viral gene hotspots** in tumor-associated niches: **KSHV.K2**, **KSHV.ORF65**, **KSHV.ORF59**, **KSHV.K9**.
- **Vascular/lymphatic program** spatially concentrated in tumor niches: **VWF**, **LYVE1** (and in the extended DE list: **CDH5, KDR, PLVAP, PROX1**).
- **FABP4/LEPR** as a metabolic/stromal axis associated with late-stage tumor-niche remodeling.
- CD8 program shifts away from the early CXCL12/CCL19 signature; **CXCR4** becomes relatively higher in nodular vs plaque within CD8s.

**Macular**

We did not explicitly run a macular-specific DE block in the results summarized above, so I'm not going to invent a marker list. Conceptually, macular would be expected to sit between "baseline skin" and the patch program; to call spatial biomarkers rigorously for macular, we'd want a targeted contrast (e.g., macular vs patch in tumor-adjacent niches and/or within immune subsets).

---

**One-sentence synthesis**

KS progression in this dataset is best explained as **stage-dependent spatial niche remodeling** that reorganizes immune positioning (early CD8-rich, CXCL12/CCL19-associated patch lesions) into **tumor-compartmentalized, infection-structured microenvironments** where **nodular tumor niches become virus-high and vascular/lymphatic-like** (KSHV.K2 with LYVE1/VWF and FABP4/LEPR), while plaque retains comparatively more ECM/epithelial-associated programs (SFRP2/PTGDS/FBLN1/KRT5/KRT15).

---

*Associated data:* `query_016_analysis_results.csv`
